# A single day of natural daylight exposure and physical activity does not conclusively change circadian phase: A Bayesian crossover experiment

**DOI:** 10.64898/2026.11.03.763886

**Authors:** Ann-Sophie Loock, Marta Milanesi, Fabian Schwendinger, Corrado Garbazza, Arno Schmidt-Trucksäss, Undine Lang, Christine Blume

**Author notes:** **Corresponding author:** Dr. habil. Christine Blume Wilhelm-Klein-Str. 27, 4002 Basel Switzerland.

## Abstract

Among the environmental signals that shape human physiology, the daily light-dark cycle is the primary zeitgeber of the human circadian system. Physical activity may provide an additional non-photic time cue and frequently co-occurs with daylight in everyday life. However, their independent and combined effects on circadian phase have rarely been disentangled under naturalistic daylight conditions. In a counterbalanced mixed-design experiment, 41 healthy adults (24 women; mean age = 23.7 years, *SD* = 4.3) participated in up to three 40-h visits on fixed individual sleep-wake schedules. Each visit comprised an 8-h exposure beginning 1.5 h after wake to natural daylight, daylight attenuated by amber-tinted glasses, or indoor electric light (within subjects; mean photopic illuminances: 1196, 460, and 75 lx), with otherwise dim-light conditions. Participants hiked for approximately 4 h or remained sedentary (between subjects; activity indexed by median heart rate reserve [HRR]; group means: 32.7% and 2.7%). Bayesian hierarchical models used individual photopic illuminance (ActTrust 2) and median HRR to predict change in dim-light melatonin onset (DLMO; primary outcome) and exploratory outcomes (daytime and evening sleepiness, mood, and wellbeing, daytime perceived exertion, and evening psychomotor vigilance). Per standard-deviation increase in log-transformed illuminance and median HRR, estimated DLMO changes were −4.22 min (95% HDI [−10.93, 2.22]; *BF*₁₀ = 0.70) and −6.48 min (95% HDI [−13.79, 1.27], *BF*₁₀ = 1.56), respectively. Evidence remained inconclusive for both predictors and their interaction. Exploratory findings were largely inconclusive, practically negligible, or favoured the absence of effects. The circadian effects of a single day of natural daylight exposure, with or without physical activity, therefore remain unclear. Resolving this question may require larger samples and brighter, repeated, or habitual exposure timed to phases of greater circadian light sensitivity.

---

The average adult in industrialised societies spends approximately 90% of their time indoors^1^, where daytime illuminance is typically substantially lower than outdoors^2,3^. At the same time, electrical lighting in the evening and at night has become ubiquitous. Hence, the modern light environment offers far less day-night contrast than the natural light-dark cycle^4,5^. Low daytime light exposure may also increase the circadian system’s sensitivity to subsequent light, such that after a period of dim prior light, nocturnal light suppressed melatonin more strongly^6^ and induced larger phase shifts^7^. Large-scale observational data link this pattern to health outcomes. In the UK Biobank, darker days and brighter nights were associated with psychiatric disorders^8^ and higher all-cause mortality^9^. Understanding how daytime light shapes circadian physiology is therefore of growing importance.

The human circadian system has an endogenous period averaging slightly longer than 24 h^10^ and therefore requires regular adjustment to the solar day, primarily through light^11^. Light information reaches the circadian pacemaker in the suprachiasmatic nuclei via the retinohypothalamic tract, principally through intrinsically photosensitive retinal ganglion cells (ipRGCs) that express the photopigment melanopsin^12–14^. Melanopsin has peak sensitivity near 480 nm^15^, consistent with the short-wavelength sensitivity of melatonin^16,17^. Accordingly, melanopic illuminance predicts melatonin suppression^18^ and circadian phase resetting^19^, whereas calibrated blue-yellow colour changes at constant melanopsin excitation yielded no conclusive differences in either response^20^. In natural environments, melanopic and photopic illuminance are nevertheless strongly correlated^3^. Light generally advances circadian phase in the biological morning, delays it in the biological evening and early night, and produces smaller responses around the middle of the biological day^21,22^. Response magnitude also depends on intensity, spectral composition and duration^23–25^. Most of this evidence, however, stems from electric light applied in the morning, evening, or night under laboratory conditions. Midday exposure, when natural illuminance peaks, has thus received comparatively little attention.

Observational studies associate more time outdoors with earlier sleep timing^26,27^. Greater daylight exposure was also associated with earlier dim-light melatonin onset (DLMO) in an ambulatory study that modelled individual intrinsic circadian period^28^. Experimentally, a week of summer camping advanced DLMO by approximately 2 h^4^, while a single weekend achieved about 69% of that shift^5^. Camping, however, alters the entire light-dark cycle and may also change sleep schedules and physical activity, preventing isolation of daytime light effects. Other experiments have manipulated daylight access through windows over several days^29,30^ or used electric light in controlled laboratory approaches. For instance, compared with a conventional LED matched for photopic illuminance and correlated colour temperature, a daylight LED improved alertness, mood, and visual comfort, while melatonin profiles measured under the respective light conditions did not differ^31^. Such approaches do not fully reproduce natural daylight, as its intensity and spectrum change continuously with solar elevation and weather^32^, and outdoor exposure is inseparable from other features of being outside. Consequently, whether a single, defined daytime exposure to natural daylight changes circadian phase remains unknown.

Physical activity may additionally influence circadian phase through non-photic pathways, although its effects are generally considered weaker than those of light^33^. Similar to light, exercise-induced phase shifts depend on timing. Exercise around 7:00 and between 13:00 and 16:00 advanced the melatonin rhythm, whereas exercise between 19:00 and 22:00 delayed it^34^; responses varied with chronotype, particularly for evening exercise^35^. Nocturnal exercise also facilitated phase delays under very dim light^36^. Although daylight and physical activity commonly co-occur outdoors, experimental evidence on their combined effect is limited. Bright light delivered between 23:00 and 02:00 combined with vigorous exercise did not produce significantly larger delays than light alone, although the study could only detect differences of about 2-2.5 h^37^. In contrast, in a pilot study with six participants, late-evening bright light followed by early-morning exercise produced larger phase delays than late-evening exercise alone^38^. Notably, all of these studies used electrical lighting at night. Whether physical activity modulates the circadian response to natural daylight has, to our knowledge, not been examined.

The present study examined whether daytime light exposure and physical activity, individually and in combination, change circadian phase within a single day. In a mixed design, participants completed a counterbalanced crossover of three 8-h light conditions beginning 1.5 h after wake (natural daylight, daylight attenuated by amber-tinted glasses, and indoor electrical lighting). Physical activity was varied between participants by allocating them to an active protocol with approximately 4 h of hiking or an inactive protocol. Individually measured photopic illuminance and physical activity, indexed by heart rate reserve, were modelled as continuous predictors, thereby retaining the variation in achieved exposure. We hypothesised that higher daytime light exposure would produce a larger change in DLMO and that physical activity would interact synergistically with light. Exploratory secondary outcomes comprised daytime and evening sleepiness, mood, and wellbeing, daytime perceived exertion, and evening psychomotor vigilance.

## Method

### Ethics and Registration

The study was approved by the Ethics Committee of Northwestern and Central Switzerland (EKNZ; approval number 2022-01140) and conducted in accordance with the Declaration of Helsinki and Swiss law. It was registered as a clinical trial (ClinicalTrials.gov: NCT05513547) and pre-registered on the Open Science Framework (OSF; https://doi.org/10.17605/OSF.IO/7B29Q). All participants provided written informed consent.

### Participants

Healthy adults aged 18-35 years were recruited through the University of Basel participant portal (www.unimarkt.unibas.ch) and social media. The age range was restricted to minimise interindividual variability in pre-receptoral light filtering^39,40^. Eligibility required a body mass index of 18.5-24.9 kg/m² (WHO normal-weight range) and a moderate level of habitual physical activity, operationally defined as regular sport or exercise without evidence of a sedentary lifestyle or competitive athletic training. Habitual activity was assessed with the short form of the International Physical Activity Questionnaire^41^ (IPAQ-SF) and self-reported sport frequency, which informed an overall eligibility judgement rather than a fixed metabolic equivalent time per week cut-off. Cardiorespiratory fitness was additionally assessed with a ramp test on a cycle ergometer, both to ensure medical safety for the outdoor activity protocol and to determine individual maximal heart rate. Eligibility further required an approved medical examination by the study physician, and written informed consent. Exclusion criteria comprised pregnancy; chronic medical, psychiatric, sleep, or circadian disorders; use of medication affecting sleep or circadian physiology; recreational drug use (urine multi-drug panel; nal von minden); extreme chronotype (mid-sleep on free days ≤ 2:00 or ≥ 7:00, Munich Chronotype Questionnaire^27^ [MCTQ]); shift work within the preceding three months; and transmeridian travel across more than two time zones within the preceding month. Non-adherence to the prescribed sleep-wake schedule during ambulatory periods (> 2 deviations per study week, or any deviation on either of the two days preceding an experimental visit) likewise led to exclusion. Screening questionnaires, including a daily sleep-wake diary with evening and morning parts, were administered in German via REDCap^42,43^ hosted on the University of Basel’s sciCORE. Full inclusion and exclusion criteria with instruments and cut-offs are provided in Supplementary Table S1. Sample size was planned with a pre-registered Sequential Bayes Factor (*BF*₁₀) design, in which recruitment was to continue until the primary analysis reached *BF*₁₀ ≥ 10 or the project’s financial or temporal resource limits were reached (anticipated maximum: 48 participants). Because recruitment proceeded more slowly than anticipated and interim analyses did not meet the stopping criterion, data collection continued until the latest feasible end point within the project period (September 2025). Thus, the final sample size was determined by this temporal constraint. Participants received CHF 1,000 for completing the full protocol, with partial compensation for early withdrawal.

### Design

A mixed design was used, combining a within-participant manipulation of light exposure (three conditions; see below) with a between-participant manipulation of physical activity (active vs. inactive; Figure 1A). Each participant completed three approximately 40-h experimental visits, spaced one week apart and scheduled on the same weekday to minimise carry-over and scheduling effects. Within each activity group, the order of light conditions was counterbalanced by blocked randomisation (*blockrand* R-package^44^), stratified by sex and age. Participants were pseudo-randomly allocated to the active or inactive group; where possible, a second participant was matched to the opposite activity group within the same testing slot to allow parallel data collection. Data were collected between May 2023 and September 2025, excluding the winter months (approximately November-March), when outdoor conditions around Basel are unfavourable.

**Figure 1.**
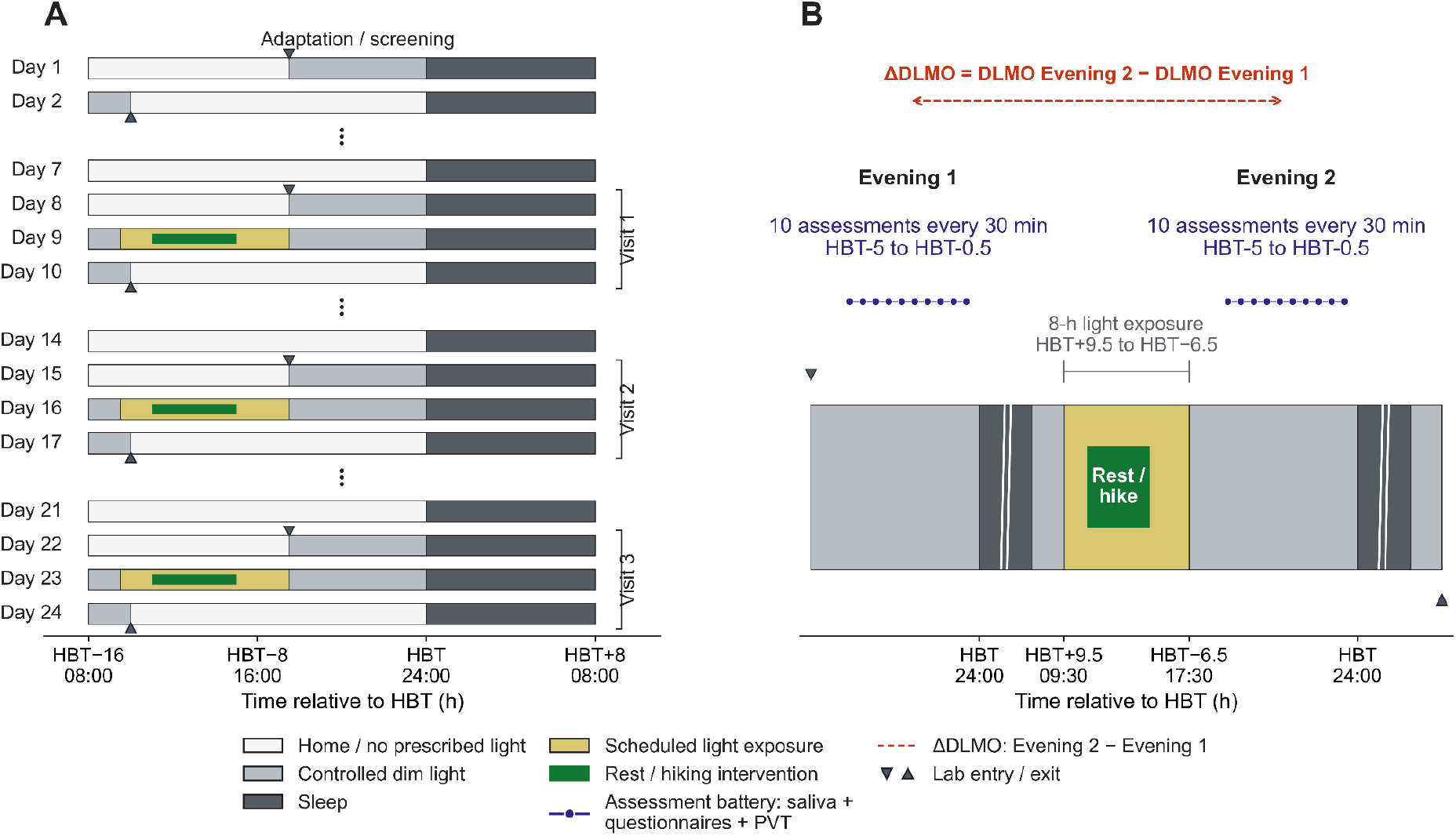
Study Protocol and Timeline of an Experimental Visit. *Note.* (A) After an adaptation night (Day 1), participants followed a fixed sleep-wake schedule at home and completed three experimental visits at weekly intervals (shown on Days 8-10, 15-17, and 22-24). The order of the daylight, glasses, and laboratory conditions was counterbalanced within each activity group. (B) Timeline of one experimental visit. DLMO was assessed on the evening before (Evening-1) and after (Evening-2) the 8-h light-exposure window, during which the active group walked approximately 4 h and the inactive group remained sedentary. Clock times refer to a habitual bedtime (HBT) of 24:00 and were adapted to each participant’s HBT. DLMO = dim-light melatonin onset; PVT = psychomotor vigilance task.

### Light Exposure

Three light-exposure conditions were implemented during an 8-h daytime window (for its placement, see *Procedure*). In the daylight condition, participants spent the window outdoors in natural light at a fixed location in the greater Basel area (Nunningerberghütte, Nunningen, Switzerland; 47.381° N, 7.624° E; 885 m a.s.l.), including the approximately 1-h drive to and from this location. In the glasses condition, participants were outdoors at the same location while wearing amber-tinted filter glasses (Uvex, Honeywell, Charlotte, NC, USA) that specifically strongly attenuate short-wavelength content, so that the glasses condition provided an intermediate level on the continuous light predictor (Supplementary Method S4). In the laboratory condition, participants remained indoors under controlled artificial lighting (a fluorescent ceiling-lamp panel; Philips Lighting, Eindhoven, The Netherlands) for the window; illuminance at eye level was a median of 51 lx (range 44-85; characterised at seven typical participant positions and measured with a calibrated spectroradiometer, JETI spectraval 1501, JETI Technische Instrumente GmbH, Jena, Germany) at a correlated colour temperature of approximately 3100 K (warm-white). In all conditions, sunglasses and headwear shading eyes (e.g., caps) were not permitted during the exposure window. Outside the exposure window, laboratory background lighting was held at dim-light levels (< 20 lx; *M* = 10.55 lx at eye level) throughout, including both evening test batteries (Supplementary Method S3, Figure S2, and Table S2).

Light exposure during the window was recorded continuously at 1-min resolution with a forehead-mounted ActTrust 2 (Condor Instruments, São Paulo, Brazil; hardware v1.3, firmware v1.6) held in a custom 3D-printed housing and oriented in the direction of gaze. Analyses used photopic illuminance (device’s LIGHT channel). Because the sensor was worn outside the filter in the glasses condition, values in that condition were multiplied by the estimated LIGHT-channel transmittance of the filter (40.38%) to approximate the light reaching the eye; no correction was applied in the daylight or laboratory conditions. The correction was applied once, at the derivation stage of the pipeline. The derivation of this factor is provided in Supplementary Method S4 (Figure S3).

Recordings were screened for technical failure before analysis, and two visit-level values were set to missing. One returned only zero values across the window (device off or technical failure). The other, a laboratory visit, showed implausibly high photopic illuminance (median 925 lx, maximum 13,981 lx) for the controlled setting, with 448 of 480 min exceeding a 100-lx plausibility threshold, indicating a sensor or orientation artefact rather than a substantive outlier (Method S4). Because minute-level illuminance is strongly right-skewed and intermittently dominated by brief extremes (e.g., direct solar glare or transient occlusion), each visit was summarised by the median rather than the mean; the transformation and modelling of this value are described in Data Analysis.

### Procedure

#### Screening

Prior to participation, candidates completed psychometric screening questionnaires in REDCap^42,43^ and a cardiopulmonary exercise test (ramp test on a bicycle ergometer) at the Department of Sport, Exercise and Health, University of Basel. The test continued to volitional exhaustion, with continuous breath-by-breath recording of gas exchange and heart rate, using a procedure similar to that described by Wagner et al^45^. Candidates subsequently completed an adaptation night in the laboratory, followed by a medical examination by a study physician. The adaptation night was scheduled to self-selected habitual bed and wake times (8-h sleep opportunity) and recorded with full polysomnography (PSG) to familiarise candidates with the laboratory setting and screen for sleep disorders.

#### Ambulatory Phase

Across the 24-day protocol, participants maintained a fixed 16:8 wake-sleep schedule, self-selecting a habitual bedtime (HBT, lights-off) and wake time (8 h apart) and thereafter adhering to both within ±30 min. Such stabilisation protocols are used to regularise sleep-wake timing and support stable circadian entrainment before laboratory assessment^46,47^. Throughout participation, participants abstained from caffeine, alcohol, and recreational drugs, and refrained from daytime naps. Adherence was verified with a sleep log^48^ and wrist actigraphy (ActiGraph CentrePoint Insight Watch, ActiGraph LLC, Pensacola, FL, USA). On each visit day (visits started in the afternoon), they additionally avoided intensive exercise and wore a collar-mounted ActTrust 2 at chest height from waking until arrival at the laboratory to record prior light history summarised as mean photopic illuminance, cumulative exposure (lux-minutes), and time above 250 lx and 1,000 lx for a check of comparable exposure across conditions (Method S4, Results S1, Table S4). Sunglass use was not restricted or recorded on these days.

#### Experimental Visits

Each visit spanned approximately 40 h across two nights in a light-, temperature-(22°C), and sound-controlled laboratory at the Centre for Chronobiology, University Psychiatric Clinics Basel, without external time cues. Participants had no access to phones or computers and no knowledge of clock time, except for a “social hour” at the start of the light-exposure window (1.5 h after wake), during which participants were allowed to use their smartphones. In the outdoor conditions, this coincided with the drive to the exposure site. Reading material, card games, and audio without display light were permitted. Meals matching individual daily metabolic needs, estimated from Harris-Benedict equation^49^ multiplied by a physical activity level of 1.4 (inactive) or 1.8 (active), were provided, with water available ad libitum.

Each visit was structured as follows (all times referenced to individual HBT; Figure 1B). On arrival (6.5 h before HBT), participants completed a urine multi-drug panel (nal von minden, Rev03.01 / 0702al000005; testing for amphetamines, benzodiazepines, cocaine, morphine/opiate, methadone, and tetrahydrocannabinol) while sleep-wake time adherence over the preceding days was checked. PSG preparation began 6 h before HBT. From 5 h until 0.5 h before HBT, participants completed the Evening-1 test battery at 30-min intervals (10 sessions; PsychoPy v2022.2.5^50^), comprising a saliva sample, subjective ratings (sleepiness, mood, wellbeing), and a psychomotor vigilance task (PVT). Between assessments they remained sedentary. Immediately after the final session and approximately 20 min before HBT, a 3-min resting-state EEG (eyes open) was recorded, and participants completed the evening part of the sleep log before an 8-h sleep opportunity in darkness (PSG-monitored). After waking, participants provided a saliva sample 5 min after wake together with subjective ratings (sleepiness, mood, wellbeing), underwent a second 3-min resting-state EEG 10 min after wake (from which resting heart rate was derived, see *Measures*), completed the morning part of the sleep log (including the sleep quality scale of the self-rating questionnaire for sleep and awakening quality [SSA]^48^) and provided a further saliva sample 35 min (with the same subjective ratings) after wake.

Following breakfast, participants donned the forehead-mounted light sensor approximately 1 h after wake, and the 8-h light-exposure window began approximately 1.5 h after wake (this included travel times in the outdoor conditions). In the active group, physical activity began approximately 3 h after wake and comprised approximately 4 h of walking in four bouts separated by short breaks - outdoors, a fixed hiking route in the greater Basel area; indoors, treadmill walking (Pluto Med, h/p/cosmos, Nussdorf-Traunstein, Germany) programmed to approximate the route (Method S2, Figure S1) - whereas inactive participants remained sedentary. Subjective ratings of sleepiness, mood, wellbeing, and perceived exertion were provided at 30-min intervals on a tablet (Amazon Fire HD 10 Kids Pro; 12 assessments) beginning immediately before the activity period (or the corresponding sedentary period) and extending beyond its end to accommodate slower walking paces and to capture immediate after-effects of activity. The exposure window ended approximately 9.5 h after wake, after which participants returned to the laboratory, and the second evening (Evening-2), night, and morning followed the same procedure as the first.

Between visits, participants resumed their normal daily life while maintaining the sleep-wake, caffeine, and drug restrictions. Data acquisition ended on the morning of day 24. A few minor scheduling deviations occurred, mostly owing to travel or weather conditions. In five visits, the light-exposure period started 5-15 min later than scheduled. Its end, and thus the return to the laboratory, was later than planned in five visits (four by 4–18 min, one by 45 min) and earlier in two (by 10 and 69 min). PSG preparation was adjusted accordingly, whereas all subsequent timings remained anchored to HBT. Because the exact visit end depended on how long participants took to shower and eat after the final morning saliva sample, total duration varied slightly.

### Measures

#### Dim-Light Melatonin Onset

Saliva samples (≥ 1 mL) were collected using Salivettes (Sarstedt AG, Sevelen, Switzerland), immediately centrifuged at 3,000 rpm for 3 min, and frozen at −28 °C until assay. Samples were collected every 30 min from 5 h before until 30 min before HBT on each of the two evenings of a visit (10 samples per evening), together with two morning samples 5 and 35 min after wake time, yielding 24 samples per visit. Salivary melatonin was assayed by radioimmunoassay (Direct Saliva Melatonin RIA, RK-DSM2; NovoLytiX GmbH, Witterswil, Switzerland; analytical sensitivity of 0.2 pg/mL, limit of quantification of 0.9 pg/mL, intra- and inter-assay coefficients of variation of 7.9% and 9.8%).

Dim-light melatonin onset (DLMO) was determined from each evening profile with a piecewise linear-parabolic fit (Hockey-Stick software v2.5^51^), using a default upper threshold of 5 pg/mL, adjusted to individual baseline values where necessary. In one case, a single baseline point was added to improve approximation of the baseline part, in line with developer recommendations. Two raters (A.L. and C.B.) determined DLMO independently and blinded to condition; discrepancies were resolved in discussion with a third expert rater.

#### Physical Activity

Relative physical activity intensity was quantified as percentage of heart rate reserve (%HRR). Heart rate (HR) during the activity period, or the corresponding sedentary period, was recorded continuously with a Polar H10 chest-strap sensor (Polar Electro Oy, Kempele, Finland) connected to the Polar Beat smartphone application. HR values were averaged within consecutive 5-min bins, and missing bin-level values were imputed separately for the active and inactive groups using predictive mean matching (*mice* package^52^). Resting heart rate (HRrest) was derived from inter-beat intervals detected in the ECG channel of the morning resting-state EEG recordings. The resulting beat-per-minute values were interpolated, smoothed, and screened for artefacts (Method S5). Maximal heart rate (HRmax) was determined during the screening cardiopulmonary exercise test. For each 5-min bin, %HRR was calculated as [(HR5min − HRrest) / (HRmax − HRrest)] × 100, where HR5min denotes the mean HR within that bin. Values below zero were set to zero. The median %HRR across bins within each activity or corresponding sedentary period was used as the continuous physical-activity predictor in all models.

#### Subjective Ratings

Subjective sleepiness, mood, and wellbeing were rated at 30-min intervals during both evening test batteries (ten assessments per evening) and, together with perceived exertion, during the activity period (twelve assessments). Sleepiness was assessed with the German version of the Karolinska Sleepiness Scale^53^ (KSS), a single-item 9-point scale from 1 (very alert) to 9 (very sleepy). Momentary mood and physical wellbeing were each rated on a 5-point Likert scale from 1 (very bad) to 5 (very good). Perceived exertion was rated on a modified Borg scale from 0 (rest) to 10 (maximal)^54^.

#### Psychomotor Vigilance

Sustained attention was measured with a modified 10-min auditory version of the Psychomotor Vigilance Task^55,56^ (PVT) administered in PsychoPy (v2022.2.5^50^). Auditory stimuli (70 dB) were presented at inter-stimulus intervals jittered between 2 and 9 s. Responses faster than 100 ms were classified as false starts and removed from all reaction-time (RT) summaries, whereas responses of 500 ms or longer were retained and additionally counted as lapses^56^. Sessions with documented technical problems were excluded. Per session, we analysed median RT and mean response speed (1/RT). The pre-registered response speed of the slowest 10% of responses was also analysed and is reported in the Supplementary Material.

## Data Analysis

### Outcomes

The primary outcome was the change in DLMO, defined within each visit as DLMO on Evening-2 minus DLMO on Evening-1 and expressed in minutes, with negative values indicating an earlier DLMO on Evening-2. All secondary outcomes were pre-registered as exploratory. These comprised sleepiness, mood, overall wellbeing, and psychomotor vigilance (median reaction time and mean response speed, 1/RT) assessed repeatedly on Evening-2, as well as sleepiness, mood, wellbeing, and perceived exertion assessed repeatedly during the activity period. Visual comfort was recorded during the exposure window but lies beyond the scope of this report. Because the secondary outcomes were exploratory and not corrected for multiplicity, they are interpreted as estimates with uncertainty rather than as confirmatory hypothesis tests.

### Predictors

Every model included continuous measures of light exposure and physical activity and their interaction. Light exposure was defined as the median photopic illuminance across the 8-h exposure window, transformed as log₁₀(*x* + 1) and *z*-standardised. The transformation accommodated the strongly right-skewed exposure distribution and the established nonlinear intensity-response relationship of the human circadian system^24^. Physical activity was defined as the median heart rate reserve (HRR) across the activity period, a shorter period nested within the light-exposure window, likewise *z*-standardised.

### Model Structure

Outcomes were analysed with Bayesian hierarchical models using Gaussian likelihoods. The model for the change in DLMO included a participant-level random intercept.

Repeated evening and daytime outcomes were modelled as time courses, with assessment centred at the midpoint of each series. The evening models examined whether the post-exposure trajectory depended on light exposure or physical activity and therefore included interaction terms of assessment with light and activity, with random intercepts and assessment slopes at both the participant and visit level. The daytime models, in contrast, addressed overall associations across the activity period and therefore included assessment as an additive fixed effect, with random intercepts at both levels and random assessment slopes at the participant level. Exact model equations and procedures are provided in Supplementary Method S6 (Equations S3-S5). The posterior predictive check for the primary model is shown in Supplementary Figure S9.

### Priors

Weakly regularising priors were scaled to the standard deviation (*SD*) of the respective outcome in each model’s analysis sample. Intercepts were assigned Normal(*ȳ*, 2 × *SD*) priors, regression coefficients Normal(0, *SD*/3) priors, and residual and random-effect standard deviations half-Normal priors scaled to the outcome *SD* (Method S6, Table S3).

### Estimation and Convergence

Models were fitted in PyMC using the No-U-Turn sampler (NUTS). Four chains ran 10,000 tuning iterations and 6,000 posterior draws per chain, with a target acceptance rate of 0.95 and a maximum tree depth of 15. Sampling quality was assessed with rank-normalised split-*R^*, effective sample size (ESS), trace plots, and divergent transitions. Models were refitted with a higher target acceptance probability when a divergent transition occurred, maximum *R^* exceeded 1.02, or minimum ESS was below 1,000. All models retained for inference had *R^* ≤ 1.01, minimum ESS > 1,000, and no divergent transitions. Model-specific sample sizes and diagnostics are reported in Supplementary Table S16.

### Inference and Reporting

Posterior distributions were summarised by their mean and 95% highest density interval (HDI). Because both predictors were *z*-standardised, the light exposure and physical activity coefficients represent the expected outcome difference per 1-*SD* increase in the respective predictor at the mean of the other, and the interaction coefficient represents the change in the light exposure coefficient per 1-*SD* increase in physical activity. Each effect was additionally characterised by its probability of direction^57^ (*pd*), the percentage of the posterior within an outcome-specific region of practical equivalence^58^ (ROPE; Table S3), and a Bayes factor (*BF*₁₀) computed with the Savage-Dickey density ratio against a point null at zero under the coefficient’s prior^59^. *BF*₁₀ of 1–3, 3–10, 10–30, and > 30 were described as anecdotal, moderate, strong, and very strong evidence for a non-zero effect, and reciprocal values as evidence for the null^60,61^.

These indices were interpreted jointly. An effect was considered *evidenced* when its 95% HDI excludes zero, most of its posterior lay outside the ROPE, and *BF*₁₀ ≥ 3; *practically negligible* when the posterior concentrated inside the ROPE, regardless of whether the HDI excluded zero; and *inconclusive* when 1/3 < *BF*₁₀ < 3, meaning the data did not discriminate between the presence and absence of an effect. Inconclusive results are not interpreted as evidence against an effect. Only *BF*₁₀ ≤ 1/3 with the posterior inside the ROPE was interpreted as *evidence against* a practically relevant effect. Derived contrasts comparing high (+1 *SD*) with low (−1 *SD*) predictor values are reported with posterior means, 95% HDIs, *pd*, and, where applicable, ROPE percentages, but without Bayes factors (Method S6).

Candidate fixed-effect specifications (e.g., for the change in DLMO: null, light only, activity only, additive, and full interaction) were compared with Pareto-smoothed importance-sampling leave-one-out cross-validation^62^ (PSIS-LOO). A model was considered predictively superior only when its difference in expected log pointwise predictive density (ΔELPD) exceeded approximately twice its standard error. ΔELPD is reported as the ELPD of the best-ranked model minus that of the comparison model. Comparisons with any Pareto-*k^* > 0.7 were deemed unreliable (Method S6, Results S7, Table S15).

### Analysis Samples and Missing Data

Complete-case deletion was applied separately for each model: an observation was excluded only when that model’s outcome or one of its predictors was missing. Consequently, analysis samples differ across outcomes. The numbers of contributing observations, visits, and participants are reported in Supplementary Table S16.

### Sensitivity Analyses

Sensitivity analyses of the primary outcome addressed the likelihood, priors, and operationalisation of light exposure. We refitted the primary model with (a) a Student’s *t*-likelihood; (b) wider coefficient priors, Normal(0, 0.5 × *SD*); (c) cumulative rather than median photopic exposure; (d) for laboratory visits with raw illuminance above 100 lx in at least 25% of recorded minutes, replacing minute-level values above 200 lx with 85 lx before recalculating the visit median; (e) adjustment for age and sex; and (f) light condition and physical activity group entered categorically rather than continuously. Full specifications and results are reported in Supplementary Method S7 and Results S2 (Tables S5-S7).

Because the activity groups already differed in psychomotor vigilance before the manipulation (Table S4), Evening-2 vigilance outcomes were additionally analysed using baseline adjustment. The reported baseline-adjusted model (M1) included the standardised Evening-1 visit-level value as a single covariate^63^. Two further models separated baseline performance and physical activity into between-and within-participant components^64^ (Method S8, Results S5, Tables S11-S12).

### Additional Exploratory Analyses

As a not pre-registered exploratory analysis, evening melatonin secretion was quantified as the area under the concentration-time curve (AUC), calculated using the trapezoidal rule over the evening sampling window (HBT −5 h to HBT −0.5 h). Visits were retained when at least six valid samples and both the first and last samples were available. The ratio of Evening-2 to Evening-1 AUC was log₁₀-transformed, with positive values indicating greater melatonin secretion on the post-exposure evening. This outcome was analysed with the same fixed effects, random effects, priors, and sampling settings as the primary DLMO model. Results are reported in Supplementary Results S3 and Table S8.

## Software

Data preprocessing and management were conducted in R 4.5.2^65^ with *data.table* and *ggplot2* packages^66,67^. Bayesian models were implemented in Python 3.11.9^68^ with PyMC 5.27.1^69^ and ArviZ 0.23.4^70^. During manuscript preparation, the authors used ChatGPT (OpenAI) and Claude (Anthropic) for coding assistance (troubleshooting and plotting) and language polishing to improve clarity, coherence, and wording. All outputs were critically reviewed and edited by the authors, who take full responsibility for the content of the publication.

### Deviations from Pre-Registration

The study departed from the pre-registered plan (OSF: https://doi.org/10.17605/OSF.IO/7B29Q) in four respects. First, although both photopic and melanopic-weighted exposure were pre-registered as light characteristics to be varied and monitored, the ActTrust 2 does not provide a validated melanopic channel, and a melanopic-weighted value could only be approximated from device-specific correction factors. Photopic illuminance was therefore retained as the primary exposure metric; devices that derive melanopic equivalent daylight illuminance directly (e.g., ActLumus) were not available at study onset. Second, whereas the pre-registration framed the confirmatory hypotheses as categorical contrasts between the three light-exposure protocols and between the two activity groups, light and physical activity were instead modelled as continuous predictors together with their interaction, thereby retaining the substantial within-group variation in achieved exposure and exertion; the categorical condition contrast is reported as a sensitivity analysis. Third, salivary melatonin was assayed by radioimmunoassay rather than the pre-registered enzyme-linked immunosorbent assay. Fourth, mean illuminance of spot measurements under dim-light conditions was 10.55 lx, slightly exceeding the pre-registered ceiling of 10 lx, and with a maximum of 16.7 lx near the background light source (all measurements are reported in Supplementary Table S2). The remaining aspects of the design and analysis followed the pre-registration.

## Results

### Participants and Analysis Samples

Of 177 individuals who completed the online screening, 48 completed the adaptation night and medical examination. Of these, 7 discontinued before the first experimental visit, for personal reasons or non-adherence to the ambulatory sleep-wake schedule. Forty-one healthy German-speaking adults (18-34 years, *M* = 23.7, *SD* = 4.3; 24 women) therefore completed at least one experimental visit; of these, 38 completed all three visits, 2 completed two, and 1 completed one, yielding 119 completed experimental visits in total. Thus, 10 of the 48 participants did not complete the full protocol (7 before the first visit and 3 after one or two). Twenty participants were assigned to the active group and 21 to the inactive group. Agreed habitual bedtimes ranged from 22:15 to 00:30 (*M* = 23:20, *SD* = 40 min).

For 9 visits, a reliable DLMO could not be estimated from the melatonin profile on one or both evenings. Hence, a change in DLMO could be determined for 110 of the 119 visits. Because the models additionally required valid light and heart-rate data, the primary analysis was based on 81 visits from 36 participants (68.1% of completed visits). Among visits with an available change in DLMO, 1 was excluded due to missing light recordings, 28 due to missing heart-rate data, and 0 due to both. Analysis samples for the remaining outcomes are reported with the corresponding results and are summarised in Table 1.

**Table 1.** Participant Flow and Analysis Samples.

| Stage | Participants | Visits | Observations |
|---|---|---|---|
| Completed online screening | 177 | — | — |
| Completed adaptation night | 48 | — | — |
| Discontinued before first visit | 7 | — | — |
| Completed ≥ 1 experimental visit | 41 | 119 | — |
| Completed all three visits | 38 | 114 | — |
| **Analysed…** |  |  |  |
| **ΔDLMO (primary)** | **36** | **81** | **81** |
| Evening time course | 37 | 84 | 827 |
| Daytime time course (sleepiness) | 37 | 88 | 1053 |
| Daytime time course (mood, wellbeing) | 37 | 87 | 1040 |
| Daytime time course (exertion) | 37 | 87 | 1024 |

*Note.* Analysis samples differ because complete-case deletion was applied per model rather than globally. DLMO = dim-light melatonin onset; ΔDLMO = Evening-2 − Evening-1. Observations refer to the units entered into the respective model (visits for visit-level outcomes, individual assessments for time-course outcomes).

Of the 110 visits with an estimable change in DLMO, 66 (60%) showed a phase advance (negative change), 43 (39%) a phase delay, and one no change. The largest mean advances occurred in the daylight condition (*M* = −14.56, *SD* = 31.43 min) and in the active group (*M* = −14.22, *SD* = 24.26 min), whereas the remaining two light conditions and the inactive group showed mean changes near zero (Table 2; Figure 2). Individual variability was substantial across all conditions and groups (*SD*s 23.93-39.94 min; for the change in DLMO against measured illuminance, see Figure S6). Descriptive statistics for all study variables are given in Table 2. summarise participant means calculated across available conditions. Light exposure values are median photopic illuminance across the 8-h exposure window, with the glasses values being filter-corrected. HRR values are participant means of visit-level median heart rate reserve. KSS = Karolinska Sleepiness Scale; PVT = psychomotor vigilance task. Daytime and evening outcomes are participant grand means across available conditions and assessments; evening outcomes include both evenings, and PVT sessions were first averaged within participant and condition. *n* refers to the number of participants contributing to each entry.

**Figure 2.**
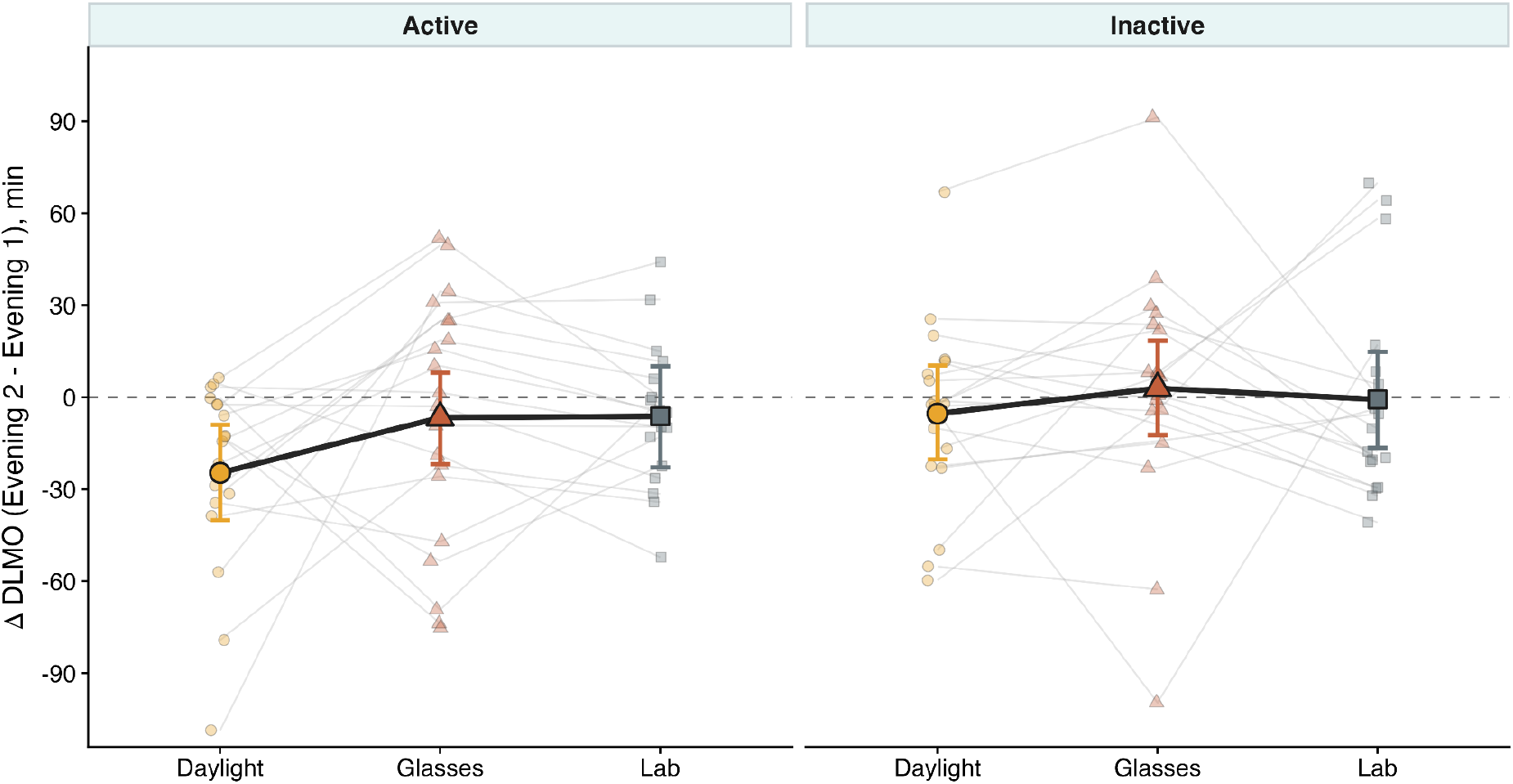
Change in DLMO Across Light Conditions and Activity Groups. *Note.* Each point shows the change in DLMO (Evening-2 – Evening-1) for one visit; lines connect visits of the same participant. Large symbols and error bars show posterior cell means and 95% credible intervals from a descriptive categorical model fitted to all 110 visits with an estimable change in DLMO. The dashed line marks no change. Negative values indicate a phase advance. These categorical estimates complement the continuous-predictor analysis, from which the primary inferential conclusions were drawn.

**Table 2.** Descriptive Statistics for Sample Characteristics, Predictors, and Outcomes.

| Variable | *n* | *M* | *SD* | Range |
|---|---|---|---|---|
| **Sample characteristics** |  |  |  |  |
| Age (years) | 41 | 23.7 | 4.3 | 18 to 34 |
| Sex (women), *n* (%) | 24 (58.5) | — | — | — |
| Habitual bedtime (hh:mm) | 41 | 23:20 | 0:40 | 22:15 to 00:30 |
| **Photopic illuminance (log<sub>10</sub>(lx + 1))** |  |  |  |  |
| Daylight | 39 | 2.94 | 0.38 | 2.01 to 3.58 |
| Glasses | 41 | 2.52 | 0.39 | 1.73 to 3.15 |
| Laboratory light | 37 | 1.74 | 0.35 | 1.11 to 2.54 |
| **Photopic illuminance (lx)** |  |  |  |  |
| Daylight | 39 | 1195.6 | 959.1 | 101.56 to 3758.73 |
| Glasses | 41 | 460.3 | 337.0 | 52.58 to 1404.89 |
| Laboratory light | 37 | 75.4 | 71.4 | 11.78 to 343.69 |
| **Heart rate reserve, median HRR (%)** |  |  |  |  |
| Active group | 18 | 32.66 | 7.67 | 20.26 to 46.00 |
| Inactive group | 19 | 2.69 | 2.63 | 0.00 to 8.41 |
| **ΔDLMO (min)** |  |  |  |  |
| Daylight | 37 | -14.56 | 31.43 | -108.60 to 66.90 |
| Glasses | 38 | -2.16 | 39.94 | -99.60 to 91.20 |
| Laboratory light | 35 | -3.10 | 28.92 | -52.20 to 69.90 |
| Active group | 20 | -14.22 | 24.26 | -75.30 to 23.55 |
| Inactive group | 21 | -1.45 | 23.93 | -58.95 to 52.80 |
| **Daytime outcomes (grand *M*)** |  |  |  |  |
| Sleepiness (KSS) | 41 | 2.69 | 0.87 | 1.03 to 4.56 |
| Mood | 41 | 4.15 | 0.48 | 3.06 to 5.00 |
| Wellbeing | 41 | 4.13 | 0.55 | 3.00 to 5.00 |
| Perceived exertion (Borg) | 41 | 2.25 | 1.32 | 1.00 to 5.17 |
| **Evening outcomes (grand *M*)** |  |  |  |  |
| Sleepiness (KSS) | 41 | 5.49 | 1.13 | 1.80 to 7.67 |
| Mood | 41 | 3.87 | 0.53 | 2.92 to 4.98 |
| Wellbeing | 41 | 3.97 | 0.58 | 2.92 to 5.00 |
| PVT median reaction time (ms) | 41 | 318.63 | 55.47 | 227.61 to 432.65 |
| PVT response speed (s<sup>−1</sup>) | 41 | 3.24 | 0.57 | 2.35 to 4.38 |

*Note.* DLMO = dim-light melatonin onset; ΔDLMO = Evening-2 – Evening-1, with negative values indicating a phase advance. Condition-specific ΔDLMO statistics summarise visit-level shifts; activity group statistics summarise participant means calculated across available conditions. Light exposure values are median photopic illuminance across the 8-h exposure window, with the glasses values being filter-corrected. HRR values are participant means of visit-level median heart rate reserve. KSS = Karolinska Sleepiness Scale; PVT = psychomotor vigilance task. Daytime and evening outcomes are participant grand means across available conditions and assessments; evening outcomes include both evenings, and PVT sessions were first averaged within participant and condition. *n* refers to the number of participants contributing to each entry.

### Manipulation Checks and Baseline Equivalence

Manipulation checks confirmed that the conditions produced the intended contrasts. Photopic illuminance differed strongly across the three light conditions (Friedman χ²(2) = 46.89, *p* < .001, *BF*₁₀ > 1000) and all pairwise contrasts differed (all Holm-corrected *p*s ≤ .001, *BF*₁₀s ≥ 100), confirming the intended separation of exposure. Median heart rate reserve during the intervention was substantially higher in the active than the inactive group (*W* = 342, *p*< .001, *BF*₁₀ > 1000) and did not differ across light conditions (χ²(2) = 3.91, *p* = .142, *BF*₁₀ = 0.17), indicating that physical activity intensity was comparable across the light manipulations. Perceived exertion was correspondingly higher in the active group (*W* = 420, *p* < .001, *BF*₁₀ > 1000) and varied modestly across light conditions (χ²(2) = 10.26, *p* = .006, *BF*₁₀ = 4.00). Pairwise, however, only the glasses–laboratory contrast was supported by the Bayes factor (Holm-corrected *p* = .007, *BF*₁₀ = 6.84), whereas the daylight-laboratory contrast was inconclusive (Holm-corrected *p* = .022, *BF*₁₀ = 1.42).

Condition effects on daytime sleepiness (χ²(2) = 8.93, *p* = .011, *BF*₁₀ = 0.83) and wellbeing (χ²(2) = 2.31, *p* = .314, *BF*₁₀ = 0.96) were inconclusive, although pairwise, sleepiness differed between the daylight and laboratory conditions (Holm-corrected *p* = .022, *BF*₁₀ = 4.07). Mood, in contrast, differed across conditions (χ²(2) = 8.38, *p* = .015, *BF*₁₀ = 7.25), primarily because of the daylight-laboratory contrast (Holm-corrected *p* = .003, *BF*₁₀ = 18.02). No group differences were evident for these outcomes (all *p*s ≥ .548, *BF*₁₀s ≤ 0.34). Full manipulation-check statistics are reported in Supplementary Results S1 (Table S4; Figures S4-S5).

Prior light history did not differ across light conditions (mean photopic illuminance and cumulative exposure: both χ²(2) = 1.00, *p*s = .607, *BF*₁₀s ≤ 0.17). Between activity groups, mean illuminance and cumulative exposure did not differ significantly (both *W* = 156, *p* = .165), although the Bayes factor was inconclusive (*BF*₁₀ = 0.59). Time above 250 lx (*W* = 124, *p* = .026) and above 1,000 lx (*W* = 132, *p* = .043) differed at *p* < .05 (uncorrected), but here, too, the Bayes factors were inconclusive (*BF*₁₀ = 1.03 and 0.68), so the data do not establish a group difference in prior light history.

Furthermore, baseline DLMO on Evening-1 was comparable across light conditions (χ²(2) = 0.32, *p* = .852, *BF*₁₀ = 0.09) and activity groups (*W* = 217, *p* = .867, *BF*₁₀ = 0.31). Evening-1 subjective ratings likewise showed no evident baseline differences (all *BF*₁₀ ≤ 0.33). One exception emerged for psychomotor vigilance: the active group already responded faster than the inactive group at Evening-1, before any intervention (all PVT metrics *p*s ≤ .001, *BF*₁₀s ≥ 14.96), whereas PVT did not show evident differences across light conditions (all *BF*₁₀ ≤ 0.47). This pre-existing group difference is taken into account when interpreting the evening PVT findings (see *Evening Time Course* and Results S5)

### Primary Outcome: Change in DLMO

The primary model of the change in DLMO was based on 81 visits from 36 participants (Table 3 and Table S5; Figure 3) and showed satisfactory sampling behaviour (maximum *R^* = 1.001, minimum ESS = 5,161, no divergent transitions). On average, DLMO occurred approximately 8 min earlier on Evening-2 than on Evening-1 (*b* = −8.05 min, 95% HDI [−15.90, −0.42]), a small overall phase advance. Residual variability was large relative to this mean (*SD* = 29.81 min, 95% HDI [24.34, 35.58]), indicating that the change in DLMO differed considerably between visits.

**Figure 3.**
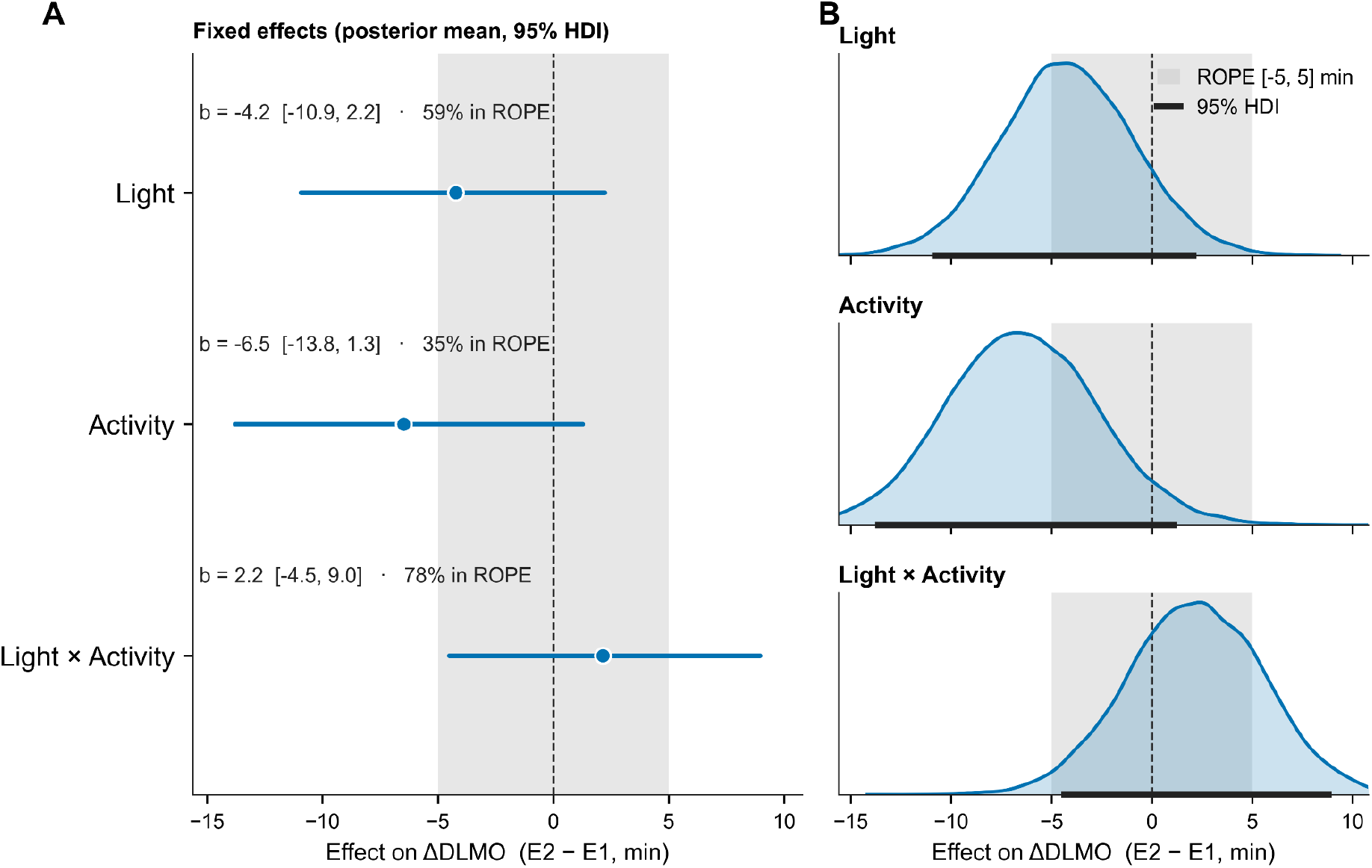
Posterior Distributions of the Effects of Light Exposure, Physical Activity, and Their Interaction on the Change in DLMO. *Note.* (A) Points and horizontal lines show posterior means and 95% highest density intervals (HDIs), respectively; annotations report *b* [95% HDI] and the percentage of the posterior within the region of practical equivalence (ROPE; −5 to 5 min). (B) Curves show the corresponding posterior densities; shaded bands indicate the ROPE, black horizontal bars the 95% HDIs, and dashed vertical lines zero. Light exposure and physical activity effects are expressed per 1-*SD* increase, with the other predictor at its mean; the interaction represents the change in the light-exposure effect per 1-*SD* increase in physical activity. Negative values indicate a phase advance.

**Table 3.** Posterior Estimates for the Primary Outcome, the Change in DLMO.

| Effect | *b* | 95% HDI | *pd* | ROPE % | *BF*<sub>10</sub> | Evidence |
|---|---|---|---|---|---|---|
| Intercept | -8.05 | [-15.90, -0.42] | — | — | — | — |
| Light | -4.22 | [-10.93, 2.22] | .90 | 58.6 | 0.70 | Inconclusive |
| Activity | -6.48 | [-13.79, 1.27] | .95 | 34.5 | 1.56 | Inconclusive |
| Light × Activity | 2.15 | [-4.50, 8.96] | .74 | 77.5 | 0.39 | Inconclusive |

*Note*. *N* = 81 visits from 36 participants. DLMO = dim-light melatonin onset; ΔDLMO = Evening-2 − Evening-1, negative values indicating a phase advance. *b* = posterior mean. Light and activity coefficients give the expected outcome difference per +1 *SD* at the mean of the other predictor; the light × activity coefficient gives the change in the light coefficient per +1 *SD* in activity; HDI = highest density interval; *pd* = probability of direction; ROPE % = percentage of the posterior within the region of practical equivalence, here ±5 min; *BF*<sub>10</sub> = Savage–Dickey density ratio against a point null. Both predictors were *z*-standardised. Residual *SD* = 29.81 min, 95% HDI [24.34, 35.58]. Sampling diagnostics: maximum *R̂* = 1.001, minimum ESS = 5,161, no divergent transitions. Leave-one-out cross-validation did not distinguish the candidate models (null vs. best: ΔELPD = 1.62, *SE* = 2.21; no Pareto-*k̂* warnings). Evidence classification: *Inconclusive*, 1/3 < *BF*<sub>10</sub> < 3 (data do not discriminate presence from absence of an effect); *Evidence for*, *BF*<sub>10</sub> ≥ 3 with most of the posterior outside the ROPE, 95% HDI excludes zero; *Present but negligible*, *BF*<sub>10</sub> ≥ 3 but ≥ 95% of the posterior within the ROPE (effect supported yet practically trivial); *Evidence against*, *BF*<sub>10</sub> ≤ 1/3 with the posterior concentrated within the ROPE. Evidence classification was not applied to the intercept, for which no ROPE or Bayes factor was defined.

For light exposure, the posterior mean corresponded to a 4-min larger phase advance per 1-*SD* increase in log₁₀ photopic illuminance (*b* = −4.22 min, 95% HDI [−10.93, 2.22], 58.6% in ±5-min ROPE, *BF*₁₀ = 0.70), however, the evidence was inconclusive, so the data neither support nor rule out a light effect. The posterior mean for physical activity likewise corresponded to a phase advance (*b* = −6.48 min per *SD*, 95% HDI [−13.79, 1.27], 34.5% in ROPE, *BF*₁₀ = 1.56). Although two-thirds of the posterior lay outside the ROPE, so that an effect of this size would be practically relevant, the evidence again remained inconclusive. The light × activity interaction was small and imprecisely estimated (*b* = 2.15 min, 95% HDI [−4.50, 8.96], 77.5% in ROPE, *BF*₁₀ = 0.39), providing no support for the hypothesised light-activity synergy while not excluding an interaction of a priori plausible size.

PSIS-LOO did not distinguish the candidate models. Even the null model predicted the change in DLMO about as well as the best-ranked model (ΔELPD = 1.62, *SE* = 2.21, Table S15). Hence, adding light exposure, physical activity, or their interaction did not improve out-of-sample prediction.

### Secondary Outcomes

#### Evening Time Course

All secondary outcomes were exploratory (see *Method – Data Analysis*). Evening models were based on 827 observations from 37 participants across 84 visits (Table 4; Figure S7; full estimates in Table S9). These outcomes were sampled at ten assessments per evening, beginning 5 h until 30 min before habitual bedtime (see *Method – Procedure*). Subjective sleepiness increased across the evening (*b* = 0.30 KSS points per assessment, 95% HDI [0.25, 0.35], *BF*₁₀ > 100), indicating that participants rated themselves progressively sleepier as the evening advanced. The association between daytime light exposure and evening sleepiness was inconclusive (*b* = 0.25 per *SD*, 95% HDI [0.03, 0.45], 99.0% in ROPE, *BF*₁₀ = 2.76), so the data neither establish nor rule out a link between brighter days and greater evening sleepiness. The light × activity interaction was likewise inconclusive (*b* = 0.15, 95% HDI [−0.07, 0.35], 99.9% in ROPE, *BF*₁₀ = 0.46). For physical activity, in contrast, the data provided moderate evidence against a practically relevant association with evening sleepiness (*b* = 0.05, 95% HDI [−0.32, 0.43], 98.8% in ROPE, *BF*₁₀ = 0.33). Derived contrasts are reported in Table S10.

**Table 4.**
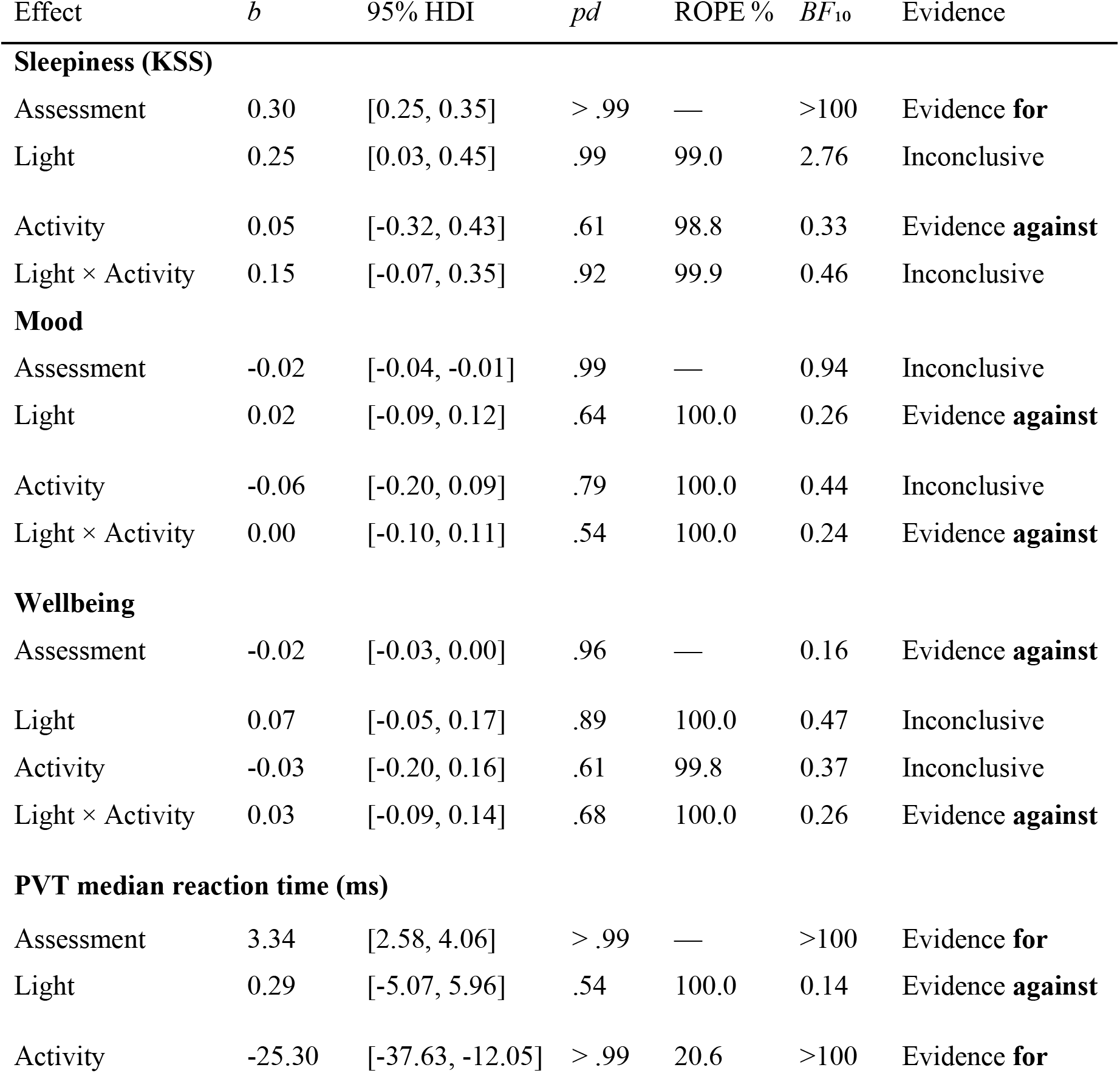
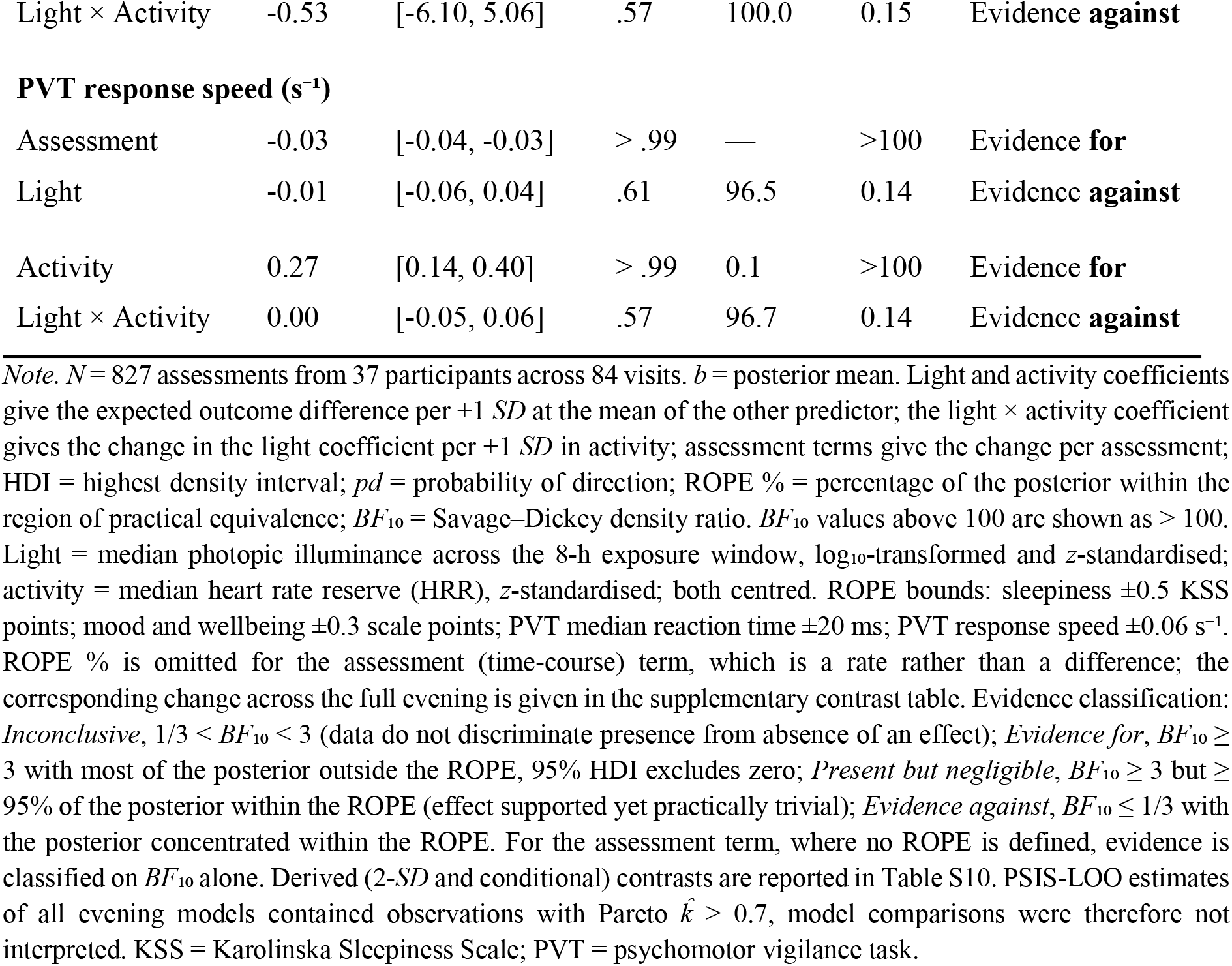
Posterior Estimates for Evening Time-Course Outcome.

| Effect | *b* | 95% HDI | *pd* | ROPE % | *BF*<sub>10</sub> | Evidence |
|---|---|---|---|---|---|---|
| **Sleepiness (KSS)** |  |  |  |  |  |  |
| Assessment | 0.30 | [0.25, 0.35] | >.99 | — | >100 | Evidence **for** |
| Light | 0.25 | [0.03, 0.45] | .99 | 99.0 | 2.76 | Inconclusive |
| Activity | 0.05 | [-0.32, 0.43] | .61 | 98.8 | 0.33 | Evidence **against** |
| Light × Activity | 0.15 | [-0.07, 0.35] | .92 | 99.9 | 0.46 | Inconclusive |
| **Mood** |  |  |  |  |  |  |
| Assessment | -0.02 | [-0.04, -0.01] | .99 | — | 0.94 | Inconclusive |
| Light | 0.02 | [-0.09, 0.12] | .64 | 100.0 | 0.26 | Evidence **against** |
| Activity | -0.06 | [-0.20, 0.09] | .79 | 100.0 | 0.44 | Inconclusive |
| Light × Activity | 0.00 | [-0.10, 0.11] | .54 | 100.0 | 0.24 | Evidence **against** |
| **Wellbeing** |  |  |  |  |  |  |
| Assessment | -0.02 | [-0.03, 0.00] | .96 | — | 0.16 | Evidence **against** |
| Light | 0.07 | [-0.05, 0.17] | .89 | 100.0 | 0.47 | Inconclusive |
| Activity | -0.03 | [-0.20, 0.16] | .61 | 99.8 | 0.37 | Inconclusive |
| Light × Activity | 0.03 | [-0.09, 0.14] | .68 | 100.0 | 0.26 | Evidence **against** |
| **PVT median reaction time (ms)** |  |  |  |  |  |  |
| Assessment | 3.34 | [2.58, 4.06] | >.99 | — | >100 | Evidence **for** |
| Light | 0.29 | [-5.07, 5.96] | .54 | 100.0 | 0.14 | Evidence **against** |
| Activity | -25.30 | [-37.63, -12.05] | >.99 | 20.6 | >100 | Evidence **for** |
| Light × Activity | -0.53 | [-6.10, 5.06] | .57 | 100.0 | 0.15 | Evidence **against** |
| **PVT response speed (s<sup>−1</sup>)** |  |  |  |  |  |  |
| Assessment | -0.03 | [-0.04, -0.03] | >.99 | — | >100 | Evidence **for** |
| Light | -0.01 | [-0.06, 0.04] | .61 | 96.5 | 0.14 | Evidence **against** |
| Activity | 0.27 | [0.14, 0.40] | >.99 | 0.1 | >100 | Evidence **for** |
| Light × Activity | 0.00 | [-0.05, 0.06] | .57 | 96.7 | 0.14 | Evidence **against** |

*Note*. *N* = 827 assessments from 37 participants across 84 visits. *b* = posterior mean. Light and activity coefficients give the expected outcome difference per +1 *SD* at the mean of the other predictor; the light × activity coefficient gives the change in the light coefficient per +1 *SD* in activity; assessment terms give the change per assessment; HDI = highest density interval; *pd* = probability of direction; ROPE % = percentage of the posterior within the region of practical equivalence; *BF*<sub>10</sub> = Savage–Dickey density ratio. *BF*<sub>10</sub> values above 100 are shown as > 100. Light = median photopic illuminance across the 8-h exposure window, log<sub>10</sub>-transformed and *z*-standardised; activity = median heart rate reserve (HRR), *z*-standardised; both centred. ROPE bounds: sleepiness ±0.5 KSS points; mood and wellbeing ±0.3 scale points; PVT median reaction time ±20 ms; PVT response speed ±0.06 s<sup>−1</sup>. ROPE % is omitted for the assessment (time-course) term, which is a rate rather than a difference; the corresponding change across the full evening is given in the supplementary contrast table. Evidence classification: *Inconclusive*, 1/3 < *BF*<sub>10</sub> < 3 (data do not discriminate presence from absence of an effect); *Evidence for*, *BF*<sub>10</sub> ≥ 3 with most of the posterior outside the ROPE, 95% HDI excludes zero; *Present but negligible*, *BF*<sub>10</sub> ≥ 3 but ≥ 95% of the posterior within the ROPE (effect supported yet practically trivial); *Evidence against*, *BF*<sub>10</sub> ≤ 1/3 with the posterior concentrated within the ROPE. For the assessment term, where no ROPE is defined, evidence is classified on *BF*<sub>10</sub> alone. Derived (2-*SD* and conditional) contrasts are reported in Table S10. PSIS-LOO estimates of all evening models contained observations with Pareto *k̂* > 0.7, model comparisons were therefore not interpreted. KSS = Karolinska Sleepiness Scale; PVT = psychomotor vigilance task.

None of the coefficients for evening mood and wellbeing was evidenced, and all posteriors lay almost entirely within the ROPE, so any associations were practically negligible. For mood, the data provided moderate evidence against practically relevant effects of light exposure (*b* = 0.02, 95% HDI [0-.09, 0.12], 100% in ROPE, *BF*₁₀ = 0.26) and of the light × activity interaction (*b* = 0.00, 95% HDI [−0.10, 0.11], 100% in ROPE, *BF*₁₀ = 0.24), whereas the activity estimate was inconclusive (*b* = −0.06 per *SD*, 95% HDI [−0.20, 0.09], 100% in ROPE, *BF*₁₀ = 0.44). For wellbeing, conversely, only the light × activity interaction yielded moderate evidence against a practically relevant effect (*b* = 0.03, 95% HDI [−0.09, 0.14], 100% in ROPE, *BF*₁₀ = 0.26), whereas the light (*b* = 0.07 per *SD*, 95% HDI [−0.05, 0.17], 100% in ROPE, *BF*₁₀ = 0.47) and activity estimates (*b* = −0.03 per *SD*, 95% HDI [−0.20, 0.16], 99.8% in ROPE, *BF*₁₀ = 0.37) were inconclusive.

Psychomotor vigilance deteriorated across the evening, with slower median RTs at later assessments (*b* = 3.34 ms per assessment, 95% HDI [2.58, 4.06], *BF*₁₀ > 100). Higher physical activity was associated with substantially faster responding (median RT: *b* = −25.30 ms per *SD*, 95% HDI [−37.63, −12.05], 20.6% in ROPE, *BF*₁₀ > 100; response speed: *b* = 0.27 s⁻¹ per *SD*, 95% HDI [0.14, 0.40], 0.1% in ROPE, *BF*₁₀ > 100), meaning that participants with higher heart rate reserve during the day responded faster on the PVT on Evening-2. In contrast, daytime light exposure was not associated with either metric, with moderate evidence against a practically relevant effect (median RT: *b* = 0.29 ms, 95% HDI [−5.07, 5.96], 100.0% in ROPE, *BF*₁₀ = 0.14; response speed: *b* = −0.01 s⁻¹, 95% HDI [−0.06, 0.04], 96.5% in ROPE, *BF*₁₀ = 0.14).

Because the activity groups already differed in vigilance on Evening-1 (see *Manipulation Checks and Baseline Equivalence*), we additionally fitted a baseline-adjusted model that controlled for Evening-1 performance (807 observations from 37 participants across 82 visits; Results S5). In this model, the activity association was no longer evidenced. For median RT, these data provided moderate evidence against a practically relevant effect (*b* = - 2.67 ms per *SD*, 95% HDI [−9.41, 3.92], 100% in ROPE, *BF*₁₀ = 0.23). For response speed, the Bayes factor likewise favoured the null (*b* = 0.03 s⁻¹ per *SD*, 95% HDI [−0.03, 0.10], 80.3% in ROPE, *BF*₁₀ = 0.27), but with a fifth of the posterior outside the ROPE, a practically relevant effect cannot be ruled out. Evening-1 performance, in contrast, strongly predicted Evening-2 performance (median RT: *b* = 49.74 ms per *SD*, 95% HDI [42.95, 56.51]; response speed: *b* = 0.50 s⁻¹ per *SD*, 95% HDI [0.44, 0.56]; both *BF*₁₀ > 100). Hence, the unadjusted activity association is more plausibly explained by pre-existing between-person differences in vigilance than by the activity manipulation.

For all evening outcomes, the assessment × light and assessment × activity interactions yielded evidence against a dependence of the evening trajectory on either predictor (all *BF*₁₀ ≤ 0.14; Table S9). Model comparisons were not interpreted because PSIS-LOO estimates of all evening models had Pareto-*k^* > 0.7 (Table S15).

#### Daytime Time Course

Daytime models were based on 1,053 observations from 37 participants across 88 visits for sleepiness, 1,040 observations from 37 participants across 87 visits for mood and wellbeing, and 1,024 observations from 37 participants across 87 visits for perceived exertion (Table 5; Figure S8; full estimates in Table S13). These outcomes were assessed twelve times at 30-min intervals during and immediately after the physical activity window (see *Method – Procedure*). Daytime subjective sleepiness increased modestly across the day (*b* = 0.09 KSS points per assessment, 95% HDI [0.06, 0.13], *BF*₁₀ > 100). The evidence regarding light exposure (*b* = - 0.16 per *SD*, 95% HDI [−0.32, 0.01], 100% in ROPE, *BF*₁₀ = 1.09) and physical activity (*b* = 0.08 per *SD*, 95% HDI [−0.18, 0.34], 99.9% in ROPE, *BF*₁₀ = 0.34) was inconclusive. For the light × activity interaction, the data provided moderate evidence against a practically relevant effect (*b* = −0.06 KSS points, 95% HDI [−0.22, 0.10], 100% in ROPE, *BF*₁₀ = 0.24).

**Table 5.** Posterior Estimates for Daytime Time-Course Outcomes.

| Effect | *b* | 95% HDI | *pd* | ROPE % | *BF*<sub>10</sub> | Evidence |
|---|---|---|---|---|---|---|
| **Sleepiness (KSS)** |  |  |  |  |  |  |
| Assessment | 0.09 | [0.06, 0.13] | >.99 | — | >100 | Evidence **for** |
| Light | -0.16 | [-0.32, 0.01] | .97 | 100.0 | 1.09 | Inconclusive |
| Activity | 0.08 | [-0.18, 0.34] | .72 | 99.9 | 0.34 | Inconclusive |
| Light × Activity | -0.06 | [-0.22, 0.10] | .76 | 100.0 | 0.24 | Evidence **against** |
| **Mood** |  |  |  |  |  |  |
| Assessment | -0.02 | [-0.03, 0.00] | .97 | — | 0.19 | Evidence **against** |
| Light | 0.14 | [0.02, 0.26] | .99 | 99.5 | 3.32 | Present but negligible |
| Activity | 0.01 | [-0.14, 0.17] | .57 | 100.0 | 0.30 | Evidence **against** |
| Light × Activity | 0.09 | [-0.02, 0.21] | .94 | 100.0 | 0.81 | Inconclusive |
| **Wellbeing** |  |  |  |  |  |  |
| Assessment | -0.02 | [-0.03, 0.00] | .98 | — | 0.24 | Evidence **against** |
| Light | 0.10 | [-0.01, 0.21] | .96 | 100.0 | 0.92 | Inconclusive |
| Activity | 0.02 | [-0.15, 0.20] | .62 | 99.9 | 0.35 | Inconclusive |
| Light × Activity | 0.06 | [-0.05, 0.17] | .85 | 100.0 | 0.37 | Inconclusive |
| **Perceived exertion (Borg)** |  |  |  |  |  |  |
| Assessment | -0.04 | [-0.06, -0.01] | >.99 | — | 0.75 | Inconclusive |
| Light | -0.15 | [-0.29, -0.02] | .99 | 100.0 | 1.17 | Inconclusive |
| Activity | 1.04 | [0.82, 1.25] | >.99 | 37.1 | >100 | Evidence **for** |
| Light × Activity | -0.10 | [-0.23, 0.03] | .93 | 100.0 | 0.30 | Evidence **against** |

*Note.* Analysis samples: sleepiness *N* = 1,053 assessments from 37 participants across 88 visits; mood and wellbeing *N* = 1,040 from 37 participants across 87 visits; perceived exertion *N* = 1,024 from 37 participants across 87 visits. *b* = posterior mean. Light and activity coefficients give the expected outcome difference per +1 *SD* at the mean of the other predictor; the light × activity coefficient gives the change in the light coefficient per +1 *SD* in activity; assessment terms give the change per assessment; HDI = highest density interval; *pd* = probability of direction; ROPE % = percentage of the posterior within the region of practical equivalence; *BF*<sub>10</sub> = Savage–Dickey density ratio. *BF*<sub>10</sub> values above 100 are shown as > 100. Light = median photopic illuminance across the 8-h exposure window, log<sub>10</sub>-transformed and *z*-standardised; activity = median heart rate reserve (HRR), *z*-standardised; both centred. ROPE bounds: sleepiness ±0.5 KSS points; mood and wellbeing ±0.3 scale points; perceived exertion ±1 Borg point. ROPE % is omitted for the assessment (time-course) term, which is a rate rather than a difference; the corresponding change across the full day is given in the supplementary contrast table. Evidence classification: *Inconclusive*, 1/3 < *BF*<sub>10</sub> < 3; *Evidence for*, *BF*<sub>10</sub> ≥ 3 with most of the posterior outside the ROPE, 95% HDI excludes zero; *Present but negligible*, *BF*<sub>10</sub> ≥ 3 but ≥ 95% of the posterior within the ROPE; *Evidence against*, *BF*<sub>10</sub> ≤ 1/3 with the posterior concentrated within the ROPE. For the assessment term, where no ROPE is defined, evidence is classified on *BF*<sub>10</sub> alone. Derived contrasts are reported in Table S14. PSIS-LOO identified no predictively superior model for mood or perceived exertion; comparisons for sleepiness and wellbeing raised Pareto-*k̂* warnings. KSS = Karolinska Sleepiness Scale.

Daytime mood was slightly better at higher light exposure (*b* = 0.14 per *SD*, 95% HDI [0.02, 0.26], 99.5% in ROPE, *BF*₁₀ = 3.32). Although the Bayes factor provided moderate evidence for this association, almost the entire posterior per 1-*SD* increase lay within the ROPE. Between high and low light exposure (±1 *SD*), however, mood differed by 0.28 scale points (95% HDI [0.04, 0.52]), with only 54.8% of this posterior inside the ROPE. Hence, a practically relevant difference between high and low light exposure cannot be ruled out. Physical activity, in contrast, showed moderate evidence against a practically relevant association with mood (*b* = 0.01 per *SD*, 95% HDI [−0.14, 0.17], 100% in ROPE, *BF*₁₀ = 0.30), and the light × activity interaction was inconclusive (*b* = 0.09, 95% HDI [−0.02, 0.21], 100% in ROPE, *BF*₁₀ = 0.81). For wellbeing, all estimates were inconclusive (light: *b* = 0.10 per *SD*, 95% HDI [−0.01, 0.21], 100% in ROPE, *BF*₁₀ = 0.92; activity: *b* = 0.02 per *SD*, 95% HDI [−0.15, 0.20], 99.9% in ROPE, *BF*₁₀ = 0.35; interaction: *b* = 0.06, 95% HDI [−0.05, 0.17], 100% in ROPE, *BF*₁₀ = 0.37).

Derived contrasts are reported in Table S14. As expected, higher physical activity was associated with markedly greater perceived exertion (*b* = 1.04 Borg points per *SD*, 95% HDI [0.82, 1.25], 37.1% in ROPE, *BF*₁₀ > 100), corresponding to a difference of 2.07 points between high and low activity (95% HDI [1.63, 2.51], 0% in ROPE). Hence, participants clearly perceived the activity manipulation. In contrast, the light-exposure estimate was inconclusive (*b* = −0.15 per *SD*, 95% HDI [−0.29, - 0.02], *BF*₁₀ = 1.17). Because its posterior lay entirely within the ±1-point ROPE, both per 1 *SD* and for the high-versus-low contrast (Table S14), any association would be practically negligible.

For mood and perceived exertion, PSIS-LOO identified no predictively superior model (null vs. additive model: mood, ΔELPD = 0.15, *SE* = 1.34; perceived exertion, ΔELPD = 5.71, *SE* = 5.20). Comparisons for sleepiness and wellbeing had Pareto-*k^* warnings and were not interpreted (Table S15).

### Sensitivity and Additional Exploratory Analyses

The primary conclusion was robust to a range of analytical choices. Re-estimating the model of the change in DLMO with a Student’s *t*-likelihood, wider priors on the fixed effects, cumulative rather than median photopic illuminance, corrected laboratory light values, or age and sex as covariates left the pattern unchanged: the light-exposure and physical-activity estimates continued to correspond to modest phase advances, but all 95% HDIs included zero, the Bayes factors remained inconclusive, and no candidate model was predictively superior where PSIS-LOO was reliable. The only exception concerned the light × activity interaction under the wider prior, for which *BF*₁₀ decreased from 0.39 to 0.26, below the ⅓ threshold. Yet, such a shift towards the null is expected when the prior is widened. A categorical parameterisation of light condition and activity group yielded the same pattern. Full specifications and results are provided in Supplementary Method S7 and Results S2 (Tables S5-S7).

In the additional exploratory analysis (87 visits from 37 participants), the change in evening melatonin secretion (log₁₀ ratio of Evening-2 to Evening-1 AUC) yielded results consistent with those for the change in DLMO. The light-exposure association was inconclusive (*b* = 0.029 per *SD*, 95% HDI [0.00, 0.06], 80.6% in ROPE, *BF*₁₀ = 2.30), the data provided moderate evidence against a practically relevant activity association (*b* = 0.00 per *SD*, 95% HDI [−0.03, 0.03], 99.3% in ROPE, *BF*₁₀ = 0.32), and the light × activity interaction was inconclusive (*b* = −0.01, 95% HDI [−0.04, 0.02], 98.1% in ROPE, *BF*₁₀ = 0.40; Results S3, Table S8). Hence, neither the timing nor the amount of evening melatonin secretion was conclusively associated with daytime light exposure or physical activity.

## Discussion

In this mixed-design experiment, a single day of natural daylight exposure, with or without physical activity, did not conclusively change circadian phase. Higher light exposure and higher physical activity were each associated with a slightly earlier DLMO on the second evening. However, the data could neither confirm nor rule out either association or their interaction, and adding either predictor did not improve out-of-sample prediction of the change in DLMO. The exploratory secondary outcomes showed a similar pattern. Associations with light and activity were predominantly inconclusive or practically negligible, and evening psychomotor vigilance provided evidence against a practically relevant association with daytime light. Taken together, these results suggest that a single day of moderate daylight exposure may be insufficient to measurably change the phase of the human circadian system, and that repeated, habitual exposure may matter more.

The inconclusive change in DLMO should be interpreted in the context of the intensity and circadian timing of the light exposure. Across the 8-h window, median photopic illuminance averaged 1,195.6 lx (*SD* = 959.1) in the daylight condition, 460.3 lx (*SD* = 337.0) in the glasses condition, and 75.4 lx (*SD* = 71.4) in the laboratory condition. The manipulation therefore produced a substantial exposure contrast, although even the daylight condition remained modest relative to established phase-shifting protocols, likely in part because much of the outdoor time was spent under tree cover and because the sensor, oriented to approximate gaze direction, captured light along the line of sight rather than from the brighter sky above. In a controlled morning protocol, 3-h exposures to 5,000-K fluorescent light at 3,000, 6,000, or 12,000 lx, beginning 1 h after waking, produced same-cycle DLMO advances of 15, 26, and 27 min, respectively, whereas 750 and 1,500 lx did not produce significant shifts^71^. Khalsa et al^21^ further illustrated the importance of circadian timing. They established the human light phase-response curve using a single 6.7-h exposure alternating between approximately 10,000 lx during fixed gaze and 5,000–9,000 lx during free gaze. Exposures centred after the core-body-temperature minimum advanced circadian phase, whereas those centred before it delayed it. Our exposure window extended from approximately 1.5 h to 9.5 h after waking. Rather than isolating the immediate post-awakening period, it thus covered the middle of the day, when the circadian system appears comparatively insensitive to phase-shifting light^22^. Although this timing was individualised to each participant’s habitual sleep-wake schedule, it was not directly anchored to an internal phase marker such as DLMO or the core-body-temperature minimum. Moreover, light exposure and physical activity were each summarised as a single median per visit, across the exposure window and the activity period, respectively. The analysis could therefore not resolve when within these periods participants received most light or activity, nor whether the timing of exposure relative to circadian phase moderated the response. The moderate intensity and broad temporal distribution of the exposure may thus have reduced the consistency of the change in DLMO at the group level.

Beyond intensity and timing, the circadian response to light appears to also depend on duration and repetition, although the duration-response relationship is markedly nonlinear. In a late-evening phase-delay protocol, a single 12-min exposure to approximately 10,000 lx shifted circadian phase by 1.07 h, whereas extending the exposure to 4 h increased the shift to only 2.65 h, indicating diminishing returns with increasing duration^25^. That protocol, however, examined phase delays at an intensity substantially higher than that achieved in our study. Robust phase advances have generally been observed when repeated bright-light exposure is combined with a shifted sleep-wake schedule. Dijk et al^72^, for example, advanced the sleep-wake schedule by 10 h over five days and administered approximately 10,000 lx for 5-8 h after each sleep episode. The melatonin rhythm advanced by 8.1 h, whereas the same sleep schedule under approximately 90-150 lx produced no significant advance. Robust advances have likewise been observed in naturalistic studies that altered the entire light-dark pattern over several days. After one week of camping under natural light, during which mean waking illuminance increased from 979 to 4,487 lx, melatonin phase markers occurred approximately 2 h earlier^4^. Even a two-day camping weekend advanced melatonin onset by 1.4 h, whereas it was delayed by 0.98 h among participants who remained in the modern electrical-lighting environment^5^. In a home setting, one week of greater daylight access through electrochromic glazing maintained DLMO timing, whereas one week with conventional blinds produced a 15-min delay and lower positive affect^29^. These findings do not imply that a single day cannot change circadian phase, as appropriately timed high-intensity pulses clearly can. Rather, they suggest that, at the moderate intensity of our protocol and with sleep-wake timing and evening light held constant, one day of additional daytime light may have been insufficient to produce a consistent change in DLMO, and repeated changes in the daily light-dark pattern may be more relevant.

Besides light, physical activity can act as a non-photic zeitgeber whose phase-shifting effect likewise depends on timing. The phase shifts reported for exercise, however, were elicited by more intense and repeated stimuli than ours. Three consecutive days of 1-h treadmill exercise at 65-75% of heart rate reserve advanced the onset of urinary 6-sulphatoxymelatonin by approximately 0.7-0.9 h when performed at 7:00, 13:00, or 16:00^34^, and five days of 30-min morning exercise at 70% of peak oxygen uptake advanced DLMO by 0.62 h^35^. In comparison, median heart rate reserve in our active group averaged 32.66% (*SD* = 7.67) during approximately 4 h of walking on a single day, well below the relative intensity used by Youngstedt et al^34^. In our protocol, activity was concentrated from late morning to mid-afternoon and thus overlapped with the afternoon advance region of the exercise phase-response curve. Consistent with this timing, higher activity was associated with an earlier DLMO (*b* = −6.48 min per *SD*, 95% HDI [−13.79, 1.27], 34.5% in ROPE, *BF*₁₀ = 1.56), even though the evidence remained inconclusive. Given the coincidence of light and activity within this advancing window, we expected them to reinforce one another. However, the interaction was inconclusive as well (*b* = 2.15 min, 95% HDI [−4.50, 8.96], 77.5% in ROPE, *BF*₁₀ = 0.39). Laboratory evidence on this combination is similarly mixed. Adding exercise to a late-night bright-light stimulus did not significantly enhance the phase delay, although that study was powered to only detect differences of approximately 2-2.5 h^37^, whereas bright light followed by early-morning exercise produced additive phase delays in another^38^. Both protocols examined phase delays rather than the advances relevant to our exposure window and used electric light of considerably higher intensity, which limits their comparability with our naturalistic exposure. Detecting an interaction of plausible size may therefore require larger samples or stronger stimuli than any of these studies provided.

Among the exploratory secondary outcomes, only daytime mood showed evidence of an association with light. Higher light exposure was associated with slightly better mood (*b* = 0.14 per *SD*, 95% HDI [0.02, 0.26], 99.5% in ROPE, *BF*₁₀ = 3.32); with nearly the entire posterior inside the ROPE, this association was practically negligible per *SD*. Between high and low light exposure, the difference amounted to roughly a third of a point on the five-point scale (0.28, 95% HDI [0.04, 0.52]). The association between daytime light and evening sleepiness remained inconclusive, even though the HDI just excluded zero (*b* = 0.25 per *SD*, 95% HDI [0.03, 0.45], 99.0% in ROPE, *BF*₁₀ = 2.76). Most remaining associations of light and activity with subjective ratings were inconclusive, while a few provided moderate evidence against a practically relevant effect (Tables 4 and 5). For evening psychomotor vigilance, the data provided evidence against a practically relevant association with daytime light and, once baseline performance was accounted for, with activity. This finding is consistent with reviews concluding that the acute alerting effects of light, though well-established at night, remain inconsistent and weak during the day in rested individuals^73,74^, which seems in line with our mostly inconclusive findings. Collectively, none of the secondary outcomes provided evidence of a practically relevant effect of a single day of natural daylight or physical activity.

### Limitations and Future Directions

Several limitations should be considered when interpreting these results, and they point to priorities for future research. First, our sample comprised healthy young adults on regular sleep-wake schedules. Living in and around Basel, they likely have substantial exposure to daylight through active, outdoor daily travel. Prior light history modulates the response to subsequent light, with a dimmer history enhancing both melatonin suppression^6^ and phase shifts^7^. Against a potentially high baseline, and given the considerable interindividual variability in the change in DLMO, a single day of modest additional daylight may thus have been insufficient to elicit a discernible change. Future studies should therefore systematically vary exposures to establish which intensity, contrast, and timing may change circadian phase most effectively, while keeping sleep-wake schedules constant. Such work would complement the extensive literature on electrical lighting. A further limitation was that, at study onset, no practical means of capturing the spectral composition of light at eye level was available. The ActTrust 2 provided photopic illuminance only and did not permit direct estimation of melanopic or other α-opic exposure. Incorporating spectral information is our main recommendation for future work, as this would enable the melanopic effects we had hoped to isolate through the glasses condition to be quantified directly. Our analyses were further constrained by recurrent failures in heart-rate recordings, which substantially reduced the amount of usable activity data. Finally, the naturalistic outdoor setting introduced variability that could not be controlled experimentally, including approximately 2 h of car travel within the outdoor window. At the same time, we consider this setting a clear strength, as much of the evidence on the effects of light and activity stems from highly controlled settings with electrical lighting, whereas studies combining real-world daylight exposure with fixed sleep-wake schedules and controlled activity remain rare. Such designs are essential if findings are to be interpretable and applicable in everyday life. Ultimately, real-world light exposure is shaped as much by behaviour (i.e., when and where people spend their time) as by the physical environment^75^. Characterising this interplay will therefore be central to translating circadian research into everyday health recommendations.

### Conclusion

To our knowledge, this is the first study to examine a single day of naturalistic daylight, combined with physical activity, under this degree of experimental control. In healthy young adults on sleep-wake schedules fixed to their habitual timing, one controlled day of outdoor daylight and activity provided no conclusive evidence of a change in circadian phase. Among the exploratory outcomes, only the association between higher light exposure and better daytime mood received moderate support, and its magnitude was practically negligible. Critically, these inconclusive findings should not be interpreted as evidence that daylight or physical activity has no effect on circadian timing or circadian health, nor do they contradict previous findings. Instead, they indicate that a single day of moderate exposure may be insufficient to elicit measurable effects under the present conditions, and that repeated or habitual daylight exposure may be more relevant^26^. Taking prior research on the health effects of light into account, we suggest that regular daylight exposure, ideally soon after waking, when light is more likely to advance circadian phase, may better support circadian health than a single day of exposure.

## Acknowledgements

We thank all participants for their time and commitment to the study. Furthermore, we are deeply grateful to the involved research interns, study helpers, and master students for their dedicated support with organisation, recruitment, data collection, and day-to-day study operations. Thank you, Béla Bernasconi, Cyrill Bühler, Irina Capatina, Patrick Carigiet, Melisa Cimendag, Samra Dilji, Emilia Essl, Michelle Fischer, Niklas Fischer, Rebecca Frei, Lina Fricke, Jasmin Fritschi, Alexandra von Gatterburg, Sarah Graf, Lena Kabaran, Saskia Knecht, Flavia Krüttli, Kim Meier, Ivona Mikljanová, Laura Mongili, Ronja Nidecker, Ellen Reichert, Julia Saner, Korben Schlier, Leia Schöllkopf, Melanie Steinmetz, Manuela Thommen, Lisa Tran, Maria Vettiger, Andrea Virag, Jannis Wieczorek, Moesha Zelger, and Filiz Zopfi. We further thank Roland Steiner (Department of Physics, University of Basel, Switzerland) for measuring the spectral transmittance of the glasses used in the study. Finally, Ann-Sophie thanks Gerhard Sachs for his invaluable technical assistance in setting up and troubleshooting the PsychoPy test battery.

## Author contributions

ASL: Methodology, Investigation, Data curation, Formal analysis, Software, Visualization, Project administration, Supervision, Writing – original draft, Writing – review & editing. MM: Investigation, Formal analysis, Software, Writing – review & editing. FS: Methodology, Investigation, Writing – review & editing. CG: Investigation. AS: Methodology, Writing – review & editing. CB: Conceptualization, Funding acquisition, Supervision, Writing – review & editing.

## Data availability

Analysis code and de-identified derived data supporting the findings will be made publicly available on Zenodo upon publication (reserved DOI: 10.5281/zenodo.23809173). During the peer review, the draft deposit is accessible to editors and reviewers through a private read-only link provided with the submission.

## Ethics declarations

The study was approved by the Ethics Committee of Northwestern and Central Switzerland (EKNZ; approval number 2022-01140) and conducted in accordance with the Declaration of Helsinki and Swiss law.

## Declaration of generative AI and AI-assisted technologies in the writing process

During the preparation of this manuscript, the authors used ChatGPT (OpenAI, chatgpt.com) and Claude (Anthropic, claude.ai) to support coding tasks (i.e., troubleshooting and plotting) and for language polishing, including improving clarity, coherence, and wording. All outputs were critically reviewed and edited by the authors, who take full responsibility for the content of the publication.

