## Supplementary Material for "A single day of natural daylight exposure and physical activity does not conclusively change circadian phase: A Bayesian crossover experiment"

Results S8. Analysis Samples, Sampling Diagnostics, and Posterior Predictive Check . 55

**References** 58

Figure S9 Posterior Predictive Check of the Primary Model for the Change in DLMO 57

#### List of Tables

| | | |
|---|---|---|
| Table S1 | Exclusion Criteria, Instruments, and Stage of Assessment | 5 |
| Table S2 | Spot Measurements of Laboratory Lighting at Eye Level | 10 |
| Table S3 | Regions of Practical Equivalence and Coefficient Prior Scales by Outcome | 21 |
| Table S4 | Manipulation Checks and Baseline Equivalence Across Light Conditions and Activity Groups | 27 |
| Table S5 | Posterior Estimates for the Change in DLMO: Primary Model and Sensitivity Analyses | 31 |
| Table S6 | Derived Contrasts for the Change in DLMO: Primary Model and Sensitivity Analyses | 33 |
| Table S7 | Change in DLMO by Light Condition and Activity Group: Categorical Model | 35 |
| Table S8 | Posterior Estimates and Derived Contrasts for the Change in Evening Melatonin AUC (Exploratory) | 36 |
| Table S9 | Posterior Estimates for Evening Time-Course Outcomes | 38 |
| Table S10 | Derived Contrasts for Evening Time-Course Outcomes | 41 |
| Table S11 | Posterior Estimates for Baseline-Adjusted Psychomotor Vigilance Outcomes (Model M1) | 44 |
| Table S12 | Light Exposure and Physical Activity Coefficients Across Baseline-Adjustment Models for Psychomotor Vigilance | 46 |
| Table S13 | Posterior Estimates for Daytime Time-Course Outcomes | 49 |
| Table S14 | Derived Contrasts for Daytime Time-Course Outcomes | 50 |
| Table S15 | Leave-One-Out Cross-Validation Comparisons of Candidate Models | 53 |
| Table S16 | Analysis Samples and Sampling Diagnostics of All Reported Models | 56 |

#### Supplementary Method

##### Method S1. Eligibility Criteria

Eligibility was assessed in several stages: an online screening with an extensive screening survey, an adaptation night with polysomnography (PSG) and a medical examination by the study physician, and adherence and drug checks at every experimental visit. Table S1 lists all exclusion criteria together with the instruments and cut-offs that were applied and the stage at which each criterion was assessed.

**Table S1**

*Exclusion Criteria, Instruments, and Stage of Assessment*

| Domain | Instrument or assessment | Exclusion criterion | Stage |
|---|---|---|---|
| Ability to understand study materials | — | Inability to understand study materials | Pre-screening |
| Ability to understand and/or follow procedures | — | Inability to understand and/or follow procedures | Pre-screening |
| Previous study participation | — | Prior participation in this study | Pre-screening |
| Age | — | <18 or >35 years | Screening survey |
| Body weight | BMI calculated from self-reported height and weight | BMI < 18.5 or > 24.9 kg/m$^2$ (outside the WHO normal-weight range) | Screening survey |
| Physical fitness | IPAQ; Self-report on frequency and type of training | No moderate endurance training or fitness (i.e., none or extreme) | Screening survey |
| Pregnancy | Self-report | Pregnancy | Screening survey |
| Shift work | Self-report | <3 months prior to study start | Screening survey |
| Transmeridian travel | Self-report | >2 time zones <1 month prior to study start | Screening survey |
| Chronotype | Munich Chronotype Questionnaire (MCTQ) | ≤2 or ≥7 | Screening survey |
| Habitual sleep duration | MCTQ | <6 or > 10 h | Screening survey |

*Table S1 (continued)*

| Domain | Instrument or assessment | Exclusion criterion | Stage |
|---|---|---|---|
| Chronobiological health | Self-report | Any indication of chronobiological disorder (e.g., delayed or advanced sleep phase syndrome) | Screening survey |
| Photosensitive epilepsy | Self-report | Self-reported photosensitive epilepsy | Screening survey |
| Colour vision | Ishihara test | Abnormal colour vision (< 80% correct identification of Ishihara plate numbers) | Screening survey |
| Physical health | Physical examination by study physician | — | Adaptation night |
| Psychiatric health | Physical examination by study physician | — | Adaptation night |
| Medications impacting on visual, neuroendocrine, sleep, and circadian physiology | Physical examination by study physician | — | Adaptation night |
| Normal sleep | Polysomnography (PSG), self-report | Any indication of sleep disorder | Adaptation night |
| Insomnia | PSG; Pittsburgh Sleep Quality Index (PSQI) | Sleep efficiency <70% or PSQI > 5 | Adaptation night |
| Sleep apnoea | PSG | Apnoea index > 10 | Adaptation night |
| Periodic limb movements (PLM) | PSG | PLM index > 15 | Adaptation night |
| Functioning vision | Self-report; physical examination by study physician | Any vision disorder | Adaptation night |
| Adherence to the stabilised sleep-wake schedule during the six days prior to and between the experimental visits | Actimetry; sleep log | Non-adherence = deviation of >30 min from scheduled times more than twice per study week or on any of the two days preceding a study visit. | Every visit |
| Drug use | Drug-Screen-Multi 6 (nal von minden) | Any positive test | Every visit |

*Note.* The screening survey was completed online before the adaptation night. BMI = body mass index; IPAQ = International Physical Activity Questionnaire (Craig et al., 2003); MCTQ = Munich ChronoType Questionnaire (Roenneberg et al., 2003), cut-offs refer to mid-sleep on free days (h); PSQI = Pittsburgh Sleep Quality Index (Buysse et al., 1989); PSG = polysomnography; PLM = periodic limb movements. — = no fixed cut-off.

#### Method S2. Outdoor Hiking Route and Treadmill Protocol

Participants in the active group walked for approximately 4 h in total during the light exposure window, divided into four bouts of approximately 1 h each. In the daylight and glasses conditions, they hiked a fixed outdoor route; in the laboratory condition, they walked on a treadmill programmed to approximate the same route. Inactive participants remained sedentary throughout the corresponding period.

##### *Outdoor Route*

The route was located near Nunningen (Solothurn, Switzerland) and started and ended at the Nunningerberghütte (885 m above sea level; Figure S1). It covered 15.7 km, elevations from 771 to 996 m, and 411 m of both ascent and descent. Three loops of 3.69–4.04 km were arranged in a cloverleaf layout around the hut; Loop 1 was repeated as the first and fourth of four approximately 1-h walking bouts. Returning to the hut after each loop allowed participants to walk without backpacks. The breaks after bouts 1 and 3 lasted 15 min, whereas the break after bout 2 was a 30-min lunch break.

##### *Treadmill Programme*

The outdoor route was converted into a treadmill programme from GPS and elevation data. The route was drawn in AllTrails (www.alltrails.com) and exported as a GPX file, and the corresponding coordinates and elevations were downloaded as a CSV file. In Python (v3.11), consecutive coordinates were converted into route sections with associated distances and gradients; Microsoft Excel (v16.71) was used to tabulate the final programmes. Because the treadmill accepts at most 40 programmable stages, each loop was condensed into 40 segments (mean segment length approximately 98 m), and the mean gradient was calculated for each segment.

**Figure S1**

*Outdoor Hiking Route Near Nunningen, Switzerland*

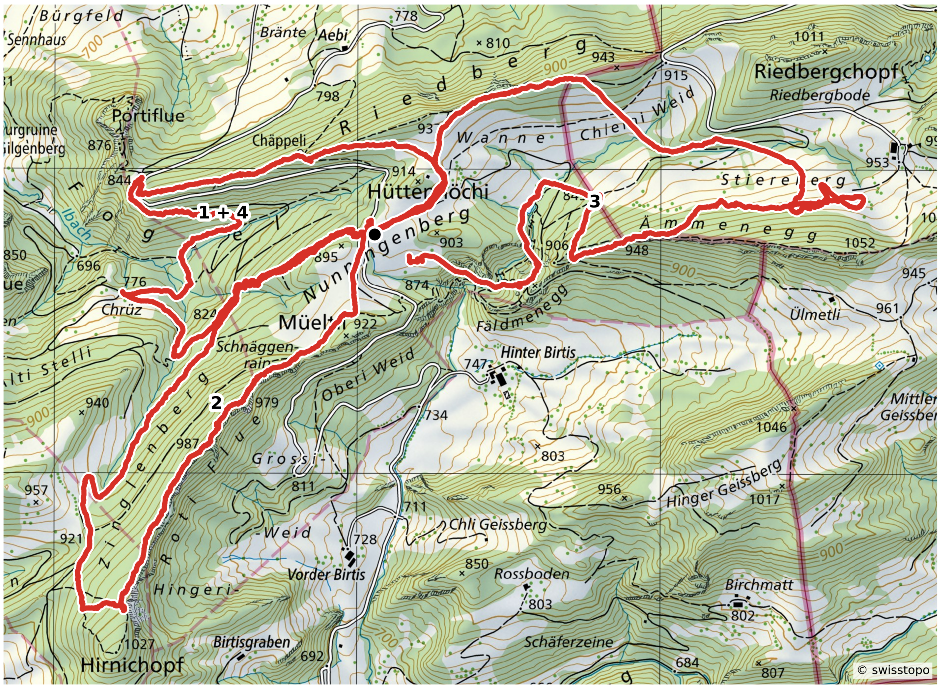

*Note.* The red line shows the hiking route; numbers indicate the order in which the loops were walked. Loop 1 was walked twice, as the first and fourth bout. The black dot marks the start and end point at the Nunningerberghütte. Base map: Federal Office of Topography swisstopo (swisstopo national map; accessed 11 October 2026). Route overlay based on the study GPX records. © swisstopo.

A constant 2-percentage-point offset was added to every segment's outdoor gradient, following the treadmill provider's technical recommendation for comfortable walking; thus, nominally level terrain was programmed at a 2% incline. This was a pragmatic protocol choice rather than an attempt to reproduce the energetic cost of outdoor walking exactly. The gradients were then adapted to the technical limits of the treadmill, which cannot simulate downhill walking and has a maximum incline of 20%: downhill segments were set to 0%, and gradients above 20% were capped at 20%. Inclines were rounded to the nearest 0.5% and distances to the nearest 10 m, and adjacent segments with comparable inclines were merged where possible. The final programmes were entered manually into a pluto med treadmill (h/p/cosmos, Nussdorf-Traunstein, Germany).

On uphill segments, the default treadmill speed was 4.0 km/h and could be adjusted by participants between 3.8 and 4.2 km/h. Because downhill walking could not be simulated, segments representing level or downhill terrain had a faster default speed of 4.5 km/h, adjustable between 4.3 and 4.7 km/h. Participants selected a speed within these ranges that felt brisk but not strenuous; no participant required a speed outside them. The bout and break structure matched that of the outdoor hike.

**Method S3. Laboratory Lighting**

To characterise the experimental lighting, eye-level spot measurements were obtained in the laboratory used by inactive participants with a calibrated spectroradiometer (JETI spectraval 1501, JETI Technische Instrumente GmbH, Jena, Germany) at typical participant positions, with the sensor oriented approximately in the direction of gaze (Figure S2; Table S2). The fluorescent ceiling-lamp panel (Philips Lighting, Eindhoven, The Netherlands) used during the 8-h exposure window provided a median photopic illuminance of 51 lx (range 44–85 lx). The dim background lighting used outside the exposure window, including PSG preparation and both evening test batteries, remained below 20 lx at all measured positions (maximum 16.7 lx). This exceeded the preregistered ceiling of 10 lx at some positions closer to the background light source. These spot measurements described the lighting installation; individual light exposure in all analyses was derived from the ActTrust 2 recordings (Method S4). Measurements were obtained on 27 January 2026, after data collection; the lighting installation and programmed settings in this laboratory were unchanged.

**Figure S2**

*Laboratory Layout and Spot-Measurement Positions*

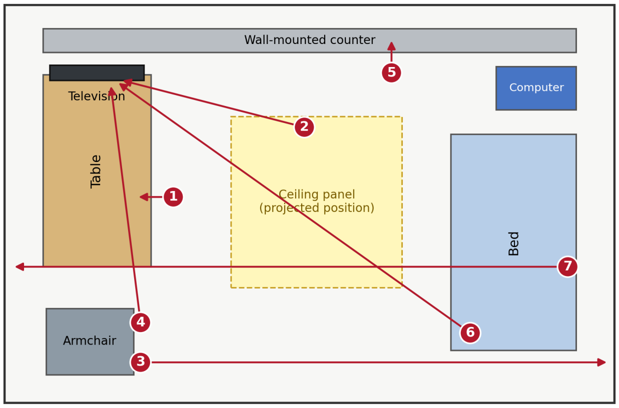

*Note.* The schematic depicts the laboratory used by inactive participants and is not to scale; arrows indicate gaze direction. Positions: 1, table (down); 2, chair towards television; 3, armchair towards opposite wall; 4, armchair towards television; 5, wall-mounted counter (down); 6, bed towards television; 7, bed towards opposite wall. The active-participant laboratory contained a treadmill in place of the table.

**Table S2**

*Spot Measurements of Laboratory Lighting at Eye Level*

| Position | Photopic illuminance (lx) | Melanopic EDI (lx) | CCT (K) |
|---|---|---|---|
| **Laboratory condition: ceiling panel during the exposure window** | | | |
| Chair at table (gaze down) | 65.3 | 26.0 | 2,672 |
| Chair towards television | 64.6 | 28.6 | 2,864 |
| Armchair towards wall | 44.2 | 21.5 | 3,116 |
| Armchair towards television | 85.1 | 39.9 | 3,026 |
| Chair at wall-mounted counter (gaze down) | 50.6 | 24.8 | 3,153 |
| Sitting on bed towards television | 50.8 | 24.4 | 3,093 |
| Sitting on bed towards wall | 46.8 | 22.9 | 3,122 |
| **Dim light: evenings and PSG preparation** | | | |
| Chair towards computer (screen on) | 16.1 | 6.1 | 2,515 |
| Chair towards computer (screen off) | 16.7 | 6.2 | 2,495 |
| Chair at table (gaze down) | 9.6 | 2.9 | 2,178 |
| Chair at counter (gaze down) | 11.0 | 4.0 | 2,453 |
| Sitting on bed towards wall | 6.2 | 2.2 | 2,433 |
| Chair in EEG position | 3.7 | 1.3 | 2,408 |

*Note.* Each row is a single vertical-plane spot measurement at the participant's eye position, taken in the direction of gaze. During the exposure-window measurements, the television showed a nature documentary; for the dim light, the computer screen showed a black slide and all displays were off otherwise. Melanopic EDI = melanopic equivalent daylight (D65) illuminance; CCT = correlated colour temperature.

**Method S4. Light Exposure Processing and Filter Correction**

***Recording and Summary Measures***

Light exposure was recorded at 1-min resolution with an ActTrust 2 (Condor Instruments, São Paulo, Brazil) mounted on the forehead in a custom 3D-printed housing and oriented in approximately the direction of gaze. Recordings were matched to participants and visits, and the minute-level light records were restricted to the recorded start and stop times of each visit's nominal 8-h exposure window. The LIGHT column in the exported sensor data contained photopic illuminance in lux. The primary light measure was the median of the minute-level values within the window, because minute-level illuminance is strongly right-skewed and intermittently dominated by brief extremes.

For the cumulative-light sensitivity analysis (Method S7), the corrected minute-level values were instead summed over the same window; at a 1-min recording interval, this sum approximates exposure in lux-minutes. Among the 117 visits with usable light data, the median number of recorded window minutes was 480 (range 411–525), and 102 visits contained exactly 480 records. These counts reflected the records retained between the visit-specific start and stop times, rather than a duration-normalised 480-min measure. The sum was not rescaled to compensate for missing minutes or differences in the number of retained records; it therefore reflected both illuminance and the number of recorded minutes. Neither the AMB LIGHT channel nor a melanopic proxy was used as a model predictor.

Prior light history was recorded on the day preceding the exposure day with an ActTrust 2 worn on a collar at chest height. Participants were instructed to wear the device from waking until arrival at the laboratory, and filter glasses were not worn. Before waking and after arrival, the device was to be stored in a dark, quiet location and was covered or placed in its case at the laboratory. The pragmatic manipulation-check summaries used all 1,440 uncorrected 1-min photopic illuminance values recorded on that calendar day; records outside the instructed wear interval were therefore retained but were expected to be zero. The summaries were mean photopic illuminance, cumulative photopic exposure, and time above 250 lx and above 1,000 lx. Cumulative exposure was calculated as the sum of the 1-min illuminance values and therefore has units of lux-minutes.

##### ***Filter-Glasses Correction***

In the glasses condition, the filter glasses strongly attenuated short-wavelength light (Figure S3), whereas the forehead-mounted sensor recorded light outside the lenses. A transmission factor specific to the LIGHT channel was therefore estimated from the wavelength-dependent transmission of the filter, $T(\lambda)$, measured at the Department of Physics, University of Basel, and the spectral response of the ActTrust 2 green channel, $G(\lambda)$. Manufacturer documentation confirms that the LIGHT output is a green-channel-derived estimate of photopic illuminance (Condor Instruments, 2024). The green-channel response curve was digitised from the same manufacturer's *ActTrust 2 Light Sensor Description* (revision 1) with WebPlotDigitizer v4.2 (Rohatgi, 2019), interpolated to a 1-nm grid from 380 to 795 nm by locally weighted regression (LOESS; span = 0.1), and aligned with the filter measurements. The filtered and unfiltered response curves were then weighted by the CIE 1924 photopic luminous efficiency function, $V(\lambda)$, obtained from LightLogR v0.10.0 (Zauner et al., 2025), and their ratio defined the transmission factor:

$$T_{\text{LIGHT}} = \frac{\sum_{\lambda=380}^{795} G(\lambda)\,T(\lambda)\,V(\lambda)}{\sum_{\lambda=380}^{795} G(\lambda)\,V(\lambda)} = 0.4038. \tag{S1}$$

Hence, the estimated sensor- and photopic-weighted transmission of the filter for the LIGHT channel was 40.38%. Every minute-level LIGHT value recorded in the glasses condition was multiplied by this factor once, before the visit summary was calculated; no factor was applied in the daylight or laboratory conditions. The corrected value is an operational estimate of the photopic illuminance behind the lenses rather than a direct corneal measurement. Applying one scalar factor across the exposure window assumes stable filter transmission and does not capture variation in the spectral composition of ambient light over time.

**Figure S3**

*Spectral Transmittance of the Short-Wavelength Filter Glasses*

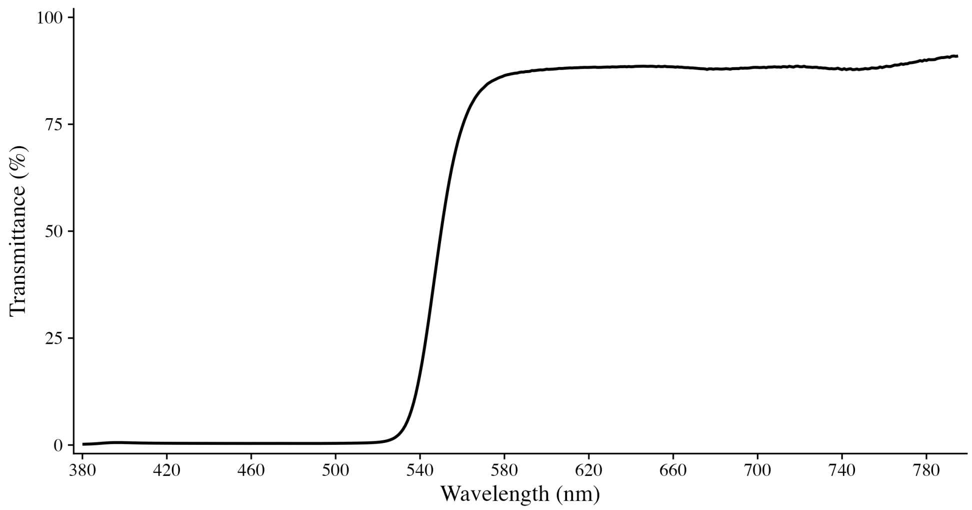

*Note.* The curve shows measured spectral transmittance of the Honeywell UVEX lenses from 380 to 795 nm (1-nm intervals), measured at the Department of Physics, University of Basel. The plotted values are the measured percentage transmittance, without sensor or photopic weighting. The sensor- and photopic-weighted transmission factor used to correct the ActTrust 2 LIGHT values was 0.4038 (40.38%; Method S4).

***Quality Control***

Two screens were applied to the visit-level data. First, statistical outliers were flagged as visit medians outside 1.5 interquartile ranges of the log-transformed values within each light-condition × activity-group cell. Second, plausibility flags marked any raw laboratory reading above 100 lx, any raw outdoor reading at or below 1 lx, and daylight-condition visits without any minute above 250 lx. These flags prompted visual inspection but did not lead to automatic exclusion. Visits with less than 80% recorded coverage between the first and last available timestamp of the exposure window were to be excluded; no visit met this criterion. After inspection, light data were set to missing for two of 119 visits. One daylight visit returned only zero values across the window (device off or technical failure). One laboratory visit showed implausibly high photopic illuminance for the controlled setting (median 925 lx, maximum 13,981 lx; 448 of 480 min above the 100-lx plausibility threshold), indicating a sensor or orientation artefact. All other flagged visits were retained unless a different model variable was missing.

**Method S5. Heart Rate Reserve**

Physical activity was quantified as the percentage of heart rate reserve (HRR), a measure of relative exercise intensity. Heart rate was recorded continuously during the activity period of each exposure day (and during the corresponding sedentary period in the inactive group) with a chest-strap Polar H10 sensor (Polar Electro Oy, Kempele, Finland) connected to the Polar Beat smartphone application. The device recorded at 130 Hz; the exported heart-rate series used for processing contained one value per second in beats per minute (bpm).

Resting heart rate ($HR_{\text{rest}}$) was derived from the ECG channel of the PSG during the 3-min eyes-open resting-state recordings obtained 10 min after waking. Both morning recordings were combined when available; otherwise, the available recording was used. Inter-beat intervals in milliseconds were converted to seconds, and their cumulative sum was used to reconstruct the time axis. The intervals were linearly interpolated to a regular 1-s grid and converted to bpm. Values below 40 or above 110 bpm were recoded as missing, missing values were linearly interpolated, and a centred three-point rolling median reduced short-term noise and artefacts. The processed values were mapped back to the original interval timestamps, and $HR_{\text{rest}}$ was the participant's mean across all available recordings. Maximal heart rate ($HR_{\text{max}}$) was the value in the "Absolute Maximalwerte" column of the screening cardiopulmonary exercise-test output (ramp protocol on a cycle ergometer to volitional exhaustion).

The activity-period heart-rate signal was averaged within 5-min bins. Missing bins were imputed separately for each activity group by predictive mean matching with the mice package (van Buuren & Groothuis-Oudshoorn, 2011), using the participant identifier and binned heart rate as the input variables. Ten completed datasets were generated with five iterations each and a fixed random seed (123). The ten imputed heart-rate values for each bin were averaged to obtain one estimate. No maximum length of consecutive missing bins was imposed; the longest imputed interval was 275 min. The resulting single set of covariate values was used in the models, without propagating between-imputation uncertainty. For each bin, HRR was computed as

$$\text{HRR} = \frac{HR - HR_{\text{rest}}}{HR_{\text{max}} - HR_{\text{rest}}} \times 100, \tag{S2}$$

with negative values set to zero. The median HRR across the activity period served as the physical-activity predictor in all models. Visits with missing or blank heart-rate recordings were excluded from the corresponding models; among visits with an available change in DLMO, this applied to 28 visits (see main text).

#### Method S6. Model Specification, Priors, and Inference

##### *Model Structure*

All outcomes were analysed with Bayesian hierarchical models with Gaussian likelihoods and the same two continuous predictors: light exposure, $L$, the median photopic illuminance across the 8-h exposure window, transformed as $\log_{10}(x + 1)$ and *z*-standardised; and physical activity, $A$, the median HRR across the activity period, likewise *z*-standardised. Both predictors were standardised across all available values of the respective analysis dataset, that is, before model-specific complete-case deletion (one value per visit for the visit-level outcomes; one row per assessment for the time-course outcomes, in which each visit's value was repeated across its assessments). Hence, 1 *SD* refers to the dispersion in that dataset and can differ slightly from the dispersion within an individual model's analysis sample. For outcomes with one observation per visit $v$ of participant $p$ (change in DLMO and change in melatonin AUC), the model was

$$y_{pv} = \beta_0 + \beta_1 L_{pv} + \beta_2 A_{pv} + \beta_3 L_{pv} A_{pv} + u_{0p} + \varepsilon_{pv}. \tag{S3}$$

Evening outcomes were measured at ten assessments $t = 0, \dots, 9$ on Evening-2, and assessment was centred at the midpoint of the series, $s_t = t - 4.5$. The evening models asked whether the within-evening trajectory depended on light exposure or physical activity and therefore included assessment-by-predictor interactions and random intercepts and assessment slopes at both the participant and the visit level:

The visit-specific intercept and assessment slope were defined as

$$\mu_{pv} = \beta_0 + \beta_2 L_{pv} + \beta_3 A_{pv} + \beta_4 L_{pv} A_{pv} + u_{0p} + w_{0pv},$$

$$\gamma_{pv} = \beta_1 + \beta_5 L_{pv} + \beta_6 A_{pv} + u_{1p} + w_{1pv},$$

so that the evening model was

$$y_{pvt} = \mu_{pv} + s_t \gamma_{pv} + \varepsilon_{pvt}. \tag{S4}$$

The daytime models estimated the overall associations during the activity period within the 8-h light-exposure window. This period included approximately 4 h of walking in the active group and corresponding sedentary behaviour in the inactive group. Light exposure and activity occurred while the twelve assessments were collected, but each visit contributed only one summary value for each predictor. The daytime specification therefore estimated associations with these visit-level summaries, while adjusting additively for assessment, rather than estimating responses to concurrent changes in light or activity. Assessment was centred at the midpoint of the series ($t = 0, \ldots, 11$; $s_t = t - 5.5$), with participant-level random intercepts and assessment slopes and a visit-level random intercept:

$$y_{pvt} = \beta_0 + \beta_1 s_t + \beta_2 L_{pv} + \beta_3 A_{pv} + \beta_4 L_{pv} A_{pv} + u_{0p} + u_{1p} s_t + w_{0pv} + \varepsilon_{pvt}. \tag{S5}$$

In all models, random effects were normally distributed with mean zero and estimated as independent components, and $\varepsilon \sim \mathcal{N}(0, \sigma^2)$. Complete-case deletion was applied per model, so that an observation was excluded only when that model's outcome or one of its predictors was missing.

##### *Priors*

Priors were weakly regularising and scaled to $s_y$, the standard deviation of the respective outcome in the model's analysis sample (Table S3). The intercept received a $\mathcal{N}(\bar{y}, 2s_y)$ prior, each regression coefficient a $\mathcal{N}(0, s_y/3)$ prior, random-intercept standard deviations half-normal priors with scale $0.5s_y$, random-slope standard deviations half-normal priors with scale $0.25s_y$, and the residual standard deviation a half-normal prior with scale $s_y$.

##### *Estimation*

Models were fitted in PyMC (Abril-Pla et al., 2023) with the No-U-Turn sampler (Hoffman & Gelman, 2014), using four chains with 10,000 tuning iterations and 6,000 posterior draws each, a target acceptance probability of 0.95, and a maximum tree depth of 15. Convergence was assessed with rank-normalised split-$\hat{R}$, bulk and tail effective sample sizes (ESS, Vehtari et al., 2021), and divergent transitions, computed with ArviZ (Kumar et al., 2019). The Bayesian fraction of missing information (BFMI), an energy diagnostic of the sampler, was additionally calculated from the saved fits for the diagnostic summary; it was not a criterion in the automatic refitting procedure. If a fit showed any divergent transition, a maximum $\hat{R}$ above 1.02, or a minimum ESS below 1,000, the target acceptance probability was increased by 0.025 and the model was refitted, up to three attempts. Table S16 reports the analysis samples and sampling diagnostics; for the categorical DLMO model, the available diagnostics were taken from the saved notebook output. Posterior predictive checks were performed for all outcomes; the check for the primary DLMO model is shown as an example (Figure S9).

***Inference***

Posterior distributions were summarised by their mean and 95% highest-density interval (HDI). Three indices were reported for each effect. The probability of direction (*pd*) is the posterior probability that the effect has the sign of its posterior mean. The proportion of the posterior inside an outcome-specific region of practical equivalence (ROPE) indexes practical relevance (Kruschke, 2018); the ROPE bounds were specified a priori for each outcome (Table S3), and the percentage of the full posterior inside the ROPE was reported. Because both predictors were *z*-standardised, the light exposure and physical activity coefficients are outcome differences per 1 *SD*, and the light × activity coefficient, the change in the light exposure coefficient per 1-*SD* increase in physical activity, is likewise an outcome difference (a difference between two outcome differences). The outcome-specific ROPE was applied to all three fitted coefficients and to derived contrasts that are direct outcome differences. It was not applied to difference-in-differences or time-course moderation contrasts, for which no separate practical-equivalence threshold was prespecified. The ROPE was not applied to intercepts, assessment terms and their interactions, which are rates per assessment, adjustment terms, or variance parameters. Bayes factors ($BF_{10}$) for the model coefficients were computed with the Savage–Dickey density ratio against a point null at zero (Wagenmakers et al., 2010), using the coefficient's prior in the fitted model. For verbal interpretation, $BF_{10}$ values of 1–3, 3–10, 10–30, and above 30 were described as anecdotal, moderate, strong, and very strong evidence for a nonzero effect, respectively; reciprocal values described evidence for the point null (Jeffreys, 1961; Lee & Wagenmakers, 2014). An effect was described as evidenced when its 95% HDI excluded zero, most of its posterior lay outside the ROPE, and $BF_{10}$ was at least 3. An effect was described as practically negligible when its posterior was concentrated within the ROPE, and as inconclusive when $1/3 < BF_{10} < 3$. Evidence against a practically relevant effect required both $BF_{10} \leq 1/3$ and posterior concentration within the ROPE; an HDI that included zero was not by itself interpreted as evidence of absence.

Derived contrasts translate the coefficients into differences between high (+1 *SD*) and low (−1 *SD*) predictor values, that is, a 2-*SD* difference, with the other predictor held at its mean. Simple effects evaluate the light contrast at high or low physical activity, and the interaction contrast is the difference between these two simple effects. For the time-course outcomes, contrasts are additionally expressed as the predicted change from the first to the final assessment and, for the evening outcomes, evaluated at the first assessment, the midpoint, and the final assessment. Derived contrasts are reported with posterior means, 95% HDIs, *pd*, and, where applicable, ROPE percentages, but without Bayes factors. For contrasts that rescale a single coefficient, the Bayes factor is identical to that of the coefficient. For contrasts that combine several coefficients, a Savage–Dickey Bayes factor would have to use the prior implied for the linear combination by the independent coefficient priors. This implied prior was not chosen for the contrast and becomes considerably wider when terms with large weights enter, such as the assessment × light term at the first or final assessment; the resulting Bayes factors would therefore favour the null hypothesis increasingly and would not be comparable across contrasts. Bayes factors were therefore omitted for derived contrasts; inference for these contrasts was based on the posterior summaries and, where applicable, ROPE percentages.

Candidate fixed-effect specifications were compared with Pareto-smoothed importance-sampling leave-one-out cross-validation (PSIS-LOO, Vehtari et al., 2017), keeping the random-effects structure and the complete-case sample constant within each outcome. For the change in

DLMO and melatonin AUC, the candidates were a null model (intercept only), light only, activity only, an additive model, and the full interaction model. For the daytime outcomes, they were a null model (intercept only), assessment only, an additive model with assessment, light, and activity, and the full model; the random intercepts and assessment slopes were retained in all of them. For the baseline-adjusted vigilance outcomes, models M0 to M3 (Method S8) were compared. A model was judged predictively superior only when the difference in expected log pointwise predictive density (ΔELPD) exceeded approximately twice its standard error, and comparisons in which any model had a Pareto $\hat{k}$ above 0.7 were treated as unreliable (Table S15). Because the reported evening models already contained such observations, no comparison of evening specifications was interpreted (Results S7).

**Table S3**

*Regions of Practical Equivalence and Coefficient Prior Scales by Outcome*

| Outcome | Units | ROPE | Coefficient prior *SD* |
|---|---|---|---|
| **Primary** | | | |
| Change in DLMO | min | ±5 | 10.759 |
| **Exploratory** | | | |
| Change in melatonin AUC, $\log_{10}$(Evening-2/Evening-1) | $\log_{10}$ units | ±0.041 | 0.0456 |
| **Evening time course** | | | |
| Sleepiness (KSS) | KSS points | ±0.5 | 0.600 |
| Mood | scale points | ±0.3 | 0.227 |
| Wellbeing | scale points | ±0.3 | 0.249 |
| PVT median reaction time | ms | ±20 | 19.781 |
| PVT mean response speed | $s^{-1}$ | ±0.06 | 0.1977 |
| PVT slowest-10% response speed | $s^{-1}$ | ±0.05 | 0.1628 |

*Table S3 (continued)*

| Outcome | Units | ROPE | Coefficient prior *SD* |
|---|---|---|---|
| **Daytime time course** | | | |
| Sleepiness (KSS) | KSS points | ±0.5 | 0.449 |
| Mood | scale points | ±0.3 | 0.265 |
| Wellbeing | scale points | ±0.3 | 0.265 |
| Perceived exertion (Borg) | Borg points | ±1 | 0.654 |

*Note.* ROPE bounds were specified a priori on the outcome scale and were applied to fitted light exposure, physical activity, and light × activity coefficients and to derived contrasts that are direct outcome differences; they were not applied to difference-in-differences or time-course moderation contrasts (Method S6). For the change in melatonin AUC, ±0.041 $\log_{10}$ units corresponds to an Evening-2/Evening-1 AUC ratio between approximately 0.91 and 1.10. Coefficient prior *SD* = $s_y/3$, where $s_y$ is the outcome *SD* in the model's analysis sample; all regression coefficients of a model shared this prior. Further priors: intercept $\mathcal{N}(\bar{y}, 2s_y)$; random-intercept *SD*s half-normal($0.5s_y$); random-slope *SD*s half-normal($0.25s_y$); residual *SD* half-normal($s_y$). KSS = Karolinska Sleepiness Scale; PVT = psychomotor vigilance task; AUC = area under the curve.

#### Method S7. Sensitivity Analyses of the Change in DLMO

The primary model (Equation S3) used the change in DLMO (Evening-2 minus Evening-1, in minutes) as the outcome, with the priors described in Method S6. Each sensitivity analysis changed one aspect of this model and retained all others:

1. **Residual distribution.** A Student's *t* likelihood replaced the Gaussian likelihood, with degrees of freedom $\nu \sim \text{Gamma}(\text{shape} = 2, \text{rate} = 0.1)$.
2. **Coefficient priors.** The prior standard deviation of all regression coefficients was widened from $s_y/3$ to $0.5s_y$; the intercept and variance priors were unchanged.
3. **Cumulative light exposure.** The sum of the corrected minute-level photopic illuminance values over the exposure window (Method S4) replaced the visit median. It was transformed as $\log_{10}(x + 1)$ and *z*-standardised in the same way as the primary light measure (Method S6).

4. **Laboratory light values.** For laboratory visits in which raw light readings exceeded 100 lx in at least 25% of the recorded minutes, individual minutes above 200 lx were replaced with 85 lx, the highest eye-level illuminance measured with the spectroradiometer in the laboratory condition (Table S2), and the visit median was recalculated. All other visits and the primary technical exclusions were left unchanged.
5. **Demographic adjustment.** Age ($z$-standardised across participants, one value per participant) and sex (coded −0.5 for male and +0.5 for female) were added as main effects, without interactions.
6. **Categorical exposure.** Light condition and pseudo-randomly allocated activity group replaced the two continuous predictors. Laboratory light and the inactive group served as reference levels; indicators for the daylight and glasses conditions were each interacted with activity group. Activity allocation was matched to the allocation schedule or to the group of an existing participant on the same assessment date to pair active and inactive participants. This model was used descriptively and retained the Gaussian likelihood, participant random intercept, and primary prior scales.

The continuous-predictor variants used complete cases for the outcome and the predictors of the respective variant, and all variants used the sampler settings and convergence criteria described in Method S6.

**Method S8. Baseline-Adjusted Psychomotor Vigilance Models**

Because the activity groups already differed in psychomotor vigilance on Evening-1, before any manipulation (Table S4), the three PVT outcomes (median reaction time, mean response speed, and response speed of the slowest 10% of responses) were additionally analysed with baseline adjustment. The preregistered slowest-decile outcome was retained here as supplementary reporting alongside the two PVT outcomes reported in the main text. For each participant and visit, the baseline was the median of the respective outcome across the available Evening-1 assessments, and it was attached to all Evening-2 assessments of that visit. Evening-2 values were analysed by covariance adjustment rather than as change scores (Vickers & Altman, 2001).

Four models were fitted to the same complete-case observations of each outcome. All of them retained the terms of the evening model (Equation S4), its random-effects structure, and its priors, with $s_y$ computed from the Evening-2 values in the outcome's complete-case sample. Baseline terms were entered as main effects only, without interactions with assessment or the predictors, and received a wider $\mathcal{N}(0, s_y)$ prior to accommodate the strong association between baseline and outcome. The models differed only in how the baseline and physical activity entered:

- **M0** contained no baseline term. It corresponded to the evening model, fitted to the observations with an available baseline.
- **M1** added the visit-level baseline as a single covariate, centred at the grand mean and divided by the *SD* of the visit-level baseline. M1 was the baseline-adjusted model reported in the main text.
- **M2** separated the baseline into a between-participant component, the participant's mean

baseline minus the grand mean, and a within-participant component, the visit baseline minus the participant's mean (Curran & Bauer, 2011). Both components were divided by the *SD* of the visit-level baseline, so that M1 is the special case in which both components share one coefficient.

- **M3** extended M2 by additionally separating physical activity into a between-participant component, the participant's mean HRR minus the grand mean, and a within-participant component, the visit's deviation from the participant's mean. Both components were divided by the *SD* of the HRR used for the standardised predictor, and the within-participant component, which reflected visit-to-visit variation in achieved HRR rather than the between-participant activity allocation, replaced physical activity in the light × activity and assessment × activity terms. The between-participant component, which is observational, entered as an additional main effect.

Comparing M0 with M1 to M3 showed whether the activity association depends on the adjustment for pre-existing differences in vigilance, and M3 isolated the part of the activity association that varies within participants across visits.

### Supplementary Results

#### Results S1. Manipulation Checks and Baseline Equivalence

All PVT metrics were retained in the Evening-1 baseline checks (Table S4). The manipulations produced the intended contrasts (Figure S4; Table S4). Median HRR was markedly higher in the active than in the inactive group, and median photopic illuminance differed strongly across the three light conditions. Prior light history, in contrast, was similar across light conditions, with Bayes factors indicating moderate evidence against a difference. Between the activity groups, the evidence on prior light history was inconclusive: mean illuminance and cumulative exposure did not differ at $p < .05$, whereas time above 250 lx and above 1,000 lx yielded $p$ values below .05 (uncorrected for multiplicity), and all four Bayes factors fell between 1/3 and 3. Figure S5 shows mean photopic illuminance and HRR over time during the light-exposure window and activity period, respectively.

**Figure S4**

*Physical Activity, Light Exposure, and Prior Light History by Group and Condition*

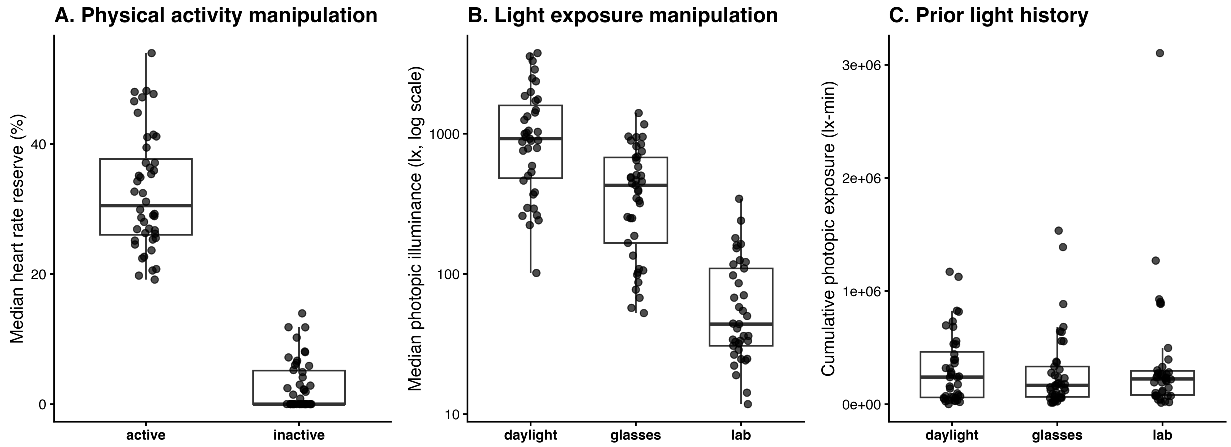

*Note.* (A) Visit-level median heart rate reserve in the inactive and active groups. (B) Visit-level median photopic illuminance during the 8-h exposure window by light condition (log scale); values in the glasses condition are corrected for filter transmission (Method S4). (C) Cumulative photopic exposure on the day before the exposure day, calculated as the sum of the 1-min illuminance values (lux-minutes). Statistical tests are reported in Table S4.

**Table S4**

*Manipulation Checks and Baseline Equivalence Across Light Conditions and Activity Groups*

| Variable | Comparison | *n* | Test statistic | *p* | $BF_{10}$ |
|---|---|---|---|---|---|
| **Exposure day: manipulation** | | | | | |
| Median heart rate reserve | Activity groups | 37 | $W = 342.0$ | < .001 | > 1000 |
| Median heart rate reserve | Light conditions | 20 | $\chi^2(2) = 3.91$ | .142 | 0.17 |
| Median photopic illuminance | Light conditions | 36 | $\chi^2(2) = 46.89$ | < .001 | > 1000 |
| **Descriptive DLMO-change checks** | | | | | |
| Change in DLMO | Light conditions | 31 | $\chi^2(2) = 3.94$ | .140 | 0.69 |
| Change in DLMO | Activity groups | 41 | $W = 148.0$ | .109 | 0.95 |
| **Exposure day: subjective ratings** | | | | | |
| Sleepiness (KSS) | Activity groups | 41 | $W = 186.5$ | .548 | 0.31 |
| Sleepiness (KSS) | Light conditions | 38 | $\chi^2(2) = 8.93$ | .011 | 0.81 |
| Mood | Activity groups | 41 | $W = 217.5$ | .855 | 0.31 |
| Mood | Light conditions | 37 | $\chi^2(2) = 8.38$ | .015 | 7.41 |
| Wellbeing | Activity groups | 41 | $W = 228.0$ | .648 | 0.34 |
| Wellbeing | Light conditions | 37 | $\chi^2(2) = 2.31$ | .314 | 0.96 |
| Perceived exertion (Borg) | Activity groups | 41 | $W = 420.0$ | < .001 | > 1000 |
| Perceived exertion (Borg) | Light conditions | 37 | $\chi^2(2) = 10.26$ | .006 | 3.99 |
| **Prior light history** | | | | | |
| Mean photopic illuminance | Light conditions | 38 | $\chi^2(2) = 1.00$ | .607 | 0.16 |
| Mean photopic illuminance | Activity groups | 41 | $W = 156.0$ | .165 | 0.59 |
| Cumulative photopic exposure (lux-minutes) | Light conditions | 38 | $\chi^2(2) = 1.00$ | .607 | 0.16 |
| Cumulative photopic exposure (lux-minutes) | Activity groups | 41 | $W = 156.0$ | .165 | 0.59 |
| Time above 250 lx | Activity groups | 41 | $W = 124.0$ | .026 | 1.03 |
| Time above 1,000 lx | Activity groups | 41 | $W = 132.0$ | .043 | 0.68 |
| **Evening-1 baseline: circadian phase and ratings** | | | | | |
| DLMO (clock time) | Light conditions | 33 | $\chi^2(2) = 0.32$ | .852 | 0.09 |
| DLMO (clock time) | Activity groups | 41 | $W = 217.0$ | .867 | 0.31 |
| Sleepiness (KSS) | Light conditions | 36 | $\chi^2(2) = 2.76$ | .252 | 0.23 |
| Sleepiness (KSS) | Activity groups | 41 | $W = 173.0$ | .341 | 0.32 |
| Mood | Light conditions | 36 | $\chi^2(2) = 1.48$ | .477 | 0.13 |
| Mood | Activity groups | 41 | $W = 211.0$ | .990 | 0.31 |
| Wellbeing | Light conditions | 36 | $\chi^2(2) = 0.68$ | .710 | 0.10 |
| Wellbeing | Activity groups | 41 | $W = 236.5$ | .497 | 0.33 |
| **Evening-1 baseline: psychomotor vigilance** | | | | | |
| PVT median reaction time | Light conditions | 36 | $\chi^2(2) = 6.00$ | .050 | 0.28 |
| PVT median reaction time | Activity groups | 41 | $W = 30.0$ | < .001 | > 1000 |
| PVT mean reaction time | Light conditions | 36 | $\chi^2(2) = 5.06$ | .080 | 0.35 |

*Table S4 (continued)*

| Variable | Comparison | $n$ | Test statistic | $p$ | $BF_{10}$ |
|---|---|---|---|---|---|
| PVT mean reaction time | Activity groups | 41 | $W = 37.0$ | < .001 | > 1000 |
| PVT fastest-10% reaction time | Light conditions | 36 | $\chi^2(2) = 3.17$ | .205 | 0.31 |
| PVT fastest-10% reaction time | Activity groups | 41 | $W = 22.0$ | < .001 | > 1000 |
| PVT slowest-10% reaction time | Light conditions | 36 | $\chi^2(2) = 0.39$ | .823 | 0.23 |
| PVT slowest-10% reaction time | Activity groups | 41 | $W = 84.0$ | < .001 | 47.24 |
| PVT mean response speed | Light conditions | 36 | $\chi^2(2) = 4.39$ | .111 | 0.18 |
| PVT mean response speed | Activity groups | 41 | $W = 394.0$ | < .001 | > 1000 |
| PVT slowest-10% response speed | Light conditions | 36 | $\chi^2(2) = 0.72$ | .697 | 0.14 |
| PVT slowest-10% response speed | Activity groups | 41 | $W = 354.0$ | < .001 | 378.53 |
| PVT lapses (≥ 500 ms) | Light conditions | 36 | $\chi^2(2) = 0.47$ | .791 | 0.19 |
| PVT lapses (≥ 500 ms) | Activity groups | 41 | $W = 85.5$ | .001 | 14.96 |
| PVT lapses (≥ 400 ms) | Light conditions | 36 | $\chi^2(2) = 3.40$ | .183 | 0.47 |
| PVT lapses (≥ 400 ms) | Activity groups | 41 | $W = 55.0$ | < .001 | > 1000 |
| **Pairwise light-condition comparisons (Holm-corrected)** | | | | | |
| Photopic illuminance | D vs. G | 36 | $V = 595.0$ | < .001 | 359.59 |
| Photopic illuminance | D vs. L | 36 | $V = 663.0$ | < .001 | > 1000 |
| Photopic illuminance | G vs. L | 36 | $V = 637.0$ | < .001 | > 1000 |
| Sleepiness (KSS) | D vs. G | 38 | $V = 240.0$ | .330 | 0.32 |
| Sleepiness (KSS) | D vs. L | 38 | $V = 161.5$ | .022 | 4.07 |
| Sleepiness (KSS) | G vs. L | 38 | $V = 203.0$ | .051 | 0.31 |
| Mood | D vs. G | 37 | $V = 358.5$ | .257 | 0.74 |
| Mood | D vs. L | 37 | $V = 369.0$ | .003 | 18.02 |
| Mood | G vs. L | 37 | $V = 326.0$ | .257 | 0.59 |
| Wellbeing | D vs. G | 37 | $V = 293.5$ | .598 | 0.34 |
| Wellbeing | D vs. L | 37 | $V = 380.5$ | .090 | 2.04 |
| Wellbeing | G vs. L | 37 | $V = 301.5$ | .598 | 0.39 |
| Perceived exertion (Borg) | D vs. G | 37 | $V = 106.5$ | .768 | 0.19 |
| Perceived exertion (Borg) | D vs. L | 37 | $V = 47.5$ | .022 | 1.42 |
| Perceived exertion (Borg) | G vs. L | 37 | $V = 23.0$ | .007 | 6.84 |

*Table S4 (continued)*

| Variable | Comparison | *n* | Test statistic | *p* | $BF_{10}$ |
|---|---|---|---|---|---|
| Median HRR | D vs. G | 20 | *V* = 103.0 | .655 | 0.29 |
| Median HRR | D vs. L | 20 | *V* = 81.0 | 1.000 | 0.29 |
| Median HRR | G vs. L | 20 | *V* = 63.0 | 1.000 | 0.23 |

*Note.* Light conditions were compared with Friedman tests and activity groups with Wilcoxon rank-sum tests; *n* = number of participants entering the test (complete cases across conditions for Friedman tests). $BF_{10}$ values are from the parametric Bayesian counterparts, not Bayes factors for the rank tests: BayesFactor::anovaBF with participant as a random effect for the within-participant light-condition comparisons, and BayesFactor::ttestBF for the between-group comparisons. Pairwise light-condition contrasts used paired Wilcoxon signed-rank tests (*V*) and paired BayesFactor::ttestBF tests. Bayes factors above 1000 are shown as > 1000. Omnibus and activity-group *p* values are uncorrected; pairwise *p* values are Holm-corrected within each set of three light-condition contrasts. *n* denotes complete participant pairs in the pairwise block. Rounded exported zeros are displayed as < .001. DLMO-change tests are descriptive checks; inference on DLMO change was based on the hierarchical models. Prior-light-history summaries used the 1,440 one-minute photopic illuminance values recorded on the calendar day before the exposure day (Method S4). Evening-1 values are participant-level summaries of the baseline evening, before any manipulation. D = daylight; G = filter glasses; L = laboratory light. DLMO = dim-light melatonin onset; KSS = Karolinska Sleepiness Scale; PVT = psychomotor vigilance task.

#### Figure S5

*Time Course of Light Exposure and Heart Rate Reserve During the Exposure Window*

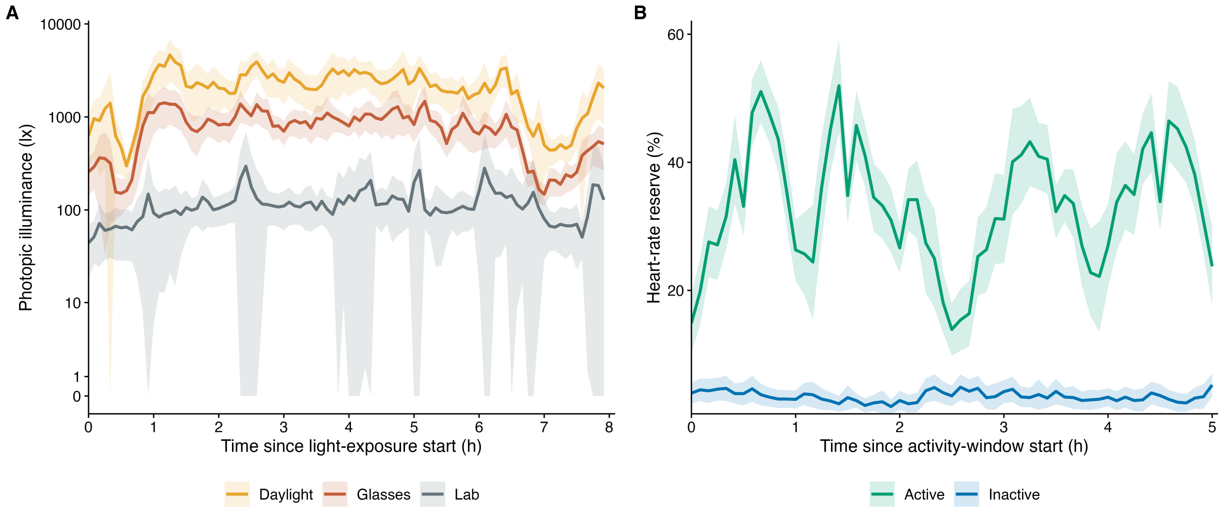

*Note.* (A) Photopic illuminance from the start of the 8-h light-exposure window by light condition (axis that is approximately linear near zero and logarithmic at higher values); values in the glasses condition are corrected for filter transmission. (B) Heart rate reserve from the start of the activity window by activity group. Lines show arithmetic means and shaded bands show pointwise 95% *t* confidence intervals. Panel A was summarised across participant-visits after averaging the 1-min observations within 5-min bins; for Panel B, repeated light-condition visits were first averaged within participant and activity group. The dips in the active group correspond to the breaks between walking bouts.

**Results S2. Change in DLMO: Primary and Sensitivity Analyses**

Figure S6 shows the change in DLMO against the measured photopic illuminance, that is, against the predictor as it entered the models rather than by condition label. Table S5 reports the primary model alongside its five continuous-predictor sensitivity variants, and Table S6 translates their coefficients into high-versus-low contrasts. Across all variants, the light exposure and physical activity estimates pointed towards a phase advance, but every 95% HDI included zero and the Bayes factors remained inconclusive. Hence, the conclusions of the primary model did not depend on the likelihood, the prior width, the light metric, the laboratory light outliers, or demographic adjustment. The light × activity interaction was likewise inconclusive in all variants except under the wider coefficient priors, where its Bayes factor was 0.26 (below 1/3); this is an expected consequence of widening the prior, which penalises the alternative hypothesis more strongly. The categorical parameterisation (Table S7) yielded the same pattern.

**Figure S6**

*Change in DLMO as a Function of Measured Photopic Illuminance, by Activity Group*

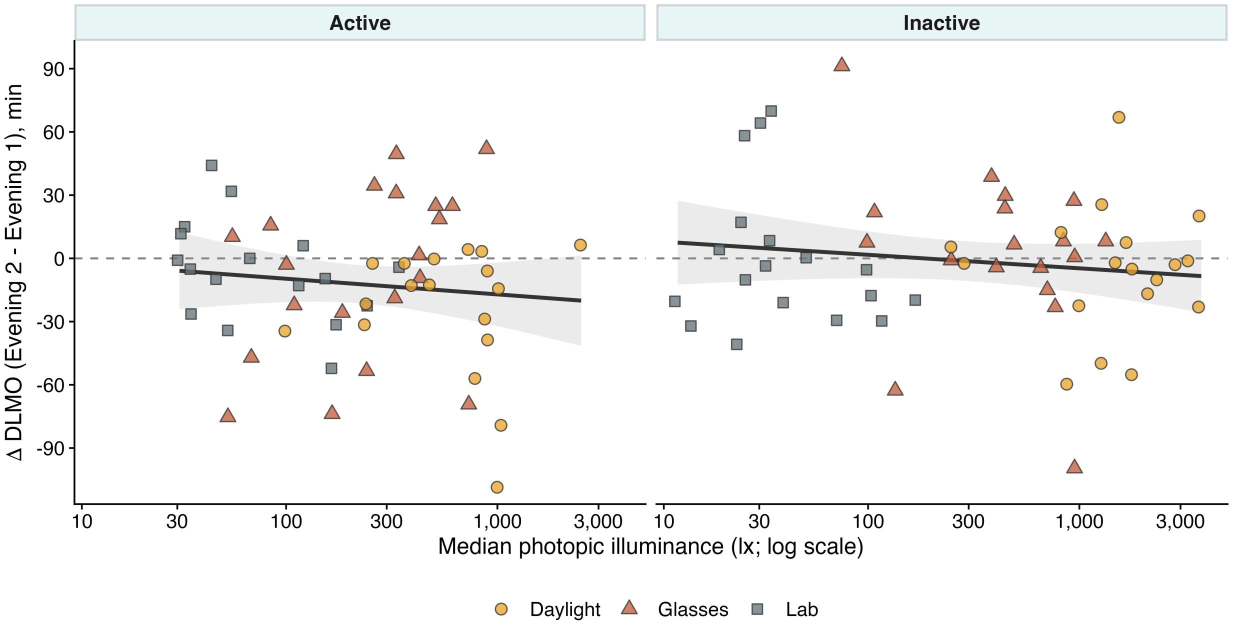

*Note.* Each point is one visit (Evening-2 minus Evening-1; negative values indicate a phase advance), plotted against the median photopic illuminance of the exposure window (log scale) and coded by light condition. Within each activity-group panel, the line is an unadjusted ordinary least-squares fit and the shaded band its 95% confidence interval; these are descriptive and are not predictions from the Bayesian model. The dashed line marks no change. Inferential results are based on the continuous models in Table S5.

**Table S5**

*Posterior Estimates for the Change in DLMO: Primary Model and Sensitivity Analyses*

| Effect | *b* | 95% HDI | *pd* | ROPE % | $BF_{10}$ |
|---|---|---|---|---|---|
| **Primary model** | | | | | |
| *Fixed effects* | | | | | |
| Intercept | −8.05 | [−15.90, −0.42] | — | — | — |
| Light exposure | −4.22 | [−10.93, 2.22] | .896 | 58.6 | 0.70 |
| Physical activity | −6.48 | [−13.79, 1.27] | .953 | 34.5 | 1.56 |
| Light × activity | 2.15 | [−4.50, 8.96] | .736 | 77.5 | 0.39 |
| *Random-effect and residual parameters* | | | | | |
| Participant intercept SD | 10.72 | [0.00, 19.78] | — | — | — |
| Residual SD | 29.81 | [24.34, 35.58] | — | — | — |
| **Student-*t* likelihood** | | | | | |
| *Fixed effects* | | | | | |
| Intercept | −7.66 | [−14.73, −0.04] | — | — | — |
| Light exposure | −3.32 | [−9.75, 3.45] | .838 | 68.7 | 0.52 |
| Physical activity | −5.78 | [−12.98, 1.42] | .942 | 41.4 | 1.20 |

*Table S5 (continued)*

| Effect | $b$ | 95% HDI | $pd$ | ROPE % | $BF_{10}$ |
|---|---|---|---|---|---|
| Light × activity | 1.56 | [−4.97, 8.27] | .678 | 82.1 | 0.34 |
| *Random-effect and residual parameters* | | | | | |
| Participant intercept SD | 9.21 | [0.00, 18.34] | — | — | — |
| Residual SD | 26.78 | [19.27, 33.71] | — | — | — |
| Degrees of freedom ($\nu$) | 16.58 | [1.62, 42.02] | — | — | — |
| **Wider coefficient priors** | | | | | |
| *Fixed effects* | | | | | |
| Intercept | −8.04 | [−15.67, −0.05] | — | — | — |
| Light exposure | −4.49 | [−11.31, 2.22] | .903 | 55.5 | 0.51 |
| Physical activity | −6.91 | [−14.43, 1.10] | .958 | 31.2 | 1.10 |
| Light × activity | 2.14 | [−4.81, 9.15] | .729 | 76.9 | 0.26 |
| *Random-effect and residual parameters* | | | | | |
| Participant intercept SD | 10.81 | [0.05, 19.84] | — | — | — |
| Residual SD | 29.80 | [24.45, 35.79] | — | — | — |
| **Cumulative light exposure** | | | | | |
| *Fixed effects* | | | | | |
| Intercept | −8.29 | [−16.04, −0.60] | — | — | — |
| Light exposure (cumulative) | −5.25 | [−11.52, 1.08] | .949 | 46.9 | 1.12 |
| Physical activity | −6.13 | [−13.70, 0.95] | .949 | 37.9 | 1.31 |
| Light × activity | 2.07 | [−4.31, 8.53] | .739 | 79.9 | 0.36 |
| *Random-effect and residual parameters* | | | | | |
| Participant intercept SD | 10.56 | [0.00, 19.55] | — | — | — |
| Residual SD | 29.67 | [24.22, 35.37] | — | — | — |
| **Corrected laboratory light values** | | | | | |
| *Fixed effects* | | | | | |
| Intercept | −8.02 | [−15.57, 0.05] | — | — | — |
| Light exposure | −4.08 | [−10.64, 2.51] | .889 | 60.4 | 0.64 |
| Physical activity | −6.53 | [−13.86, 0.96] | .954 | 34.0 | 1.51 |
| Light × activity | 2.12 | [−4.75, 8.60] | .733 | 77.9 | 0.38 |
| *Random-effect and residual parameters* | | | | | |
| Participant intercept SD | 10.79 | [0.01, 19.84] | — | — | — |
| Residual SD | 29.84 | [24.29, 35.61] | — | — | — |
| **Adjusted for age and sex** | | | | | |
| *Fixed effects* | | | | | |
| Intercept | −9.07 | [−17.14, −1.18] | — | — | — |
| Light exposure | −3.97 | [−10.54, 2.75] | .880 | 61.4 | 0.63 |
| Physical activity | −6.50 | [−13.85, 0.99] | .956 | 34.5 | 1.54 |
| Light × activity | 2.58 | [−4.03, 9.61] | .769 | 74.1 | 0.42 |
| Age | −0.32 | [−7.85, 6.62] | .534 | — | 0.34 |

*Table S5 (continued)*

| Effect | *b* | 95% HDI | *pd* | ROPE % | $BF_{10}$ |
|---|---|---|---|---|---|
| Sex (female − male) | 6.84 | [−6.21, 20.11] | .848 | — | 1.10 |
| *Random-effect and residual parameters* | | | | | |
| Participant intercept SD | 10.31 | [0.00, 19.60] | — | — | — |
| Residual SD | 29.93 | [24.39, 35.71] | — | — | — |

*Note.* Outcome: DLMO on Evening-2 minus DLMO on Evening-1 (min); negative values indicate a phase advance. Analysis sample: $N$ = 81 visits from 36 participants. Variants are specified in Method S7. Values are posterior means (*b*) with 95% highest-density intervals (HDI) on the outcome scale. Light exposure (median photopic illuminance, $\log_{10}$[lx + 1]) and physical activity (median heart rate reserve) were *z*-standardised (Method S6). Their main-effect coefficients give the expected outcome difference per 1-*SD* increase at the mean of the other predictor; the light × activity coefficient gives the change in the light exposure coefficient per 1-*SD* increase in physical activity. *pd* = probability of direction; ROPE % = percentage of the full posterior inside the outcome-specific region of practical equivalence (Table S3), shown for coefficients that are outcome differences (light exposure, physical activity, and light × activity) but not for intercepts, assessment terms (rates per assessment), adjustment terms, or variance parameters; $BF_{10}$ = Savage–Dickey Bayes factor against a point null at zero, computed with the prior of the respective coefficient; values above 1000 are shown as > 1000; SD = standard deviation; — = not applicable.

**Table S6**

*Derived Contrasts for the Change in DLMO: Primary Model and Sensitivity Analyses*

| Contrast | *b* | 95% HDI | *pd* | ROPE % |
|---|---|---|---|---|
| **Primary model** | | | | |
| Light exposure | −8.45 | [−21.86, 4.45] | .896 | 27.8 |
| Physical activity | −12.95 | [−27.57, 2.53] | .953 | 13.4 |
| Light exposure at high activity | −4.14 | [−26.10, 17.74] | .646 | 32.9 |
| Light exposure at low activity | −12.75 | [−28.41, 2.33] | .947 | 14.5 |
| Light × activity | 8.62 | [−18.00, 35.84] | .736 | — |
| **Student-*t* likelihood** | | | | |
| Light exposure | −6.63 | [−19.50, 6.91] | .838 | 36.3 |
| Physical activity | −11.57 | [−25.96, 2.83] | .942 | 17.2 |
| Light exposure at high activity | −3.51 | [−24.59, 17.83] | .633 | 34.4 |
| Light exposure at low activity | −9.76 | [−25.00, 6.47] | .887 | 24.4 |
| Light × activity | 6.25 | [−19.88, 33.10] | .678 | — |
| **Wider coefficient priors** | | | | |
| Light exposure | −8.99 | [−22.62, 4.43] | .903 | 26.0 |
| Physical activity | −13.83 | [−28.86, 2.19] | .958 | 12.0 |
| Light exposure at high activity | −4.71 | [−27.98, 17.66] | .657 | 31.5 |
| Light exposure at low activity | −13.27 | [−29.26, 2.11] | .954 | 13.6 |
| Light × activity | 8.57 | [−19.24, 36.60] | .729 | — |

*Table S6 (continued)*

| Contrast | *b* | 95% HDI | *pd* | ROPE % |
|---|---|---|---|---|
| **Cumulative light exposure** | | | | |
| Light exposure | −10.51 | [−23.04, 2.15] | .949 | 18.5 |
| Physical activity | −12.25 | [−27.40, 1.89] | .949 | 15.5 |
| Light exposure at high activity | −6.36 | [−26.38, 13.78] | .736 | 31.4 |
| Light exposure at low activity | −14.65 | [−30.09, 1.42] | .965 | 10.7 |
| Light × activity | 8.29 | [−17.25, 34.14] | .739 | — |
| **Corrected laboratory light values** | | | | |
| Light exposure | −8.16 | [−21.28, 5.03] | .889 | 28.9 |
| Physical activity | −13.05 | [−27.72, 1.92] | .954 | 13.4 |
| Light exposure at high activity | −3.92 | [−25.29, 17.71] | .642 | 33.3 |
| Light exposure at low activity | −12.41 | [−27.87, 2.97] | .943 | 15.8 |
| Light × activity | 8.49 | [−19.00, 34.40] | .733 | — |
| **Adjusted for age and sex** | | | | |
| Light exposure | −7.94 | [−21.09, 5.51] | .880 | 29.9 |
| Physical activity | −12.99 | [−27.70, 1.98] | .956 | 13.6 |
| Light exposure at high activity | −2.79 | [−25.23, 19.59] | .599 | 33.1 |
| Light exposure at low activity | −13.09 | [−28.66, 1.76] | .954 | 13.6 |
| Light × activity | 10.31 | [−16.11, 38.42] | .769 | — |

*Note.* Outcome: change in DLMO (min); negative values indicate a phase advance. Values are posterior means (*b*) with 95% highest-density intervals (HDI) on the outcome scale. Rows labelled light exposure or physical activity compare high (+1 *SD*) with low (−1 *SD*) values of that predictor, that is, a 2-*SD* difference, with the other predictor at its mean. Rows labelled at high or low activity evaluate the light exposure contrast at +1 *SD* or −1 *SD* physical activity, and the light × activity row is the difference between these two rows (difference-in-differences). *pd* = probability of direction; ROPE % = percentage of the full posterior inside the outcome-specific region of practical equivalence (Table S3). Because the contrasts span 2 *SD*s, their ROPE percentages are not directly comparable with those of the per-*SD* coefficients. ROPE percentages are not reported for difference-in-differences or time-course moderation contrasts because no separate practical-equivalence threshold was prespecified for those estimands. Bayes factors are not reported for derived contrasts (Method S6): for contrasts that rescale a single coefficient, they equal the Bayes factor of that coefficient in the corresponding estimates table; for contrasts that combine several coefficients, they would depend on a prior that is implied by the coefficient priors rather than specified for the contrast.

**Table S7**

*Change in DLMO by Light Condition and Activity Group: Categorical Model*

| Contrast | *b* (min) | 95% HDI | *pd* |
|---|---|---|---|
| **Light condition contrasts** | | | |
| Daylight vs. laboratory, inactive | −5.38 | [−18.74, 7.45] | .792 |
| Glasses vs. laboratory, inactive | 2.34 | [−10.92, 15.62] | .640 |
| Daylight vs. glasses, inactive | −7.72 | [−23.36, 7.94] | .831 |
| **Activity group contrasts** | | | |
| Active vs. inactive, laboratory | −6.07 | [−19.45, 7.00] | .817 |
| Active vs. inactive, daylight | −14.08 | [−30.59, 3.26] | .947 |
| Active vs. inactive, glasses | −7.10 | [−23.81, 9.62] | .799 |
| **Interaction contrasts** | | | |
| Activity × (daylight vs. laboratory) | −8.02 | [−24.28, 7.23] | .840 |
| Activity × (glasses vs. laboratory) | −1.04 | [−16.86, 14.50] | .547 |

*Note.* Posterior means (*b*, min) and 95% highest-density intervals from the categorical sensitivity model (Method S7), with laboratory light and the inactive group as reference levels; negative values indicate a larger phase advance in the first-named condition or group. Interaction contrasts give the difference in the light-condition contrast between the active and the inactive group. *pd* = probability of direction. The model was used descriptively; ROPE proportions and Bayes factors are not reported for it. Analysis sample: $N$ = 110 visits from 41 participants. Nine visits with missing change in DLMO were excluded. Assigned light condition and activity group were available for all visits; measured illuminance and HRR were not required by this model.

#### Results S3. Exploratory Analysis of Evening Melatonin Secretion

The exploratory analysis of evening melatonin secretion converged with the DLMO results (Table S8): the light exposure association was inconclusive, the data provided moderate evidence against a practically relevant physical activity association, and the light × activity interaction was inconclusive.

**Table S8**

*Posterior Estimates and Derived Contrasts for the Change in Evening Melatonin AUC (Exploratory)*

| Effect | *b* | 95% HDI | *pd* | ROPE % | $BF_{10}$ |
|---|---|---|---|---|---|
| **Model parameters** | | | | | |
| *Fixed effects* | | | | | |
| Intercept | 0.017 | [−0.013, 0.048] | — | — | — |
| Light exposure | 0.029 | [0.001, 0.057] | .977 | 80.6 | 2.30 |
| Physical activity | 0.001 | [−0.028, 0.030] | .531 | 99.3 | 0.32 |
| Light × activity | −0.010 | [−0.039, 0.020] | .748 | 98.1 | 0.40 |
| *Random-effect and residual parameters* | | | | | |
| Participant intercept SD | 0.027 | [0.000, 0.061] | — | — | — |
| Residual SD | 0.134 | [0.112, 0.155] | — | — | — |
| **Derived contrasts** | | | | | |
| Light exposure | 0.057 | [0.002, 0.114] | .977 | 27.9 | — |
| Physical activity | 0.002 | [−0.057, 0.060] | .531 | 83.0 | — |
| Light exposure at high activity | 0.038 | [−0.056, 0.131] | .789 | 47.4 | — |
| Light exposure at low activity | 0.077 | [0.013, 0.142] | .989 | 13.3 | — |
| Light × activity | −0.039 | [−0.154, 0.078] | .748 | — | — |

*Note.* Outcome: $\log_{10}$(AUC on Evening-2 / AUC on Evening-1); positive values indicate greater melatonin secretion on the post-exposure evening, and a difference of 0.041 $\log_{10}$ units corresponds to a ratio of approximately 1.10. The model had the same structure, priors, and sampler settings as the primary DLMO model. Analysis sample: $N = 87$ visits from 37 participants. Values are given to three decimals because of the small outcome scale. Values are posterior means (*b*) with 95% highest-density intervals (HDI) on the outcome scale. Light exposure (median photopic illuminance, $\log_{10}$[lx + 1]) and physical activity (median heart rate reserve) were *z*-standardised (Method S6). Their main-effect coefficients give the expected outcome difference per 1-*SD* increase at the mean of the other predictor; the light × activity coefficient gives the change in the light exposure coefficient per 1-*SD* increase in physical activity. *pd* = probability of direction; ROPE % = percentage of the full posterior inside the outcome-specific region of practical equivalence (Table S3), shown for coefficients that are outcome differences (light exposure, physical activity, and light × activity) but not for intercepts, assessment terms (rates per assessment), adjustment terms, or variance parameters; $BF_{10}$ = Savage–Dickey Bayes factor against a point null at zero, computed with the prior of the respective coefficient; values above 1000 are shown as > 1000; SD = standard deviation; — = not applicable. Derived contrasts are defined as in Table S6; Bayes factors are not reported for them (Method S6).

#### Results S4. Evening Time Course

Figure S7 shows the light exposure and physical activity coefficients relative to their ROPEs; full model estimates and derived contrasts are reported in Tables S9–S10.

**Figure S7**

*Posterior Distributions of Light Exposure and Physical Activity Effects on Evening Outcomes*

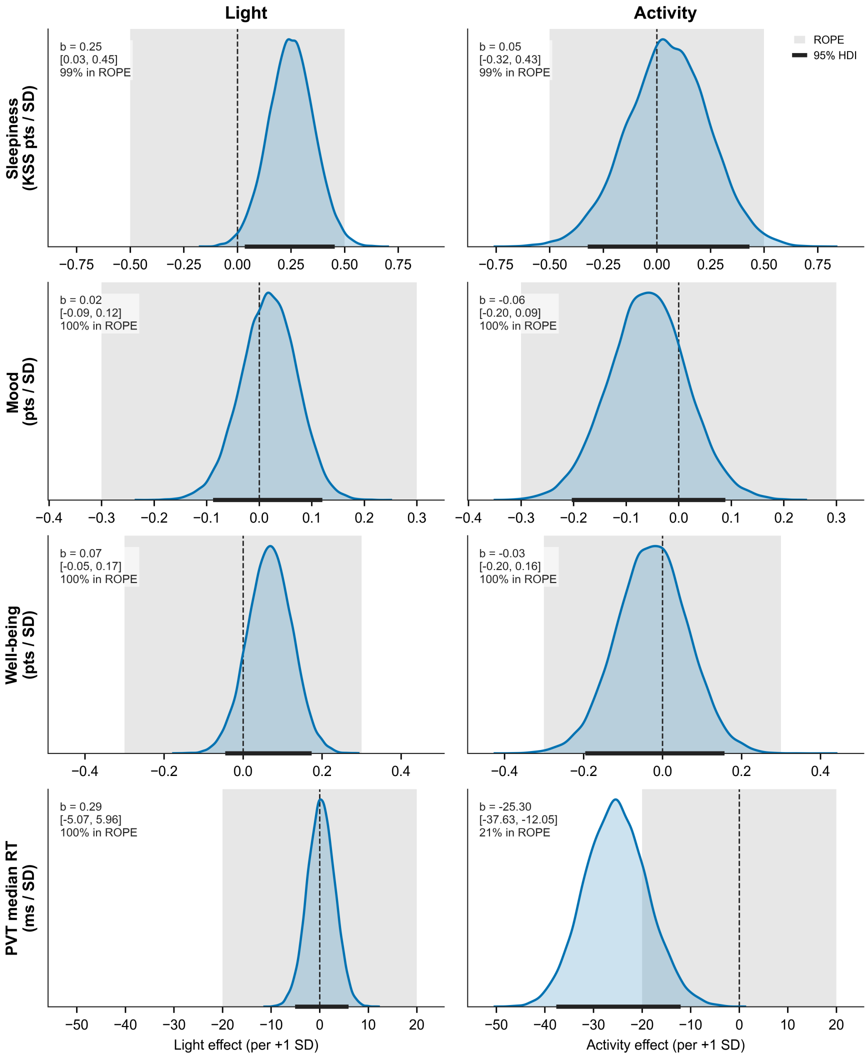

*Note.* Posterior densities of the light exposure (left) and physical activity (right) coefficients per 1-*SD* increase for sleepiness (KSS), mood, wellbeing, and PVT median reaction time on Evening-2. Curves show posterior densities; shaded grey bands indicate the outcome-specific region of practical equivalence (ROPE), black horizontal bars the 95% highest-density intervals (HDIs), and dashed vertical lines zero. Annotations report *b* [95% HDI] and the percentage of the posterior within the ROPE. Effects are expressed per 1-*SD* increase, with the other predictor at its mean. Estimates correspond to Table S9.

For all outcomes, the 95% HDIs of the assessment × light and assessment × activity terms included zero, and their Bayes factors were below 1/3. Hence, the data favoured the absence of a dependence of the evening trajectory on either predictor.

**Table S9**

*Posterior Estimates for Evening Time-Course Outcomes*

| Effect | $b$ | 95% HDI | $pd$ | ROPE % | $BF_{10}$ |
|---|---|---|---|---|---|
| **Sleepiness (KSS)** | | | | | |
| *Fixed effects* | | | | | |
| Intercept | 5.61 | [5.18, 6.03] | — | — | — |
| Assessment (time course) | 0.30 | [0.25, 0.35] | > .999 | — | > 1000 |
| Light exposure | 0.25 | [0.03, 0.45] | .991 | 99.0 | 2.76 |
| Physical activity | 0.05 | [−0.32, 0.43] | .611 | 98.8 | 0.33 |
| Light × activity | 0.15 | [−0.07, 0.35] | .915 | 99.9 | 0.46 |
| Assessment × light | 0.02 | [−0.01, 0.06] | .903 | — | 0.07 |
| Assessment × activity | 0.02 | [−0.03, 0.07] | .790 | — | 0.06 |
| *Random-effect and residual parameters* | | | | | |
| Participant intercept SD | 1.14 | [0.81, 1.49] | — | — | — |
| Participant assessment-slope SD | 0.10 | [0.02, 0.17] | — | — | — |
| Visit intercept SD | 0.80 | [0.63, 0.99] | — | — | — |
| Visit assessment-slope SD | 0.14 | [0.10, 0.18] | — | — | — |
| Residual SD | 0.67 | [0.63, 0.70] | — | — | — |
| **Mood** | | | | | |
| *Fixed effects* | | | | | |
| Intercept | 3.78 | [3.62, 3.94] | — | — | — |
| Assessment (time course) | −0.02 | [−0.04, −0.01] | .993 | — | 0.94 |
| Light exposure | 0.02 | [−0.09, 0.12] | .641 | 100.0 | 0.26 |
| Physical activity | −0.06 | [−0.20, 0.09] | .788 | 100.0 | 0.44 |
| Light × activity | 0.00 | [−0.10, 0.11] | .535 | 100.0 | 0.24 |
| Assessment × light | −0.01 | [−0.03, 0.00] | .944 | — | 0.12 |
| Assessment × activity | 0.00 | [−0.02, 0.01] | .609 | — | 0.04 |
| *Random-effect and residual parameters* | | | | | |
| Participant intercept SD | 0.37 | [0.22, 0.55] | — | — | — |
| Participant assessment-slope SD | 0.01 | [0.00, 0.03] | — | — | — |
| Visit intercept SD | 0.41 | [0.32, 0.51] | — | — | — |
| Visit assessment-slope SD | 0.06 | [0.04, 0.07] | — | — | — |
| Residual SD | 0.38 | [0.36, 0.40] | — | — | — |
| **Wellbeing** | | | | | |
| *Fixed effects* | | | | | |
| Intercept | 3.85 | [3.66, 4.04] | — | — | — |

*Table S9 (continued)*

| Effect | *b* | 95% HDI | *pd* | ROPE % | $BF_{10}$ |
|---|---|---|---|---|---|
| Assessment (time course) | −0.02 | [−0.03, 0.00] | .955 | — | 0.16 |
| Light exposure | 0.07 | [−0.05, 0.17] | .889 | 100.0 | 0.47 |
| Physical activity | −0.03 | [−0.20, 0.16] | .609 | 99.8 | 0.37 |
| Light × activity | 0.03 | [−0.09, 0.14] | .683 | 100.0 | 0.26 |
| Assessment × light | −0.01 | [−0.03, 0.00] | .962 | — | 0.14 |
| Assessment × activity | 0.01 | [−0.01, 0.03] | .770 | — | 0.05 |
| *Random-effect and residual parameters* | | | | | |
| Participant intercept SD | 0.50 | [0.33, 0.68] | — | — | — |
| Participant assessment-slope SD | 0.04 | [0.02, 0.06] | — | — | — |
| Visit intercept SD | 0.44 | [0.34, 0.54] | — | — | — |
| Visit assessment-slope SD | 0.04 | [0.03, 0.06] | — | — | — |
| Residual SD | 0.34 | [0.32, 0.36] | — | — | — |
| **PVT median reaction time (ms)** | | | | | |
| *Fixed effects* | | | | | |
| Intercept | 318.98 | [304.97, 333.64] | — | — | — |
| Assessment (time course) | 3.34 | [2.58, 4.06] | > .999 | — | > 1000 |
| Light exposure | 0.29 | [−5.07, 5.96] | .542 | 100.0 | 0.14 |
| Physical activity | −25.30 | [−37.63, −12.05] | > .999 | 20.6 | 253.94 |
| Light × activity | −0.53 | [−6.10, 5.06] | .571 | 100.0 | 0.15 |
| Assessment × light | 0.12 | [−0.52, 0.77] | .647 | — | 0.02 |
| Assessment × activity | −0.01 | [−0.79, 0.76] | .517 | — | 0.02 |
| *Random-effect and residual parameters* | | | | | |
| Participant intercept SD | 41.48 | [30.16, 53.19] | — | — | — |
| Participant assessment-slope SD | 1.10 | [0.00, 2.03] | — | — | — |
| Visit intercept SD | 20.67 | [16.17, 25.83] | — | — | — |
| Visit assessment-slope SD | 2.26 | [1.54, 3.00] | — | — | — |
| Residual SD | 16.45 | [15.58, 17.36] | — | — | — |
| **PVT mean response speed ($s^{-1}$)** | | | | | |
| *Fixed effects* | | | | | |
| Intercept | 3.228 | [3.090, 3.371] | — | — | — |
| Assessment (time course) | −0.034 | [−0.041, −0.027] | > .999 | — | > 1000 |
| Light exposure | −0.008 | [−0.063, 0.044] | .611 | 96.5 | 0.14 |
| Physical activity | 0.271 | [0.141, 0.396] | > .999 | 0.1 | 716.45 |
| Light × activity | 0.005 | [−0.048, 0.060] | .566 | 96.7 | 0.14 |
| Assessment × light | 0.000 | [−0.006, 0.005] | .555 | — | 0.01 |
| Assessment × activity | −0.007 | [−0.014, 0.001] | .968 | — | 0.10 |
| *Random-effect and residual parameters* | | | | | |
| Participant intercept SD | 0.406 | [0.297, 0.520] | — | — | — |
| Participant assessment-slope SD | 0.012 | [0.000, 0.021] | — | — | — |
| Visit intercept SD | 0.200 | [0.155, 0.247] | — | — | — |

*Table S9 (continued)*

| Effect | $b$ | 95% HDI | $pd$ | ROPE % | $BF_{10}$ |
|---|---|---|---|---|---|
| Visit assessment-slope SD | 0.020 | [0.013, 0.027] | — | — | — |
| Residual SD | 0.144 | [0.137, 0.152] | — | — | — |
| **PVT slowest-10% response speed ($s^{-1}$)** | | | | | |
| *Fixed effects* | | | | | |
| Intercept | 2.312 | [2.191, 2.439] | — | — | — |
| Assessment (time course) | −0.041 | [−0.049, −0.033] | > .999 | — | > 1000 |
| Light exposure | −0.003 | [−0.050, 0.043] | .550 | 96.4 | 0.14 |
| Physical activity | 0.162 | [0.056, 0.265] | .998 | 2.0 | 21.25 |
| Light × activity | 0.003 | [−0.042, 0.050] | .555 | 96.4 | 0.15 |
| Assessment × light | −0.003 | [−0.009, 0.003] | .798 | — | 0.03 |
| Assessment × activity | −0.005 | [−0.013, 0.003] | .891 | — | 0.05 |
| *Random-effect and residual parameters* | | | | | |
| Participant intercept SD | 0.353 | [0.262, 0.445] | — | — | — |
| Participant assessment-slope SD | 0.014 | [0.001, 0.024] | — | — | — |
| Visit intercept SD | 0.166 | [0.127, 0.207] | — | — | — |
| Visit assessment-slope SD | 0.020 | [0.011, 0.029] | — | — | — |
| Residual SD | 0.183 | [0.173, 0.193] | — | — | — |

*Note.* Analysis sample: $N = 827$ assessments from 37 participants across 84 visits. Assessment was centred at the midpoint of the ten Evening-2 assessments (30-min intervals), so main effects refer to the evening midpoint and the assessment term gives the change per assessment. Random-effect parameters are the *SD*s of participant- and visit-level intercepts and assessment slopes. Values for response speed ($s^{-1}$) are given to three decimals because of the small outcome scale. Values are posterior means ($b$) with 95% highest-density intervals (HDI) on the outcome scale. Light exposure (median photopic illuminance, $\log_{10}[\text{lx} + 1]$) and physical activity (median heart rate reserve) were $z$-standardised (Method S6). Their main-effect coefficients give the expected outcome difference per 1-*SD* increase at the mean of the other predictor; the light × activity coefficient gives the change in the light exposure coefficient per 1-*SD* increase in physical activity; assessment terms give the change per assessment. $pd$ = probability of direction; ROPE % = percentage of the full posterior inside the outcome-specific region of practical equivalence (Table S3), shown for coefficients that are outcome differences (light exposure, physical activity, and light × activity) but not for intercepts, assessment terms (rates per assessment), adjustment terms, or variance parameters; $BF_{10}$ = Savage–Dickey Bayes factor against a point null at zero, computed with the prior of the respective coefficient; values above 1000 are shown as > 1000; SD = standard deviation; — = not applicable. KSS = Karolinska Sleepiness Scale; PVT = psychomotor vigilance task.

**Table S10**

*Derived Contrasts for Evening Time-Course Outcomes*

| Contrast | *b* | 95% HDI | *pd* | ROPE % |
|---|---|---|---|---|
| **Sleepiness (KSS)** | | | | |
| Full-evening change | 2.70 | [2.24, 3.14] | > .999 | 0.0 |
| Light exposure: first assessment | 0.29 | [−0.22, 0.83] | .865 | 77.9 |
| Light exposure: midpoint | 0.50 | [0.07, 0.91] | .991 | 49.5 |
| Light exposure: final assessment | 0.71 | [0.19, 1.24] | .995 | 21.0 |
| Physical activity: first assessment | −0.08 | [−0.98, 0.76] | .566 | 73.6 |
| Physical activity: midpoint | 0.10 | [−0.64, 0.86] | .611 | 79.3 |
| Physical activity: final assessment | 0.28 | [−0.61, 1.14] | .737 | 64.4 |
| Light exposure at high activity: midpoint | 0.80 | [0.08, 1.50] | .986 | 20.0 |
| Light exposure at low activity: midpoint | 0.21 | [−0.24, 0.66] | .820 | 89.9 |
| Light × activity: midpoint | 0.59 | [−0.29, 1.42] | .915 | — |
| Assessment × light: full evening | 0.42 | [−0.22, 1.05] | .903 | — |
| Assessment × activity: full evening | 0.36 | [−0.53, 1.26] | .790 | — |
| **Mood** | | | | |
| Full-evening change | −0.20 | [−0.35, −0.05] | .993 | 91.6 |
| Light exposure: first assessment | 0.15 | [−0.10, 0.40] | .878 | 88.6 |
| Light exposure: midpoint | 0.04 | [−0.18, 0.24] | .641 | 99.3 |
| Light exposure: final assessment | −0.07 | [−0.33, 0.17] | .718 | 95.7 |
| Physical activity: first assessment | −0.09 | [−0.42, 0.24] | .715 | 88.2 |
| Physical activity: midpoint | −0.12 | [−0.41, 0.18] | .788 | 88.9 |
| Physical activity: final assessment | −0.14 | [−0.46, 0.19] | .799 | 83.1 |
| Light exposure at high activity: midpoint | 0.05 | [−0.30, 0.40] | .606 | 90.0 |
| Light exposure at low activity: midpoint | 0.03 | [−0.20, 0.26] | .599 | 98.6 |
| Light × activity: midpoint | 0.02 | [−0.40, 0.43] | .535 | — |
| Assessment × light: full evening | −0.22 | [−0.50, 0.06] | .944 | — |
| Assessment × activity: full evening | −0.04 | [−0.34, 0.27] | .609 | — |
| **Wellbeing** | | | | |
| Full-evening change | −0.14 | [−0.31, 0.02] | .955 | 97.1 |
| Light exposure: first assessment | 0.25 | [0.00, 0.50] | .973 | 65.8 |
| Light exposure: midpoint | 0.14 | [−0.09, 0.35] | .889 | 92.9 |
| Light exposure: final assessment | 0.03 | [−0.23, 0.28] | .577 | 97.8 |
| Physical activity: first assessment | −0.11 | [−0.49, 0.28] | .714 | 81.3 |
| Physical activity: midpoint | −0.05 | [−0.39, 0.31] | .609 | 89.2 |
| Physical activity: final assessment | 0.01 | [−0.38, 0.40] | .522 | 87.2 |
| Light exposure at high activity: midpoint | 0.19 | [−0.19, 0.55] | .842 | 71.6 |
| Light exposure at low activity: midpoint | 0.08 | [−0.16, 0.33] | .750 | 95.9 |
| Light × activity: midpoint | 0.11 | [−0.35, 0.55] | .683 | — |

*Table S10 (continued)*

| Contrast | *b* | 95% HDI | *pd* | ROPE % |
|---|---|---|---|---|
| Assessment × light: full evening | −0.22 | [−0.46, 0.02] | .962 | — |
| Assessment × activity: full evening | 0.12 | [−0.21, 0.46] | .770 | — |
| **PVT median reaction time (ms)** | | | | |
| Full-evening change | 30.08 | [23.19, 36.56] | > .999 | 0.2 |
| Light exposure: first assessment | −0.51 | [−13.08, 11.86] | .532 | 99.8 |
| Light exposure: midpoint | 0.59 | [−10.14, 11.93] | .542 | 99.9 |
| Light exposure: final assessment | 1.68 | [−10.77, 13.89] | .603 | 99.8 |
| Physical activity: first assessment | −50.47 | [−77.10, −24.12] | > .999 | 1.4 |
| Physical activity: midpoint | −50.59 | [−75.25, −24.10] | > .999 | 1.2 |
| Physical activity: final assessment | −50.72 | [−77.17, −24.22] | > .999 | 1.4 |
| Light exposure at high activity: midpoint | −0.47 | [−18.98, 18.75] | .518 | 96.2 |
| Light exposure at low activity: midpoint | 1.64 | [−9.98, 13.87] | .609 | 99.8 |
| Light × activity: midpoint | −2.11 | [−24.40, 20.23] | .571 | — |
| Assessment × light: full evening | 2.19 | [−9.35, 13.79] | .647 | — |
| Assessment × activity: full evening | −0.25 | [−14.24, 13.68] | .517 | — |
| **PVT mean response speed ($s^{-1}$)** | | | | |
| Full-evening change | −0.309 | [−0.373, −0.244] | > .999 | 0.0 |
| Light exposure: first assessment | −0.012 | [−0.130, 0.107] | .576 | 68.2 |
| Light exposure: midpoint | −0.015 | [−0.126, 0.088] | .611 | 71.9 |
| Light exposure: final assessment | −0.019 | [−0.137, 0.100] | .619 | 66.2 |
| Physical activity: first assessment | 0.602 | [0.342, 0.866] | > .999 | 0.0 |
| Physical activity: midpoint | 0.542 | [0.282, 0.793] | > .999 | 0.0 |
| Physical activity: final assessment | 0.481 | [0.221, 0.745] | > .999 | 0.1 |
| Light exposure at high activity: midpoint | −0.006 | [−0.190, 0.176] | .524 | 49.0 |
| Light exposure at low activity: midpoint | −0.024 | [−0.141, 0.091] | .665 | 65.9 |
| Light × activity: midpoint | 0.018 | [−0.193, 0.242] | .566 | — |
| Assessment × light: full evening | −0.007 | [−0.110, 0.096] | .555 | — |
| Assessment × activity: full evening | −0.120 | [−0.250, 0.010] | .968 | — |
| **PVT slowest-10% response speed ($s^{-1}$)** | | | | |
| Full-evening change | −0.366 | [−0.439, −0.295] | > .999 | 0.0 |
| Light exposure: first assessment | 0.019 | [−0.091, 0.126] | .634 | 61.5 |
| Light exposure: midpoint | −0.006 | [−0.100, 0.086] | .550 | 71.5 |
| Light exposure: final assessment | −0.030 | [−0.140, 0.076] | .706 | 56.8 |
| Physical activity: first assessment | 0.369 | [0.144, 0.588] | .999 | 0.3 |
| Physical activity: midpoint | 0.323 | [0.112, 0.531] | .998 | 0.6 |
| Physical activity: final assessment | 0.278 | [0.058, 0.500] | .991 | 2.1 |
| Light exposure at high activity: midpoint | 0.001 | [−0.153, 0.160] | .503 | 47.7 |
| Light exposure at low activity: midpoint | −0.012 | [−0.111, 0.084] | .594 | 67.4 |

*Table S10 (continued)*

| Contrast | *b* | 95% HDI | *pd* | ROPE % |
|---|---|---|---|---|
| Light × activity: midpoint | 0.013 | [−0.170, 0.200] | .555 | — |
| Assessment × light: full evening | −0.049 | [−0.167, 0.061] | .798 | — |
| Assessment × activity: full evening | −0.091 | [−0.238, 0.056] | .891 | — |

*Note.* Full-evening change = predicted change from the first to the final of the ten Evening-2 assessments at mean light exposure and physical activity. First assessment, midpoint, and final assessment indicate where a contrast is evaluated; the midpoint is the centre of the assessment series, at which the main effects in Table S9 are estimated. Assessment × light and assessment × activity = difference in full-evening change between high and low values of the predictor. Values for response speed ($s^{-1}$) are given to three decimals. Values are posterior means (*b*) with 95% highest-density intervals (HDI) on the outcome scale. Rows labelled light exposure or physical activity compare high (+1 *SD*) with low (−1 *SD*) values of that predictor, that is, a 2-*SD* difference, with the other predictor at its mean. Rows labelled at high or low activity evaluate the light exposure contrast at +1 *SD* or −1 *SD* physical activity, and the light × activity row is the difference between these two rows (difference-in-differences). *pd* = probability of direction; ROPE % = percentage of the full posterior inside the outcome-specific region of practical equivalence (Table S3). Because the contrasts span 2 *SD*s, their ROPE percentages are not directly comparable with those of the per-*SD* coefficients. ROPE percentages are not reported for difference-in-differences or time-course moderation contrasts because no separate practical-equivalence threshold was prespecified for those estimands. Bayes factors are not reported for derived contrasts (Method S6): for contrasts that rescale a single coefficient, they equal the Bayes factor of that coefficient in the corresponding estimates table; for contrasts that combine several coefficients, they would depend on a prior that is implied by the coefficient priors rather than specified for the contrast. KSS = Karolinska Sleepiness Scale; PVT = psychomotor vigilance task.

#### Results S5. Baseline-Adjusted Psychomotor Vigilance

Table S11 reports all parameters of the baseline-adjusted model presented in the main text (M1) for the two PVT outcomes reported in the main text and the preregistered slowest-decile response speed. Table S12, in contrast, shows only the light exposure and physical activity coefficients, across all four models (M0–M3; Method S8), and thus how these coefficients depend on the adjustment. Without baseline adjustment (M0), higher physical activity was associated with faster responses on all three outcomes, with Bayes factors above 3. Once Evening-1 performance was taken into account, this association was no longer evidenced: in M1 and M2, all 95% HDIs included zero and all Bayes factors were below 1/3. In M3, the within-participant activity component, which reflected visit-to-visit variation in achieved HRR rather than the between-participant activity allocation, was estimated with considerably wider HDIs, and its Bayes factors were inconclusive. The attenuation after baseline adjustment was consistent with a contribution of pre-existing between-participant differences in vigilance to the unadjusted association. These observational decompositions did not isolate a causal effect of activity allocation. The light exposure coefficients, in contrast, remained small in all models, with Bayes factors below 1/3. Note that M0 was not identical to the primary evening model (Results S4), because it was fitted only to the assessments for which an Evening-1 baseline was available.

**Table S11**

*Posterior Estimates for Baseline-Adjusted Psychomotor Vigilance Outcomes (Model M1)*

| Effect | *b* | 95% HDI | *pd* | ROPE % | $BF_{10}$ |
|---|---|---|---|---|---|
| **PVT median reaction time (ms)** | | | | | |
| *Fixed effects* | | | | | |
| Intercept | 319.77 | [314.94, 324.65] | — | — | — |
| Assessment (time course) | 3.20 | [2.43, 3.93] | > .999 | — | > 1000 |
| Light exposure | 1.70 | [−3.01, 6.32] | .766 | 100.0 | 0.16 |
| Physical activity | −2.67 | [−9.41, 3.92] | .784 | 100.0 | 0.23 |
| Light × activity | 2.67 | [−2.10, 7.39] | .870 | 100.0 | 0.23 |
| Assessment × light | −0.02 | [−0.64, 0.64] | .523 | — | 0.02 |
| Assessment × activity | 0.06 | [−0.71, 0.84] | .551 | — | 0.02 |
| Baseline (visit level) | 49.74 | [42.95, 56.51] | > .999 | — | > 1000 |
| *Random-effect and residual parameters* | | | | | |
| Participant intercept SD | 6.11 | [0.00, 12.79] | — | — | — |
| Participant assessment-slope SD | 1.03 | [0.00, 1.96] | — | — | — |
| Visit intercept SD | 18.82 | [15.03, 22.62] | — | — | — |
| Visit assessment-slope SD | 2.22 | [1.50, 2.96] | — | — | — |
| Residual SD | 16.51 | [15.59, 17.40] | — | — | — |
| **PVT mean response speed ($s^{-1}$)** | | | | | |
| *Fixed effects* | | | | | |
| Intercept | 3.217 | [3.171, 3.262] | — | — | — |
| Assessment (time course) | −0.033 | [−0.040, −0.026] | > .999 | — | > 1000 |
| Light exposure | −0.025 | [−0.069, 0.019] | .866 | 94.1 | 0.20 |
| Physical activity | 0.032 | [−0.034, 0.095] | .838 | 80.3 | 0.27 |
| Light × activity | −0.030 | [−0.075, 0.014] | .904 | 90.8 | 0.27 |
| Assessment × light | 0.001 | [−0.004, 0.007] | .671 | — | 0.02 |
| Assessment × activity | −0.007 | [−0.014, 0.000] | .975 | — | 0.12 |

*Table S11 (continued)*

| Effect | *b* | 95% HDI | *pd* | ROPE % | $BF_{10}$ |
|---|---|---|---|---|---|
| Baseline (visit level) | 0.503 | [0.438, 0.565] | > .999 | — | > 1000 |
| *Random-effect and residual parameters* | | | | | |
| Participant intercept SD | 0.050 | [0.000, 0.107] | — | — | — |
| Participant assessment-slope SD | 0.011 | [0.000, 0.019] | — | — | — |
| Visit intercept SD | 0.178 | [0.146, 0.213] | — | — | — |
| Visit assessment-slope SD | 0.020 | [0.013, 0.026] | — | — | — |
| Residual SD | 0.144 | [0.136, 0.152] | — | — | — |
| **PVT slowest-10% response speed ($s^{-1}$)** | | | | | |
| *Fixed effects* | | | | | |
| Intercept | 2.298 | [2.256, 2.340] | — | — | — |
| Assessment (time course) | −0.039 | [−0.047, −0.031] | > .999 | — | > 1000 |
| Light exposure | −0.026 | [−0.066, 0.014] | .895 | 88.7 | 0.27 |
| Physical activity | 0.008 | [−0.045, 0.059] | .624 | 92.9 | 0.17 |
| Light × activity | −0.035 | [−0.076, 0.006] | .954 | 76.1 | 0.54 |
| Assessment × light | −0.002 | [−0.008, 0.005] | .691 | — | 0.02 |
| Assessment × activity | −0.004 | [−0.013, 0.004] | .860 | — | 0.04 |
| Baseline (visit level) | 0.407 | [0.353, 0.460] | > .999 | — | > 1000 |
| *Random-effect and residual parameters* | | | | | |
| Participant intercept SD | 0.051 | [0.000, 0.107] | — | — | — |
| Participant assessment-slope SD | 0.013 | [0.000, 0.023] | — | — | — |
| Visit intercept SD | 0.156 | [0.125, 0.190] | — | — | — |
| Visit assessment-slope SD | 0.020 | [0.011, 0.029] | — | — | — |
| Residual SD | 0.183 | [0.173, 0.193] | — | — | — |

*Note.* Analysis sample: $N$ = 807 assessments from 37 participants across 82 visits. M1 contains all terms of the evening model plus the Evening-1 baseline (median of the respective outcome on Evening-1) as a single visit-level covariate, centred at the grand mean and divided by its *SD* (Method S8); the baseline coefficient therefore gives the outcome difference per 1 *SD* of the baseline. Values for response speed ($s^{-1}$) are given to three decimals because of the small outcome scale. Values are posterior means (*b*) with 95% highest-density intervals (HDI) on the outcome scale. Light exposure (median photopic illuminance, $\log_{10}[\mathrm{lx} + 1]$) and physical activity (median heart rate reserve) were *z*-standardised (Method S6). Their main-effect coefficients give the expected outcome difference per 1-*SD* increase at the mean of the other predictor; the light × activity coefficient gives the change in the light exposure coefficient per 1-*SD* increase in physical activity; assessment terms give the change per assessment. *pd* = probability of direction; ROPE % = percentage of the full posterior inside the outcome-specific region of practical equivalence (Table S3), shown for coefficients that are outcome differences (light exposure, physical activity, and light × activity) but not for intercepts, assessment terms (rates per assessment), adjustment terms, or variance parameters; $BF_{10}$ = Savage–Dickey Bayes factor against a point null at zero, computed with the prior of the respective coefficient; values above 1000 are shown as > 1000; SD = standard deviation; — = not applicable. PVT = psychomotor vigilance task.

**Table S12**

*Light Exposure and Physical Activity Coefficients Across Baseline-Adjustment Models for Psychomotor Vigilance*

| Effect | *b* | 95% HDI | *pd* | ROPE % | $BF_{10}$ |
|---|---|---|---|---|---|
| **PVT median reaction time (ms)** | | | | | |
| *M0: No baseline adjustment* | | | | | |
| Light exposure | 0.67 | [−5.20, 6.61] | .586 | 100.0 | 0.15 |
| Physical activity | −25.63 | [−38.65, −12.89] | > .999 | 19.4 | 204.38 |
| *M1: Single visit-level baseline (reported model)* | | | | | |
| Light exposure | 1.70 | [−3.01, 6.32] | .766 | 100.0 | 0.16 |
| Physical activity | −2.67 | [−9.41, 3.92] | .784 | 100.0 | 0.23 |
| *M2: Decomposed baseline* | | | | | |
| Light exposure | 0.63 | [−3.63, 4.90] | .613 | 100.0 | 0.12 |
| Physical activity | 2.37 | [−4.14, 9.41] | .757 | 100.0 | 0.22 |
| *M3: Decomposed baseline and activity* | | | | | |
| Light exposure | −0.07 | [−4.16, 3.87] | .514 | 100.0 | 0.11 |
| Physical activity, within participant | 0.08 | [−17.58, 17.79] | .504 | 97.4 | 0.46 |
| Physical activity, between participants | 2.51 | [−4.65, 9.63] | .756 | — | 0.22 |
| **PVT mean response speed ($s^{-1}$)** | | | | | |
| *M0: No baseline adjustment* | | | | | |
| Light exposure | −0.013 | [−0.069, 0.046] | .669 | 94.1 | 0.17 |
| Physical activity | 0.274 | [0.149, 0.399] | > .999 | 0.1 | > 1000 |
| *M1: Single visit-level baseline (reported model)* | | | | | |
| Light exposure | −0.025 | [−0.069, 0.019] | .866 | 94.1 | 0.20 |
| Physical activity | 0.032 | [−0.034, 0.095] | .838 | 80.3 | 0.27 |
| *M2: Decomposed baseline* | | | | | |
| Light exposure | −0.014 | [−0.054, 0.026] | .758 | 98.6 | 0.14 |
| Physical activity | −0.016 | [−0.077, 0.047] | .691 | 91.5 | 0.18 |
| *M3: Decomposed baseline and activity* | | | | | |
| Light exposure | −0.008 | [−0.046, 0.030] | .654 | 99.6 | 0.11 |
| Physical activity, within participant | 0.026 | [−0.138, 0.193] | .623 | 50.6 | 0.46 |
| Physical activity, between participants | −0.018 | [−0.083, 0.047] | .708 | — | 0.19 |
| **PVT slowest-10% response speed ($s^{-1}$)** | | | | | |
| *M0: No baseline adjustment* | | | | | |
| Light exposure | −0.006 | [−0.054, 0.043] | .597 | 94.8 | 0.16 |
| Physical activity | 0.165 | [0.063, 0.272] | .999 | 1.6 | 24.61 |
| *M1: Single visit-level baseline (reported model)* | | | | | |
| Light exposure | −0.026 | [−0.066, 0.014] | .895 | 88.7 | 0.27 |
| Physical activity | 0.008 | [−0.045, 0.059] | .624 | 92.9 | 0.17 |

*Table S12 (continued)*

| Effect | $b$ | 95% HDI | *pd* | ROPE % | $BF_{10}$ |
|---|---|---|---|---|---|
| *M2: Decomposed baseline* | | | | | |
| Light exposure | −0.020 | [−0.057, 0.016] | .866 | 94.3 | 0.21 |
| Physical activity | −0.013 | [−0.065, 0.035] | .707 | 92.4 | 0.18 |
| *M3: Decomposed baseline and activity* | | | | | |
| Light exposure | −0.012 | [−0.047, 0.023] | .756 | 98.2 | 0.14 |
| Physical activity, within participant | 0.036 | [−0.119, 0.190] | .683 | 43.6 | 0.54 |
| Physical activity, between participants | −0.014 | [−0.066, 0.038] | .702 | — | 0.18 |

*Note.* M0 = no baseline adjustment; M1 = single standardised visit-level baseline (model reported in the main text and in Table S11); M2 = baseline decomposed into between- and within-participant components; M3 = M2 with physical activity additionally decomposed into a between-participant and a within-participant component (Method S8). All four models were fitted to the same complete-case observations. Analysis sample: $N$ = 807 assessments from 37 participants across 82 visits. In M3, the within-participant activity component replaces physical activity in all interaction terms; the between-participant component is an observational adjustment term, for which ROPE % is not shown. Values for response speed ($s^{-1}$) are given to three decimals. Values are posterior means ($b$) with 95% highest-density intervals (HDI) on the outcome scale. Light exposure (median photopic illuminance, $\log_{10}[lx + 1]$) and physical activity (median heart rate reserve) were $z$-standardised (Method S6). Their main-effect coefficients give the expected outcome difference per 1-*SD* increase at the mean of the other predictor; the light × activity coefficient gives the change in the light exposure coefficient per 1-*SD* increase in physical activity; assessment terms give the change per assessment. *pd* = probability of direction; ROPE % = percentage of the full posterior inside the outcome-specific region of practical equivalence (Table S3), shown for coefficients that are outcome differences (light exposure, physical activity, and light × activity) but not for intercepts, assessment terms (rates per assessment), adjustment terms, or variance parameters; $BF_{10}$ = Savage–Dickey Bayes factor against a point null at zero, computed with the prior of the respective coefficient; values above 1000 are shown as > 1000; SD = standard deviation; — = not applicable.

#### Results S6. Daytime Time Course

Figure S8 shows the daytime light exposure and physical activity coefficients; Tables S13–S14 report full model estimates and derived contrasts.

**Figure S8**

*Posterior Distributions of Light Exposure and Physical Activity Effects on Daytime Outcomes*

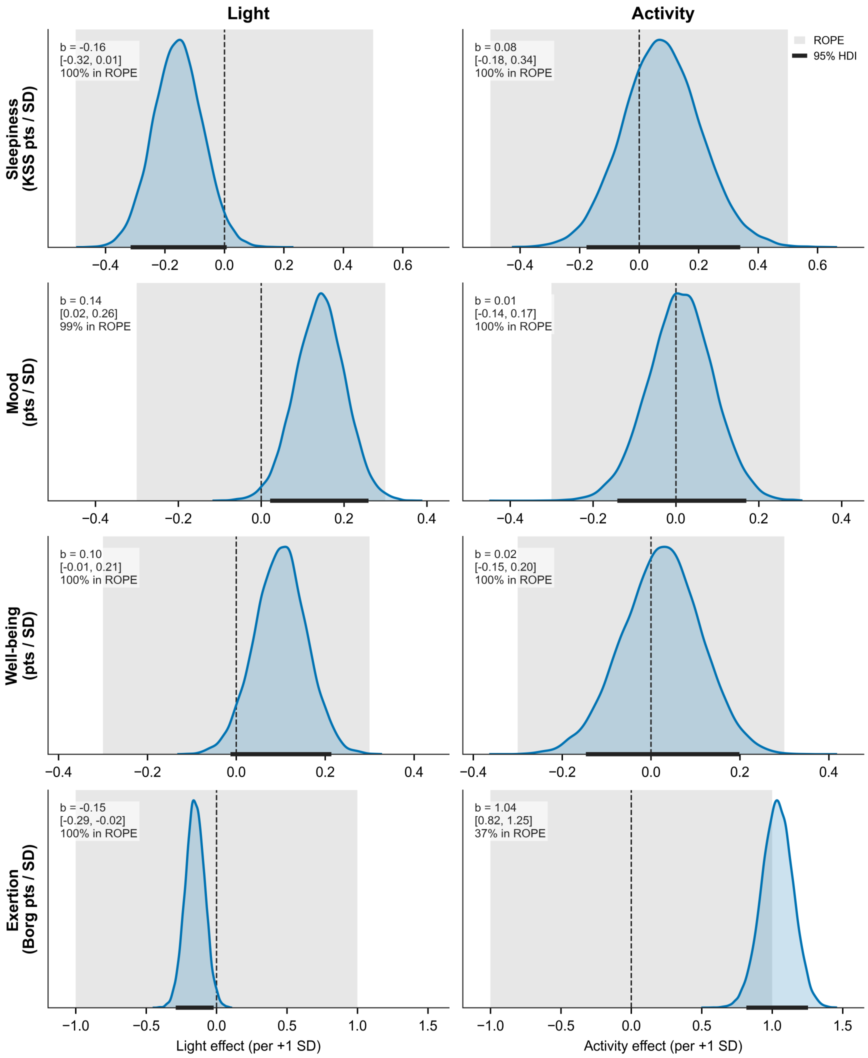

*Note.* Posterior densities of the light exposure (left) and physical activity (right) coefficients per 1-*SD* increase for sleepiness (KSS), mood, wellbeing, and perceived exertion (Borg) during the activity period. Curves show posterior densities; shaded grey bands indicate the outcome-specific region of practical equivalence (ROPE), black horizontal bars the 95% highest-density intervals (HDIs), and dashed vertical lines zero. Annotations report *b* [95% HDI] and the percentage of the posterior within the ROPE. Effects are expressed per 1-*SD* increase, with the other predictor at its mean. Estimates correspond to Table S13.

**Table S13**

*Posterior Estimates for Daytime Time-Course Outcomes*

| Effect | $b$ | 95% HDI | $pd$ | ROPE % | $BF_{10}$ |
|---|---|---|---|---|---|
| **Sleepiness (KSS)** | | | | | |
| *Fixed effects* | | | | | |
| Intercept | 2.76 | [2.49, 3.04] | — | — | — |
| Assessment (time course) | 0.09 | [0.06, 0.13] | > .999 | — | 147.12 |
| Light exposure | −0.16 | [−0.32, 0.01] | .970 | 100.0 | 1.09 |
| Physical activity | 0.08 | [−0.18, 0.34] | .721 | 99.9 | 0.34 |
| Light × activity | −0.06 | [−0.22, 0.10] | .756 | 100.0 | 0.24 |
| *Random-effect and residual parameters* | | | | | |
| Participant intercept SD | 0.72 | [0.48, 0.97] | — | — | — |
| Participant assessment-slope SD | 0.10 | [0.07, 0.13] | — | — | — |
| Visit intercept SD | 0.64 | [0.50, 0.80] | — | — | — |
| Residual SD | 0.85 | [0.82, 0.89] | — | — | — |
| **Mood** | | | | | |
| *Fixed effects* | | | | | |
| Intercept | 4.11 | [3.94, 4.28] | — | — | — |
| Assessment (time course) | −0.02 | [−0.03, 0.00] | .970 | — | 0.19 |
| Light exposure | 0.14 | [0.02, 0.26] | .989 | 99.5 | 3.32 |
| Physical activity | 0.01 | [−0.14, 0.17] | .565 | 100.0 | 0.30 |
| Light × activity | 0.09 | [−0.02, 0.21] | .943 | 100.0 | 0.81 |
| *Random-effect and residual parameters* | | | | | |
| Participant intercept SD | 0.36 | [0.17, 0.57] | — | — | — |
| Participant assessment-slope SD | 0.04 | [0.03, 0.06] | — | — | — |
| Visit intercept SD | 0.48 | [0.38, 0.59] | — | — | — |
| Residual SD | 0.50 | [0.48, 0.52] | — | — | — |
| **Wellbeing** | | | | | |
| *Fixed effects* | | | | | |
| Intercept | 4.07 | [3.88, 4.25] | — | — | — |
| Assessment (time course) | −0.02 | [−0.03, 0.00] | .977 | — | 0.24 |
| Light exposure | 0.10 | [−0.01, 0.21] | .956 | 100.0 | 0.92 |
| Physical activity | 0.02 | [−0.15, 0.20] | .615 | 99.9 | 0.35 |
| Light × activity | 0.06 | [−0.05, 0.17] | .848 | 100.0 | 0.37 |
| *Random-effect and residual parameters* | | | | | |
| Participant intercept SD | 0.46 | [0.28, 0.64] | — | — | — |
| Participant assessment-slope SD | 0.05 | [0.03, 0.06] | — | — | — |
| Visit intercept SD | 0.47 | [0.37, 0.58] | — | — | — |
| Residual SD | 0.42 | [0.40, 0.44] | — | — | — |
| **Perceived exertion (Borg)** | | | | | |
| *Fixed effects* | | | | | |
| Intercept | 2.24 | [2.00, 2.47] | — | — | — |

*Table S13 (continued)*

| Effect | $b$ | 95% HDI | $pd$ | ROPE % | $BF_{10}$ |
|---|---|---|---|---|---|
| Assessment (time course) | −0.04 | [−0.06, −0.01] | .997 | — | 0.75 |
| Light exposure | −0.15 | [−0.29, −0.02] | .986 | 100.0 | 1.17 |
| Physical activity | 1.04 | [0.82, 1.25] | > .999 | 37.1 | > 1000 |
| Light × activity | −0.10 | [−0.23, 0.03] | .928 | 100.0 | 0.30 |
| *Random-effect and residual parameters* | | | | | |
| Participant intercept SD | 0.60 | [0.40, 0.81] | — | — | — |
| Participant assessment-slope SD | 0.01 | [0.00, 0.03] | — | — | — |
| Visit intercept SD | 0.35 | [0.15, 0.55] | — | — | — |
| Residual SD | 1.47 | [1.40, 1.54] | — | — | — |

*Note.* Analysis samples: sleepiness, $N$ = 1,053 assessments from 37 participants across 88 visits; mood and wellbeing, $N$ = 1,040 assessments from 37 participants across 87 visits; perceived exertion, $N$ = 1,024 assessments from 37 participants across 87 visits. Assessment was centred at the midpoint of the twelve daytime assessments (30-min intervals) and entered additively, so the assessment term gives the change per assessment. Random-effect parameters are the *SD*s of the participant-level intercept and assessment slope and of the visit-level intercept. Values are posterior means ($b$) with 95% highest-density intervals (HDI) on the outcome scale. Light exposure (median photopic illuminance, $\log_{10}[\text{lx} + 1]$) and physical activity (median heart rate reserve) were $z$-standardised (Method S6). Their main-effect coefficients give the expected outcome difference per 1-*SD* increase at the mean of the other predictor; the light × activity coefficient gives the change in the light exposure coefficient per 1-*SD* increase in physical activity; assessment terms give the change per assessment. $pd$ = probability of direction; ROPE % = percentage of the full posterior inside the outcome-specific region of practical equivalence (Table S3), shown for coefficients that are outcome differences (light exposure, physical activity, and light × activity) but not for intercepts, assessment terms (rates per assessment), adjustment terms, or variance parameters; $BF_{10}$ = Savage–Dickey Bayes factor against a point null at zero, computed with the prior of the respective coefficient; values above 1000 are shown as > 1000; SD = standard deviation; — = not applicable. KSS = Karolinska Sleepiness Scale.

**Table S14**

*Derived Contrasts for Daytime Time-Course Outcomes*

| Contrast | $b$ | 95% HDI | $pd$ | ROPE % |
|---|---|---|---|---|
| **Sleepiness (KSS)** | | | | |
| Full-day change | 1.01 | [0.63, 1.42] | > .999 | 0.7 |
| Light exposure | −0.31 | [−0.63, 0.02] | .970 | 87.5 |
| Physical activity | 0.15 | [−0.35, 0.68] | .721 | 89.9 |
| Light exposure at high activity | −0.43 | [−0.97, 0.11] | .940 | 60.3 |
| Light exposure at low activity | −0.20 | [−0.56, 0.16] | .856 | 95.3 |
| Light × activity | −0.23 | [−0.89, 0.41] | .756 | — |
| **Mood** | | | | |
| Full-day change | −0.17 | [−0.34, 0.01] | .970 | 92.5 |
| Light exposure | 0.28 | [0.04, 0.52] | .989 | 54.8 |

*Table S14 (continued)*

| Contrast | *b* | 95% HDI | *pd* | ROPE % |
|---|---|---|---|---|
| Physical activity | 0.02 | [−0.28, 0.34] | .565 | 93.8 |
| Light exposure at high activity | 0.47 | [0.08, 0.86] | .990 | 18.9 |
| Light exposure at low activity | 0.10 | [−0.17, 0.35] | .765 | 93.7 |
| Light × activity | 0.38 | [−0.09, 0.85] | .943 | — |
| **Wellbeing** | | | | |
| Full-day change | −0.19 | [−0.37, −0.01] | .977 | 89.2 |
| Light exposure | 0.20 | [−0.03, 0.43] | .956 | 80.9 |
| Physical activity | 0.05 | [−0.29, 0.40] | .615 | 90.0 |
| Light exposure at high activity | 0.32 | [−0.05, 0.71] | .951 | 45.7 |
| Light exposure at low activity | 0.08 | [−0.17, 0.34] | .732 | 95.2 |
| Light × activity | 0.24 | [−0.22, 0.70] | .848 | — |
| **Perceived exertion (Borg)** | | | | |
| Full-day change | −0.40 | [−0.70, −0.12] | .997 | 100.0 |
| Light exposure | −0.31 | [−0.58, −0.04] | .986 | 100.0 |
| Physical activity | 2.07 | [1.63, 2.51] | > .999 | 0.0 |
| Light exposure at high activity | −0.50 | [−0.93, −0.04] | .987 | 98.3 |
| Light exposure at low activity | −0.11 | [−0.40, 0.20] | .758 | 100.0 |
| Light × activity | −0.40 | [−0.93, 0.14] | .928 | — |

*Note.* Full-day change is the predicted change from the first to the final daytime assessment at mean light exposure and physical activity. Values are posterior means (*b*) with 95% highest-density intervals (HDI) on the outcome scale. Rows labelled light exposure or physical activity compare high (+1 *SD*) with low (−1 *SD*) values of that predictor, that is, a 2-*SD* difference, with the other predictor at its mean. Rows labelled at high or low activity evaluate the light exposure contrast at +1 *SD* or −1 *SD* physical activity, and the light × activity row is the difference between these two rows (difference-in-differences). *pd* = probability of direction; ROPE % = percentage of the full posterior inside the outcome-specific region of practical equivalence (Table S3). Because the contrasts span 2 *SD*s, their ROPE percentages are not directly comparable with those of the per-*SD* coefficients. ROPE percentages are not reported for difference-in-differences or time-course moderation contrasts because no separate practical-equivalence threshold was prespecified for those estimands. Bayes factors are not reported for derived contrasts (Method S6): for contrasts that rescale a single coefficient, they equal the Bayes factor of that coefficient in the corresponding estimates table; for contrasts that combine several coefficients, they would depend on a prior that is implied by the coefficient priors rather than specified for the contrast.

**Results S7. Model Comparison**

Table S15 reports the PSIS-LOO comparisons of the candidate fixed-effect specifications. Among the comparisons judged reliable, there was no evidence that light exposure, physical activity, or their interaction improved out-of-sample prediction. In every comparison set with reliable PSIS-LOO estimates, all models predicted the data about as well as the best-ranked model, that is, no ΔELPD exceeded approximately twice its standard error. This applied to the change in DLMO in the primary analysis and in the Student-*t*, cumulative-light, and corrected-laboratory-light variants, as well as to daytime mood and perceived exertion, and it concurs with the inconclusive or practically negligible parameter estimates. The only difference that exceeded twice its standard error occurred for the change in melatonin AUC, where the light-only model outperformed the additive model; however, this comparison set contained observations with Pareto $\hat{k} > 0.7$ and was therefore not interpreted. For the same reason, the comparisons for the wider-prior and the age- and sex-adjusted DLMO variants, daytime sleepiness and wellbeing, and all baseline-adjusted vigilance outcomes were not interpreted (marked in the table). For the evening time-course outcomes, finally, the PSIS-LOO estimates of the reported models already contained such observations for every outcome. Hence, any comparison involving these models would be unreliable, and only their individual PSIS-LOO estimates are reported.

**Table S15**

*Leave-One-Out Cross-Validation Comparisons of Candidate Models*

| Model | ELPD (*SE*) | ΔELPD (*SE*) | $p_{LOO}$ | Weight | Max $\hat{k}$ | $\hat{k} > 0.7$ |
|---|---|---|---|---|---|---|
| **Change in DLMO (primary model)** | | | | | | |
| Light + activity | −396.3 (7.6) | — | 12.6 | .71 | 0.54 | 0 |
| Activity only | −397.0 (8.2) | 0.65 (1.57) | 10.8 | .23 | 0.67 | 0 |
| Full (light × activity) | −397.1 (7.5) | 0.75 (0.69) | 13.2 | .00 | 0.59 | 0 |
| Light only | −397.2 (7.7) | 0.84 (1.38) | 13.6 | .06 | 0.65 | 0 |
| Null (intercept only) | −397.9 (8.3) | 1.62 (2.21) | 11.6 | .00 | 0.59 | 0 |
| **Change in DLMO: Student-*t* likelihood** | | | | | | |
| Activity only | −395.5 (8.5) | — | 10.3 | .18 | 0.50 | 0 |
| Light + activity | −395.7 (8.0) | 0.25 (1.20) | 12.0 | .42 | 0.50 | 0 |
| Null (intercept only) | −395.9 (8.9) | 0.40 (1.35) | 10.4 | .39 | 0.55 | 0 |
| Light only | −396.3 (8.3) | 0.89 (1.46) | 12.4 | .00 | 0.52 | 0 |
| Full (light × activity) | −396.7 (7.8) | 1.22 (1.41) | 13.0 | .00 | 0.56 | 0 |
| **Change in DLMO: Wider coefficient priors[a]** | | | | | | |
| Light + activity | −396.5 (7.6) | — | 12.8 | .42 | 0.77 | 1 |
| Activity only | −396.9 (8.2) | 0.37 (1.60) | 10.9 | .35 | 0.56 | 0 |
| Light only | −397.1 (7.6) | 0.57 (1.53) | 13.5 | .23 | 0.56 | 0 |
| Full (light × activity) | −397.3 (7.4) | 0.76 (0.71) | 13.5 | .00 | 0.66 | 0 |
| Null (intercept only) | −397.9 (8.3) | 1.42 (2.33) | 11.6 | .00 | 0.59 | 0 |
| **Change in DLMO: Cumulative light exposure** | | | | | | |
| Light + activity | −395.8 (7.6) | — | 12.4 | .71 | 0.67 | 0 |
| Light only | −396.5 (7.6) | 0.70 (1.34) | 13.4 | .12 | 0.63 | 0 |
| Full (light × activity) | −396.7 (7.4) | 0.81 (0.67) | 13.2 | .00 | 0.67 | 0 |
| Activity only | −397.0 (8.2) | 1.12 (1.85) | 10.8 | .17 | 0.67 | 0 |
| Null (intercept only) | −397.9 (8.3) | 2.09 (2.43) | 11.6 | .00 | 0.59 | 0 |
| **Change in DLMO: Corrected laboratory light values** | | | | | | |
| Light + activity | −396.4 (7.6) | — | 12.5 | .69 | 0.69 | 0 |
| Activity only | −397.0 (8.2) | 0.58 (1.48) | 10.8 | .23 | 0.67 | 0 |
| Full (light × activity) | −397.1 (7.4) | 0.66 (0.69) | 13.2 | .00 | 0.61 | 0 |
| Light only | −397.3 (7.7) | 0.90 (1.44) | 13.5 | .08 | 0.65 | 0 |
| Null (intercept only) | −397.9 (8.3) | 1.55 (2.16) | 11.6 | .00 | 0.59 | 0 |
| **Change in DLMO: Adjusted for age and sex[a]** | | | | | | |
| Light + activity | −397.2 (7.8) | — | 13.6 | .56 | 0.67 | 0 |
| Activity only | −397.6 (8.4) | 0.41 (1.49) | 11.8 | .32 | 0.72 | 1 |
| Full (light × activity) | −397.8 (7.6) | 0.61 (0.82) | 13.9 | .00 | 0.68 | 0 |
| Light only | −398.1 (7.9) | 0.85 (1.48) | 14.3 | .12 | 0.63 | 0 |
| Null (intercept only) | −398.8 (8.5) | 1.59 (2.11) | 12.4 | .00 | 0.76 | 1 |
| **Change in melatonin AUC (exploratory)[a]** | | | | | | |
| Light only | 47.8 (10.6) | — | 8.6 | .95 | 0.79 | 2 |

*Table S15 (continued)*

| Model | ELPD (*SE*) | ΔELPD (*SE*) | $p_{\text{LOO}}$ | Weight | Max $\hat{k}$ | $\hat{k} > 0.7$ |
|---|---|---|---|---|---|---|
| Light + activity | 46.9 (10.5) | 0.89 (0.17) | 9.4 | .00 | 0.83 | 2 |
| Full (light × activity) | 46.1 (10.2) | 1.70 (0.86) | 10.3 | .00 | 0.76 | 2 |
| Null (intercept only) | 45.7 (11.3) | 2.06 (2.11) | 6.9 | .05 | 0.68 | 0 |
| Activity only | 44.6 (11.3) | 3.19 (2.16) | 8.1 | .00 | 0.86 | 1 |
| **Daytime: Sleepiness (KSS)[a]** | | | | | | |
| Assessment + light + activity | −1390.4 (42.2) | — | 113.3 | > .99 | 0.96 | 1 |
| Full (+ light × activity) | −1390.7 (42.1) | 0.35 (0.41) | 113.7 | .00 | 0.93 | 2 |
| Assessment only | −1391.2 (42.3) | 0.79 (0.63) | 114.2 | .00 | 0.92 | 2 |
| Null (intercept only) | −1393.0 (42.6) | 2.58 (2.12) | 117.3 | .00 | 0.81 | 3 |
| **Daytime: Mood** | | | | | | |
| Assessment + light + activity | −809.5 (33.3) | — | 107.5 | .00 | 0.66 | 0 |
| Assessment only | −809.5 (33.5) | 0.00 (0.60) | 107.8 | .14 | 0.69 | 0 |
| Full (+ light × activity) | −809.5 (33.3) | 0.02 (0.56) | 107.7 | .46 | 0.69 | 0 |
| Null (intercept only) | −809.7 (33.3) | 0.15 (1.34) | 108.3 | .40 | 0.64 | 0 |
| **Daytime: Wellbeing[a]** | | | | | | |
| Assessment only | −633.7 (31.0) | — | 113.8 | .95 | 0.80 | 1 |
| Assessment + light + activity | −634.0 (30.9) | 0.34 (0.41) | 114.1 | .00 | 0.80 | 1 |
| Null (intercept only) | −634.2 (31.0) | 0.59 (1.14) | 114.6 | .05 | 0.73 | 1 |
| Full (+ light × activity) | −634.3 (30.9) | 0.69 (0.56) | 114.4 | .00 | 0.82 | 1 |
| **Daytime: Perceived exertion (Borg)** | | | | | | |
| Assessment + light + activity | −1877.6 (38.9) | — | 60.1 | .29 | 0.44 | 0 |
| Full (+ light × activity) | −1877.9 (38.9) | 0.32 (1.42) | 60.5 | .38 | 0.52 | 0 |
| Assessment only | −1880.6 (39.1) | 3.00 (4.32) | 65.9 | .18 | 0.57 | 0 |
| Null (intercept only) | −1883.3 (38.4) | 5.71 (5.20) | 65.0 | .15 | 0.47 | 0 |
| **Baseline-adjusted evening: PVT median reaction time (ms)[a]** | | | | | | |
| M3 | −3485.5 (37.6) | — | 132.1 | .82 | 1.07 | 6 |
| M2 | −3486.6 (37.6) | 1.03 (1.41) | 134.3 | .00 | 1.07 | 5 |
| M1 | −3487.2 (37.7) | 1.62 (1.83) | 134.4 | .00 | 0.93 | 6 |
| M0 | −3487.9 (37.4) | 2.33 (2.72) | 136.4 | .18 | 1.02 | 6 |
| **Baseline-adjusted evening: PVT mean response speed ($s^{-1}$)[a]** | | | | | | |
| M2 | 339.5 (42.5) | — | 135.4 | .18 | 1.47 | 6 |
| M3 | 339.5 (42.9) | 0.05 (1.62) | 135.2 | .56 | 1.57 | 4 |
| M1 | 338.9 (42.3) | 0.58 (1.25) | 135.6 | .02 | 1.46 | 3 |
| M0 | 338.0 (41.7) | 1.48 (2.33) | 138.4 | .24 | 1.50 | 3 |
| **Baseline-adjusted evening: PVT slowest-10% response speed ($s^{-1}$)[a]** | | | | | | |
| M2 | 157.7 (32.9) | — | 116.9 | .40 | 0.82 | 2 |
| M3 | 157.4 (32.9) | 0.30 (1.38) | 116.6 | .37 | 0.78 | 3 |
| M1 | 156.2 (32.9) | 1.46 (1.51) | 118.3 | .00 | 0.80 | 2 |
| M0 | 154.8 (32.5) | 2.86 (3.30) | 121.4 | .22 | 0.75 | 2 |

*Table S15 (continued)*

| Model | ELPD (*SE*) | ΔELPD (*SE*) | $p_{\text{LOO}}$ | Weight | Max $\hat{k}$ | $\hat{k} > 0.7$ |
|---|---|---|---|---|---|---|
| **Evening time course: reported full models (no comparison)**[a] | | | | | | |
| Sleepiness | −925.4 (36.6) | — | 149.3 | — | 0.86 | 6 |
| Mood | −458.6 (45.0) | — | 140.4 | — | 1.64 | 2 |
| Wellbeing | −361.4 (40.3) | — | 144.7 | — | 1.38 | 4 |
| PVT median RT | −3571.9 (37.6) | — | 140.0 | — | 1.03 | 5 |
| PVT mean response speed | 345.3 (40.4) | — | 139.5 | — | 1.31 | 5 |
| PVT slowest-10% speed | 159.3 (32.5) | — | 124.2 | — | 0.73 | 1 |

*Note.* Models are listed by predictive accuracy within each comparison set. ELPD = expected log pointwise predictive density estimated with Pareto-smoothed importance-sampling leave-one-out cross-validation (PSIS-LOO); ΔELPD = ELPD of the best-ranked model minus ELPD of the comparison model (positive = worse predictive accuracy than the best-ranked model); $p_{\text{LOO}}$ = effective number of parameters; Weight = stacking weight; $\hat{k}$ = Pareto shape diagnostic. A model was considered predictively superior only if ΔELPD exceeded approximately 2 *SE*. For the daytime outcomes, all candidate models retained the random intercepts and assessment slopes; the null model contained no fixed assessment term. For the evening time course, PSIS-LOO was computed for the reported full models only. [a] At least one model of the set had observations with $\hat{k} > 0.7$; the comparison is unreliable and was not interpreted.

#### Results S8. Analysis Samples, Sampling Diagnostics, and Posterior Predictive Check

All reported models met the convergence criteria described in Method S6 (Table S16). Because complete-case deletion was applied per model, the analysis samples differ across outcomes. Figure S9 shows the posterior predictive check of the primary model for the change in DLMO. The simulated data sets reproduced the location and overall range of the observed values, but the observed distribution was more sharply peaked around zero than the simulated distributions, a pattern consistent with heavier-than-normal tails. This motivated the Student-*t* sensitivity analysis (Method S7), which led to the same conclusions as the primary model (Results S2).

**Table S16**

*Analysis Samples and Sampling Diagnostics of All Reported Models*

| Outcome / model | $N_{obs}$ | $n_{part}$ | $n_{vis}$ | Max $\hat{R}$ | Min $ESS_{bulk}$ | Min $ESS_{tail}$ | Div. | Min BFMI |
|---|---|---|---|---|---|---|---|---|
| **Change in DLMO** | | | | | | | | |
| Primary model | 81 | 36 | — | 1.001 | 5,161 | 6,459 | 0 | 0.64 |
| Student-*t* likelihood | 81 | 36 | — | 1.001 | 5,370 | 8,873 | 0 | 0.67 |
| Wider coefficient priors | 81 | 36 | — | 1.001 | 4,819 | 7,887 | 0 | 0.66 |
| Cumulative light exposure | 81 | 36 | — | 1.001 | 5,227 | 6,937 | 0 | 0.65 |
| Corrected laboratory light values | 81 | 36 | — | 1.001 | 5,352 | 8,570 | 0 | 0.65 |
| Adjusted for age and sex | 81 | 36 | — | 1.002 | 4,626 | 8,230 | 0 | 0.67 |
| **Change in melatonin AUC (exploratory)** | | | | | | | | |
| Primary model | 87 | 37 | — | 1.001 | 6,745 | 7,710 | 0 | 0.85 |
| **Evening time course** | | | | | | | | |
| Sleepiness | 827 | 37 | 84 | 1.001 | 4,133 | 4,316 | 0 | 0.69 |
| Mood | 827 | 37 | 84 | 1.001 | 3,635 | 4,106 | 0 | 0.69 |
| Wellbeing | 827 | 37 | 84 | 1.001 | 5,973 | 4,960 | 0 | 0.70 |
| PVT median RT | 827 | 37 | 84 | 1.001 | 3,229 | 6,998 | 0 | 0.69 |
| PVT mean response speed | 827 | 37 | 84 | 1.002 | 2,762 | 3,773 | 0 | 0.71 |
| PVT slowest-10% speed | 827 | 37 | 84 | 1.001 | 3,540 | 4,217 | 0 | 0.70 |
| **Evening time course, baseline-adjusted** | | | | | | | | |
| PVT median RT, M0 | 807 | 37 | 82 | 1.001 | 3,673 | 9,402 | 0 | 0.70 |
| PVT median RT, M1 | 807 | 37 | 82 | 1.006 | 1,165 | 4,343 | 0 | 0.68 |
| PVT median RT, M2 | 807 | 37 | 82 | 1.003 | 1,676 | 2,783 | 0 | 0.67 |
| PVT median RT, M3 | 807 | 37 | 82 | 1.001 | 1,953 | 4,088 | 0 | 0.69 |
| PVT mean response speed, M0 | 807 | 37 | 82 | 1.001 | 3,234 | 4,313 | 0 | 0.70 |
| PVT mean response speed, M1 | 807 | 37 | 82 | 1.005 | 1,111 | 3,535 | 0 | 0.68 |
| PVT mean response speed, M2 | 807 | 37 | 82 | 1.005 | 1,752 | 4,557 | 0 | 0.68 |
| PVT mean response speed, M3 | 807 | 37 | 82 | 1.001 | 1,994 | 4,965 | 0 | 0.68 |
| PVT slowest-10% speed, M0 | 807 | 37 | 82 | 1.001 | 3,002 | 3,495 | 0 | 0.67 |
| PVT slowest-10% speed, M1 | 807 | 37 | 82 | 1.002 | 1,344 | 3,480 | 0 | 0.66 |
| PVT slowest-10% speed, M2 | 807 | 37 | 82 | 1.002 | 1,599 | 3,761 | 0 | 0.68 |
| PVT slowest-10% speed, M3 | 807 | 37 | 82 | 1.002 | 1,651 | 3,811 | 0 | 0.67 |
| **Daytime time course** | | | | | | | | |
| Sleepiness | 1,053 | 37 | 88 | 1.001 | 8,473 | 11,513 | 0 | 0.72 |
| Mood | 1,040 | 37 | 87 | 1.001 | 2,628 | 1,939 | 0 | 0.69 |
| Wellbeing | 1,040 | 37 | 87 | 1.001 | 8,215 | 9,128 | 0 | 0.71 |
| Perceived exertion | 1,024 | 37 | 87 | 1.001 | 4,809 | 4,885 | 0 | 0.75 |
| **Change in DLMO: categorical model** | | | | | | | | |
| Light condition × activity group | 110 | 41 | — | 1.001 | 5,522 | — | 0 | — |

*Note.* $N_{obs}$ = observations entering the model (visits for the change in DLMO and melatonin AUC; assessments for time-course outcomes); $n_{part}$ = participants; $n_{vis}$ = visits (— where observations are visits). $\hat{R}$ = rank-normalised split-$\hat{R}$; ESS = effective sample size; Div. = divergent transitions; BFMI = Bayesian fraction of missing information. Extremes were taken over all posterior variables, including latent random effects. For the categorical model, the saved notebook output provided the maximum $\hat{R}$, minimum bulk ESS, and number of divergences; tail ESS and BFMI were not included in that output (—). All models were sampled with 4 chains of 6,000 post-warm-up draws.

**Figure S9**

*Posterior Predictive Check of the Primary Model for the Change in DLMO*

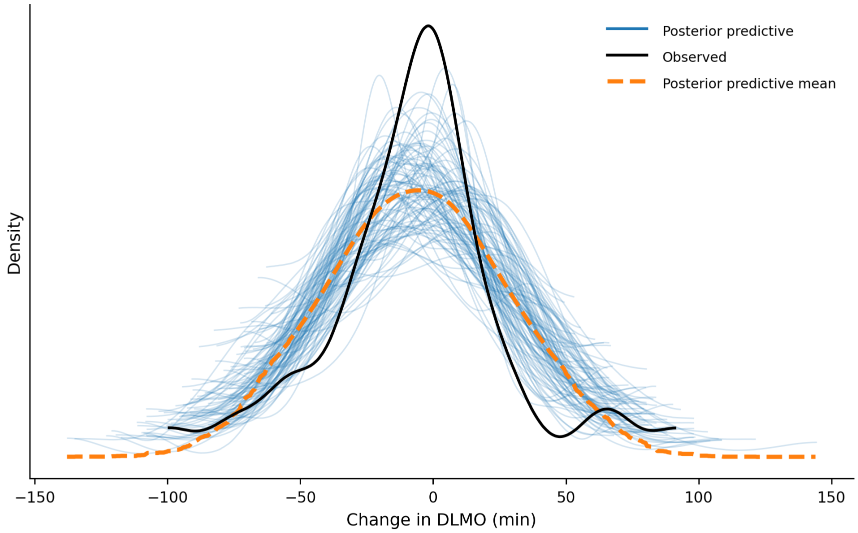

*Note.* The black line shows the kernel density of the observed change in DLMO (Evening-2 minus Evening-1, min), thin blue lines show the densities of 100 data sets simulated from the posterior predictive distribution of the primary model (Gaussian likelihood; Table S5), and the dashed orange line shows their mean.
